# Genetic parallelism underpins convergent mimicry coloration across Lepidoptera

**DOI:** 10.1101/2025.06.26.661542

**Authors:** Yacine Ben Chehida, Eva S.M. van der Heijden, Edward J. Page, Patricio A. Salazar C., Neil Rosser, Kimberly Gabriela Gavilanes Córdova, Mónica Sánchez-Prado, María José Sánchez-Carvajal, Franz Chandi, Alex P Arias-Cruz, Maya Radford, Gerardo Lamas, Chris Jiggins, James Mallet, Melanie McClure, Camilo Salazar, Marianne Elias, Caroline N. Bacquet, Nicola J. Nadeau, Kanchon K. Dasmahapatra, Joana I. Meier

## Abstract

Convergent evolution, the repeated evolution of similar phenotypes in response to the same selective pressures across multiple lineages, is widespread in nature. The extent to which the same genetic mechanisms contribute to convergent evolution could reveal whether the pathway towards these optimal endpoints is flexible or constrained to follow a particular route. Although mimicry of aposematic colour patterns is well known in Lepidoptera, our knowledge of the genetic basis of these convergent patterns is mostly restricted to a few closely-related species. Here we study the genetic basis of mimicry across seven species of Ithomiini and *Heliconius* butterflies and a day-flying *Chetone* moth, representing lineages that diverged between ∼1-120 Mya, each presenting similar colour pattern switches. In all the butterfly species, the genetic variants most strongly associated with convergent colour pattern switches are similarly located in non-coding regions near the genes *ivory* and *optix*. Colour pattern variation in the moth is associated with a ∼1 Mb inversion around *ivory* paralleling the supergene architecture of the co-mimic *Heliconius numata*. In contrast to previous studies in *Heliconius*, there is limited evidence of alleles shared by means of hybridization in convergence among closely-related ithomiine species. Repeated parallel evolution of regulatory switches via reuse of the same two genes suggests that convergent colour pattern evolution is highly constrained, even across large evolutionary timescales.

## Main

Convergent or parallel evolution is a natural experiment where unrelated species independently evolve similar traits in response to similar selective pressures. It informs us about the extent to which evolution is repeatable and thus predictable^1–4^. Highly divergent lineages can show strong trait convergence, for example associated with the repeated colonisation of land, water or air^5^ or the repeated evolution of resistance to challenges like drugs, insecticides or drought^6,7^.

Trait convergence in different species can be caused by genetic changes at different loci or the same loci (“gene reuse”). Gene reuse is predicted to be more common among closely related lineages or when developmental pathways towards shared fitness optima are constrained^8,9^. Where genes are reused, convergence may result from independent evolution of the same phenotype, or because the same alleles are reused (“allele sharing”), either from ancestral standing variation^10^, or as a result of introgression among hybridizing species^9,11^. Allele sharing is expected mainly among very closely related species^12,13^. Examples demonstrating convergent evolution through each of these mechanisms are known, but few systems exist where the same phenotypes have evolved multiple times in highly replicated responses to the same selective pressures. Here we employ such a system over a large range of divergence times spanning ∼1-120 My.

Mimicry rings are spectacular examples of convergent evolution in which multiple, often unrelated, sympatric taxa converge on the same phenotype usually in the context of aposematic signalling^14–17^. The genetics of convergence in mimicry rings has been studied mainly in butterflies (but see^18,19^) where changes in the regulatory regions of a limited number of genes typically underpin phenotypic changes. In *Papilio* butterflies, female-limited Batesian mimicry (where palatable species mimic toxic species) is controlled by reuse of the gene *doublesex*^20^. In *Heliconius* butterflies, where the mimicry is Müllerian (all the species involved are unpalatable and share the cost of educating predators), convergent black/red/yellow patterning between co-mimetic subspecies of *Heliconius erato* and *Heliconius melpomene* (∼10 MY divergent; Figure 1) results from the reuse of the *optix*, *ivory* and *WntA* genes via independent mutations at regulatory regions^21,22^. Mimicry among more closely related *Heliconius* species often results from allele sharing via introgression at these genes^23–25^. Structural variants such as inversions can maintain tightly linked groups of genes, preventing recombination and the production of non-mimetic phenotypes, as demonstrated in *Heliconius numata* where colour pattern differences between distinct mimetic morphs are controlled by multiple overlapping inversions containing *ivory*^26^.

**Fig. 1.**
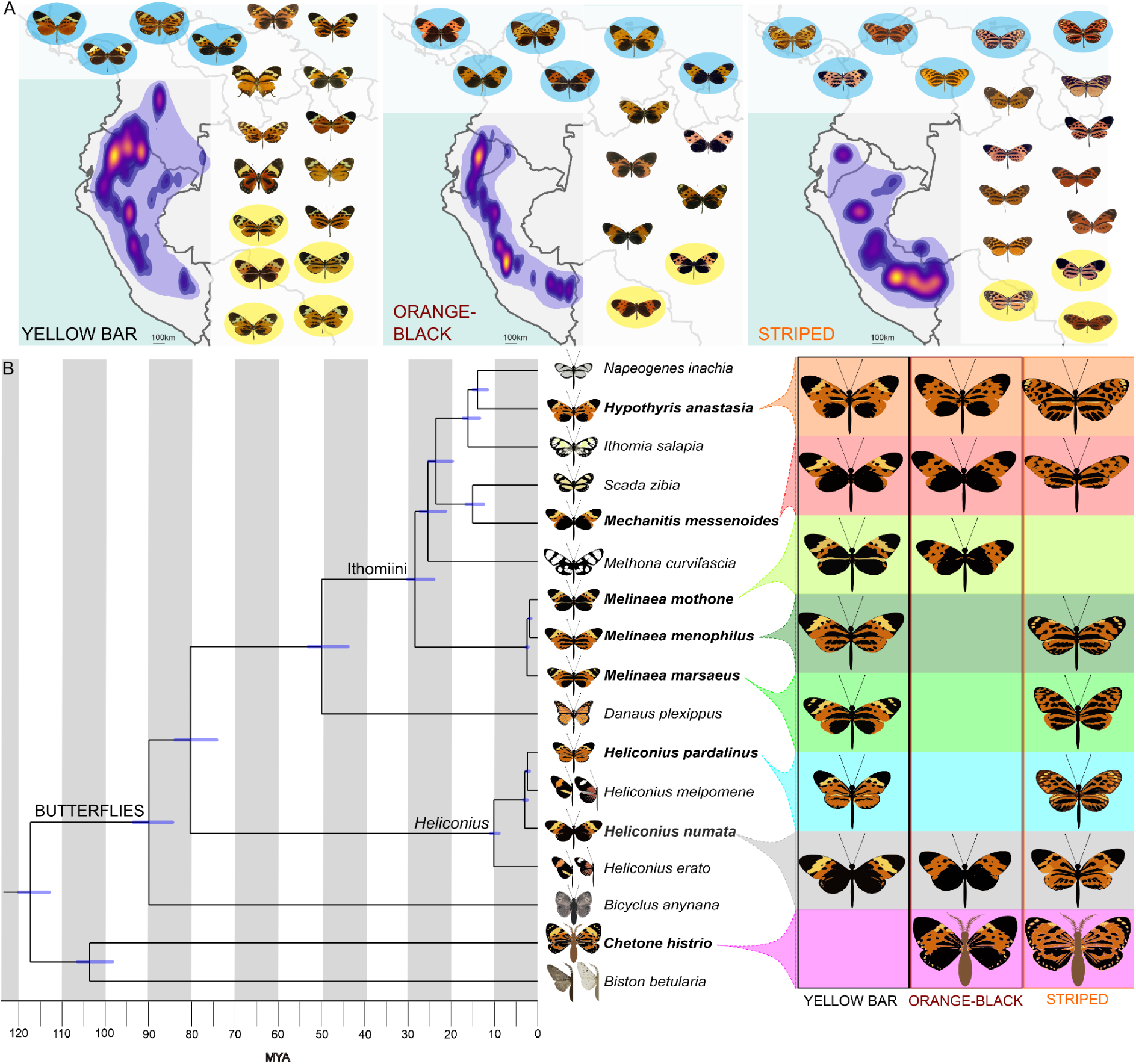
Phylogenetic relationships and mimetic phenotypes. a, Geographic distributions of the three tiger sub-mimicry rings: yellow-bar, orange/black and striped, based on Doré et al. 2021^27^ together with representative mimicry ring member taxa (detailed in Extended Data Fig. 1). The purple outline shows the maximum range of each sub-mimicry ring with the approximate abundance of taxa indicated via heatmap colouration. Taxa with a blue background were investigated using GWA/QTL. Taxa used in gene flow analyses are shown on a yellow background. **b,** Time-calibrated phylogeny including in bold the eight lepidopteran species investigated in detail. Blue bars represent 95% confidence intervals of the node ages. Large images depict the phenotypic variation of the species investigated in this study. Smaller images show other representative non-mimetic taxa.

The neotropical “tiger” mimicry ring to which *Hel. numata* and some other heliconiine species belong is exceptionally species-rich, including over 100 species from five lepidopteran families (Figure 1, Extended Data Fig. 1). It is dominated by many chemically-defended ithomiine species, and also includes day-flying moths among other taxa (Fig. 1; ^27,28^). Many of the species exhibit colour-pattern variation, where two or more subspecies are members of different sub-mimicry rings (orange/black, yellow-bar or striped) (Figure 1). Here we use this replicated natural experiment in convergent evolution, where mimetic lineages have diverged between ∼1-120 million years ago, to i) test how divergence time shapes gene reuse during repeated adaptation^9^ and; ii) where the same genes are reused, to test the contribution of introgressive allele sharing. Specifically, we use within-species genome-wide association (GWA) analyses to understand the genetic architecture of two mimetic phenotypes, the presence/absence of the forewing yellow bar and the extent of hindwing melanisation, in seven species from five genera (Fig. 1).

### Forewing yellow bar in Ithomiini butterflies: repeated reuse of *ivory*

Using whole-genome sequences of 285 wild-caught individuals, GWA was used to find genotypic associations with the presence or absence of the forewing yellow bar in the ithomiine species *Melinaea mothone* (49 specimens), *Melinaea menophilus* (64 specimens), *Mechanitis messenoides* (111 specimens) and *Hypothyris anastasia* (61 specimens). In all comparisons, clusters of significantly associated SNPs were identified in the long non-coding RNA *ivory* (near the gene *cortex*) which controls melanisation patterns across Lepidoptera (Fig. 2, Extended Data Fig. 2; ^29–34)^. In all cases, alleles associated with the presence of the yellow bar are recessive (Extended Data Figs. 2-6). Apart from *Melinaea menophilus* where the peak of association is relatively broad, for each species a small number of SNPs in narrow 1155-2140 bp genomic intervals are perfectly associated with the phenotype (Fig. 2 and Extended Data Figs. 2-6). We find surprising concordance in the location of the genomic intervals controlling an identical mimetic phenotype across these four species (Figs. 2 and 3). While there is no clear sequence homology in the identified regions, in all four species they lie within the first intron of *ivory*, 25,800-33,500 bp downstream of the *ivory* promoter and a short distance upstream of the E230 *cis*-regulatory element^32^. Tian et al. 2024^33^ demonstrated that the microRNA *mir-193* derived from *ivory* is likely the main effector gene, repressing multiple pigmentation genes. The concordant GWA peaks may indicate the existence of conserved transcriptional control of *ivory* expression unchanged over ∼28 million years of evolution in the Ithomiini.

**Fig. 2.**
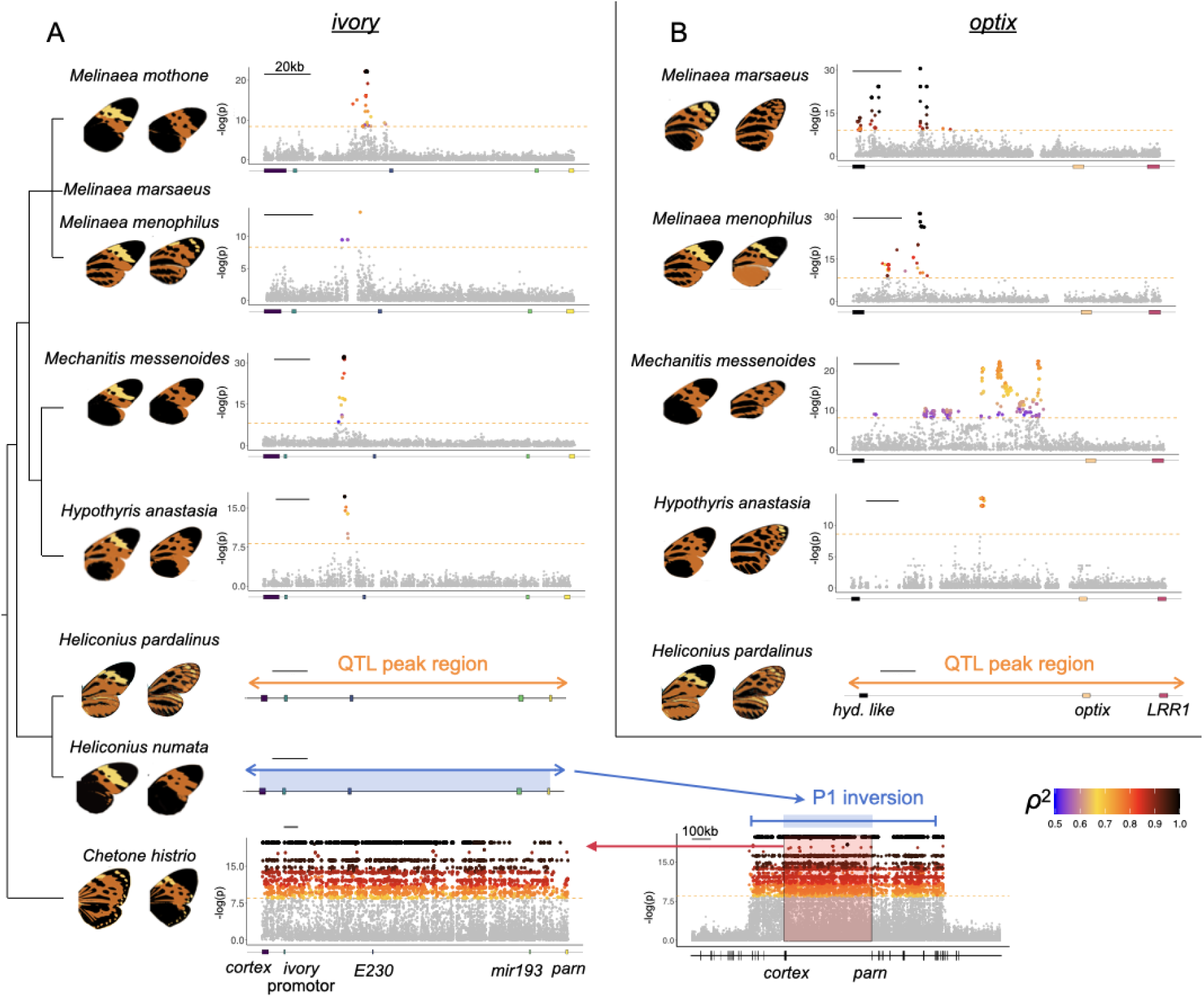
***Ivory* and *optix* control convergent phenotypes across multiple species.** Zoomed Manhattan plots of genome-wide association for wing pattern variation involving **a,** *ivory* and **b,** *optix*. SNPs above the Bonferroni-corrected significance threshold (orange dashed line) are coloured according to the strength of correlation (ρ^2^, squared Spearman’s rank correlation coefficient) between genotype and the phenotypes compared as shown in the wing images. Black points represent SNPs fully associated with the phenotypes. The bottom right Manhattan plot shows the wider ∼1 Mb region of high association in *Chetone histrio* around the *ivory* region. The results for *Heliconius numata* were retrieved from Jay et al. 2022. *Heliconius pardalinus* results are based on QTL mapping (Extended Data Fig. 16). E230: *cis*-regulatory element 230^32^; *hyd. like*: hydrolyse like; *LRR1*: Leucine Rich Repeat Protein 1. Association plots across the whole genome are shown in Extended Data Fig. 2. The ∼400 kb P1 inversion is associated with colour pattern variation in *Heliconius numata*^26^ and corresponds closely in location to the *Chetone histrio* inversion as shown by the blue arrow (Extended Data Fig. 24).

**Fig. 3.**
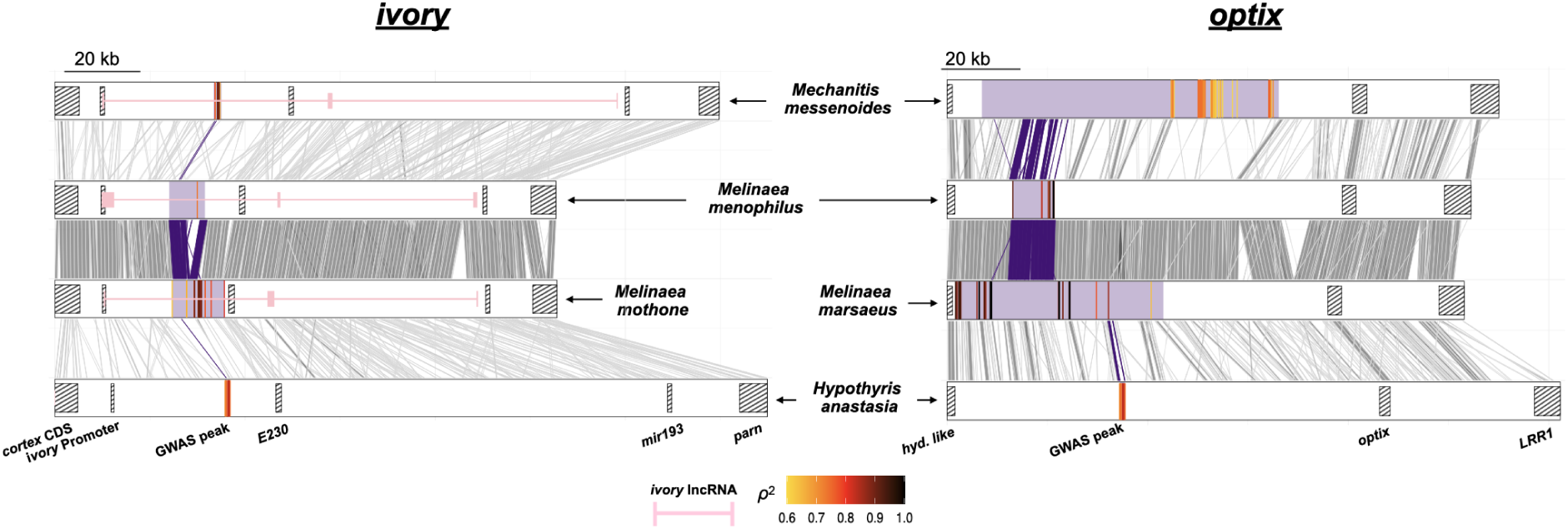
Concordant locations of SNPs associated with phenotypes at *ivory* and *optix*. Purple-shaded blocks represent intervals containing SNPs significantly associated with wing phenotypes (Fig. 1). Homologous regions (1,000 bp segments) are shown with grey lines between species pairs. Purple lines connect regions of homology within GWA intervals that are shared across adjacent species pairs, highlighting conserved areas. Within each species, the SNPs most strongly associated with the phenotype are coloured according to the strength of correlation with the phenotype (ρ^2^, squared Spearman’s rank correlation coefficient). Partial *ivory* annotations are shown in pink for three of the species. Hatched boxes mark annotated genomic features: E230: *cis*-regulatory element 230^32^; *hyd. like*: hydrolyse like; *LRR1*: Leucine Rich Repeat Protein 1.

### Hindwing melanisation in Ithomiini butterflies: repeated reuse of *optix*

We next used GWA to uncover genotypic associations with variation in orange/black hindwing patterning in *Melinaea marsaeus*, *Melinaea menophilus*, *Mechanitis messenoides* and *Hypothyris anastasia*. In all comparisons, we found peaks of association upstream of the known colour patterning gene *optix*, with genotype-phenotype correlations ranging from 0.54 to 1 (Fig. 2). In most cases these were usually the SNPs with the strongest associations across the genomes (Extended Data Fig. 2, 7-9). Additional associated SNPs in some species are likely a result of population structure correlated with the phenotypes. The regions of peak association in the two *Melinaea* species correspond closely, and they are near but not overlapping the region identified in *Hyp. anastasia*. All three genomic intervals fall within the wider associated region identified in *Mechanitis messenoides* (Fig. 3).

In *Mechanitis messenoides* we additionally investigated genetic associations with orange/black patterning in the forewing base and tip, which also yielded associated SNPs near *optix*. The SNPs showing the strongest genotype-phenotype correlations for the orange/black patterns in different wing regions fall in three separate clusters (Fig. 4, Extended Data Fig. 10-13). These may correspond to separate *cis*-regulatory modules of *optix* controlling different aspects of wing melanisation, similar to those proposed for different red/black phenotypes in *Heliconius*^25,35^. In contrast, orange/black patterning in the forewing tip of *Melinaea menophilus* was strongly associated (genotype-phenotype correlations of 0.87) with SNPs located between the *Hox* genes *Antennapedia* (*Antp*) and *Ultrabithorax* (*Ubx*) (Extended Data Fig. 14-15). As *Ubx* expression in butterflies is restricted to the hindwing, this phenotype likely arises through modulation of *Antp*^36^. This is similar to *Bicyclus anynana*, where *Antp* and *Ubx* promote the eyespot development in fore- and hindwing, respectively^37^.

**Fig. 4.**
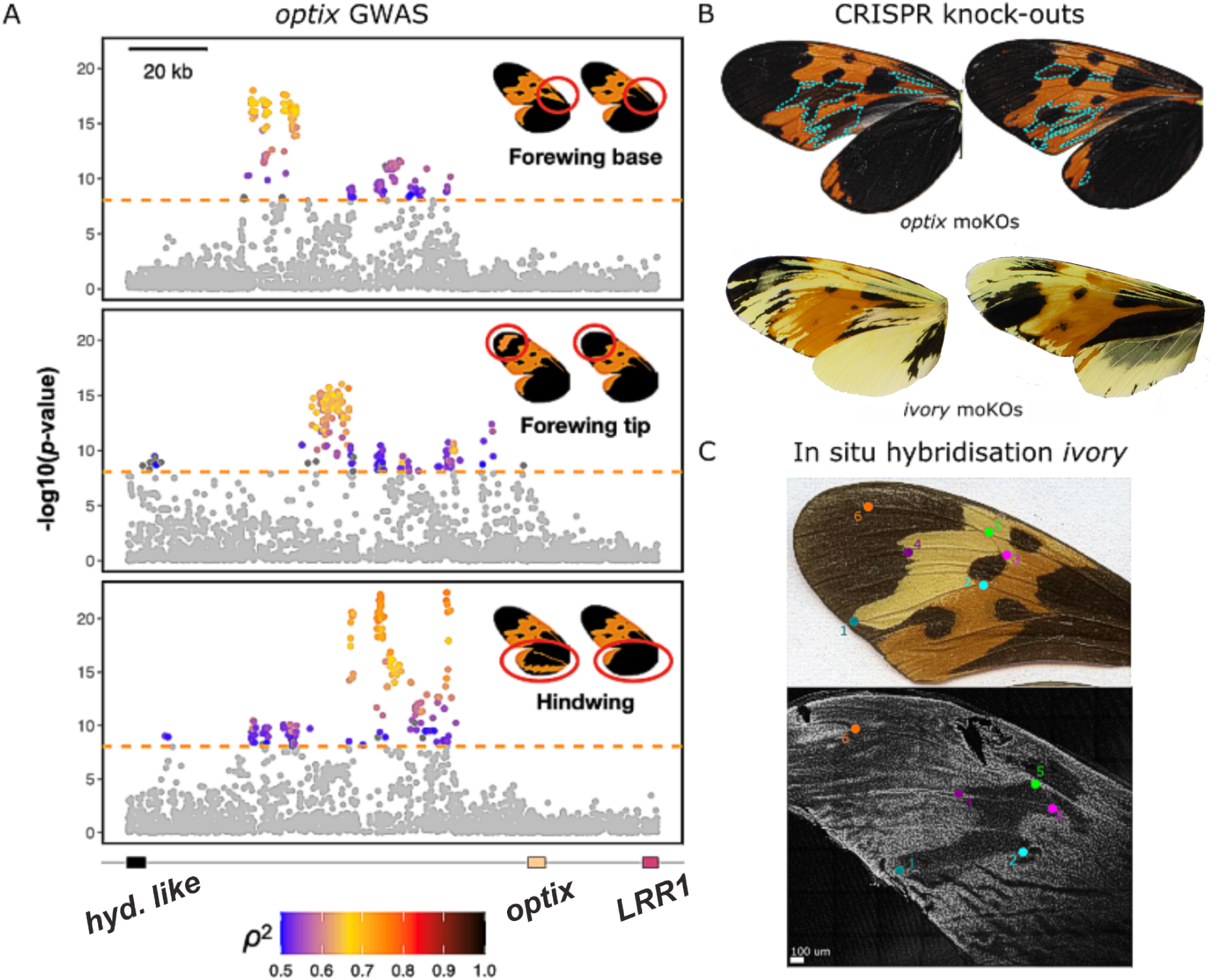
**Detailed phenotypic and functional analysis in *Mechanitis messenoides***. a, Genome-wide association analysis for separate forewing base, forewing tip and hindwing black/orange phenotypes (red-circled regions on wings) indicate three distinct *cis-*regulatory modules of *optix.* SNPs above the significance threshold (dashed orange line) are coloured according to the strength of association between genotype and phenotype (squared Spearman’s rank correlation coefficient, ρ²). *hyd. like* (hydrolase-like) and *LRR1* (leucine-rich repeat protein 1). Genome-wide association plots are shown in Extended Data Figure 10. **b,** CRISPR mutagenesis of *optix* and *ivory in Mec. messenoides.* Wings of mosaic knockout individuals are shown: orange scales turn black in *optix* mutants (blue dashed lines highlighting mutant patches), orange and black scales turn yellow in *ivory* mutants. Additional mutants are shown in Extended Data Figure 33. **c,** HCR *in-situ* hybridisation for *ivory* in *Mec. messenoides messenoides*. Top shows an adult forewing, and the bottom, a day 3 pupal forewing stained for *ivory*. The coloured dots indicate vein-based wing landmarks demarcating the yellow bar region which lacks *ivory* expression. Extended Data Figure 34 shows *Mec. messenoides deceptus*.

### Genetic architecture of colour patterning in *Heliconius pardalinus*

Loci controlling colour patterning in *Hel. pardalinus* were investigated by mapping variation segregating in 82 backcross individuals between subspecies^38^. The only significant quantitative trait locus (QTL) for forewing yellow patterning is on chromosome 15 and contains *ivory* (Fig. 2, Extended Data Fig. 16). A QTL for forewing orange patterning is found on chromosome 18, and QTLs for hindwing orange patterning are found on chromosomes 13^39^ and 18 (Fig. 2, Extended Data Fig. 16). Both the chromosome 18 QTLs encompass *optix*. These results are consistent with our findings from Ithomiini, showing that both *ivory* and *optix* predictably control convergent phenotypes across 80 My of divergence time.

### Genetic architecture of colour patterning in the moth *Chetone histrio*

Unlike the other species in which different mimetic forms within species have broadly parapatric distributions (Extended Data Figs. 17-22), the moth *Chetone histrio* is locally polymorphic in Peru with individuals belonging to either the striped (*C. histrio histrio*) or orange/black (*C. histrio hydra*) sub-mimicry rings which differ in multiple colour pattern elements across both fore- and hindwings (Extended Data Figure 23). Here it mimics, among other taxa, the similarly locally-polymorphic species *Heliconius numata*. Unlike in the previous GWA results, the comparison between genomes of Peruvian *C. histrio histrio* (20 individuals) and *C. histrio hydra* (15 individuals) shows a ∼1 Mb block of SNPs in perfect association with the phenotype (Fig. 2). This genomic interval includes *ivory* and several other flanking genes. Examination of Illumina read-pair orientation and mapped insert sizes demonstrates that this block corresponds to a 1.018 Mb inversion (Extended Data Fig. 24). All *C. histrio histrio* are homozygous for one inversion-type, and the *C. histrio hydra are* either heterozygous or homozygous for the other inversion-type. Colour pattern polymorphism in *Hel. numata* is also maintained via an inversion architecture, and in both species the inversions likely maintain allelic combinations (“supergenes”) that simultaneously control multiple colour pattern elements^26^. One of the breakpoints of the *Hel. numata* P1 inversion is similarly located to that in *C. histrio*, falling between the sucrose-6-phosphate hydrolase and glutaminyl-peptide cyclotransferase genes^40^. In the two species, the other breakpoint is located within eight genes of each other (Extended Data Fig. 24). The similarities in genomic architecture between *C. histrio* and *Hel. numata* demonstrate striking parallel evolution between lineages that diverged ∼120 MYA not only in gene usage but also in genetic architecture and the location of the inversion mutations.

### Limited evidence for introgressive allele sharing

Convergent evolution among very closely related *Heliconius* species often results from the sharing of adaptive alleles among species via occasional hybridisation. Both the forewing yellow bar and the hindwing black mimetic phenotypes are found in multiple *Melinaea* and *Hypothyris* species (and to a limited extent in *Mechanitis*) allowing us to test the extent to which allele sharing via introgression is responsible for colour pattern convergence across a wider range of non-*Heliconius* species comparisons. In addition to the species used in the GWAs, we analysed the genomes of 222 individuals of 24 species (8 *Melinaea*, 9 *Hypothyris* and 7 *Mechanitis* species); Extended Data Figs. 3-9 and Table S4). Although we detect ongoing interspecific gene flow in sympatry among many species in all three genera (Extended Data Fig. 25), inspection of all significantly associated GWA SNPs at both *ivory* and *optix* regions failed to detect instances where the GWA SNP alleles were shared with congeneric species with mimetic phenotypes (Extended Data Figs. 3-9). Twisst^41^ and Relate^42^ analyses also failed to detect signals of allele sharing (Extended Data Figs. 26-28). The only exception is in *Melinaea* where narrow signals of introgression near *optix* are present among multiple species sharing melanic hindwing patterns (Extended Data Figs. 28 and 29). Despite the *ivory* and *optix* regions being repeated targets of selection across multiple species, and evidence for interspecific gene flow at the genome-wide level^43^ (Extended Data Figure 25), we do not find evidence of long-term balancing selection maintaining diversity at these regions in *Mechanitis*, *Melinaea* or *Hypothyris* (Extended Data Figs. 30-32). Although we cannot rule out allele sharing because it is possible that our top GWA SNPs do not include the causative genetic variants, in the Ithomiine genera examined we do not find the relatively long introgressed haplotype blocks that characterise mimetic convergence among closely-related *Heliconius* species, despite ongoing interspecific gene flow in both groups^23,24^. While most *Heliconius* species have a highly conserved karyotype^44^, the frequent karyotypic differences present even between closely-related ithomiine species may generate intrinsic postzygotic isolation which limits introgression^43,45,46^.

### Functional characterisation in *Mechanitis messenoides*

To confirm that the genes nearest the GWA peaks, *ivory* and *optix*, are causally involved in colour pattern formation in ithomiine butterflies, we performed CRISPR-Cas9 gene knock-outs in *Mec. messenoides*. In mosaic *ivory* knockout individuals, black and orange scales turn yellow, and *optix*-knockout causes orange scales to turn black (Fig. 4b and Extended Data Fig. 33; consistent with findings in other butterflies^32,47,48^). *In situ* hybridization of pupal forewing discs (50 hours post pupation) using *ivory* probes demonstrates that while *ivory* expression occurs across the entire forewing in the non-yellow barred *Mec. messenoides deceptus*, in the yellow barred *Mec. messenoides messenoides*, a region lacking *ivory* expression prefigures the adult forewing yellow bar phenotype (Fig. 4c and Extended Data Fig. 34). In contrast, staining with antibodies against *cortex*, a gene overlapping with *ivory* that has been reported to affect colour patterns in Lepidoptera^34^, shows no spatial association with the forewing yellow bar phenotype, further supporting the role of *ivory* rather than *cortex* in wing melanisation as recently reported for other butterflies^48^ (Extended data Fig. 35). Comparing yellow barred with non-yellow barred individuals, we do not find that *ivory* is differentially expressed in forewing pupal wing discs of *Mec. messenoides* or *Mel. menophilus* (Extended Data Fig. 36). However, this is not unexpected as whole forewing tissues were used and the anticipated difference in *ivory* expression is ∼20% (proportional to the area of the yellow bar region compared to the whole wing) and thus not detectable in these experiments.

In *Mec. messenoides*, only 5 SNPs located in a ∼1.5 kb interval (SUPER6:6,877,302-6,878,798) are fully associated with the yellow-bar phenotype. These are candidate binding sites for transcription factors controlling *ivory* expression. In this interval we find eight sequence motifs which contain a fixed SNP and are present in all homozygotes of one subspecies and absent in all homozygotes of the other subspecies. Of these, four correspond closely to binding sites of known transcription factors that are also expressed in pupal wing discs of *Mec. messenoides* (Sox15, Ftz-F1, ttk and br-Z4; Extended Data Fig. 37). Sox15 is potentially implicated in the differentiation of lepidopteran scale cells^49^. In *Junonia* butterflies, Ftz-F1 has been shown to bind to the *ivory* promoter^32^. These SNPs and transcription factors are strong candidates for future investigation of transcriptional control of *ivory*.

### Repeatable and predictable evolution

The neotropical tiger mimicry ring is exceptionally species-rich, comprising over 100 species from divergent lepidopteran lineages. In this system the underlying genetic architecture has played a fundamental role in shaping the evolutionary trajectory at both the macro and microevolutionary levels. Phenotypic convergence in all species studied in this mimicry ring is characterised by a simple genetic architecture where a few large effect genes, primarily *ivory* and *optix*, are reused repeatedly. Convergent phenotypic switches occur via independent mutations in the regulatory regions of these genes rather than reuse of standing genetic variation^10^ or alleles shared via introgression from other species^12,23^.

While *ivory* and *optix* are known to control colour patterning across Lepidoptera, when controlling highly similar phenotypes, evolution is surprisingly predictable with convergence caused by recurrent mutations at very similar regions of these genes. The repeatability of evolution also extends to the similar inversion architectures that maintain different allelic combinations in the locally polymorphic butterfly and moth species *Hel. numata* and *C. histrio*. These results suggest that developmental pathways controlling the convergent phenotypes are highly constrained. The different tiger sub-mimicry rings represent locally adaptive fitness peaks. Our results show that the paths to reach these peaks are not only constrained, but also that the steps along these paths are few and large in size, i.e. using large-effect loci. The limited number of paths leading to these fitness peaks may enable diverse taxa to more easily join this species-rich mimicry ring. Once occupying a particular fitness peak, species may then jump via regulatory changes with few or no pleiotropic effects to alternate peaks representing other locally prevalent colour patterns. The outcome of “replaying life’s tape” has been a longstanding question in evolutionary biology^3^. Our repeated discovery of convergent adaptation via narrow and predictable pathways over 120 million years suggests that the running of this tape may be more predictable than expected.

## Methods

### Sample collection and DNA extraction

Samples were collected from across Ecuador, Peru and Colombia (Extended Data Figures 17-23; Supplementary Table S1) between 2002-2024. Wings were removed from specimens and kept as vouchers in glassine envelopes. For most samples the bodies were preserved in NaCl-saturated DMSO solution and stored at -20°C. Some specimens, including those used for genome assembly, were flash frozen in liquid nitrogen and stored at -80°C. Single dried legs were used for a few specimens of *Melinaea mothone* and *Melinaea isocomma* held at the Natural History Museum London. Museum legs were extracted using a Lysis-C buffer and a Qiagen MinElute DNA extraction kit (adapted from^50^). Other DNA extractions were carried out using the Qiagen DNeasy Blood and Tissue Kit, Qiagen MagAttract High Molecular Weight kit, QiaAmp DNA mini kit, or a PureLink digestion and lysis step followed by a magnetic bead DNA extraction^51^. DNA concentration was quantified using Qubit Fluorometer (Invitrogen) and purity assessed using NanoDrop (Thermo Fisher Scientific). For each sample, 350 bp fragment libraries were made using NEBNext Ultra II FS Kit, or using TN5-transposase-mediated tagmentation (adapted from^52^), or following the manufacturer’s guidelines with the Illumina DNA PCR-free library prep kit and sequenced (150 bp paired-end) on Illumina NovaSeq 6000 or NovaSeq X machines.

### Phenotyping wings

Wings of specimens were photographed and used to score the colour pattern phenotypes. *Chetone histrio* individuals were classed as *Chetone histrio histrio* and *Chetone histrio hydra* with no intermediates (Extended Data Fig. 23). Yellow bar phenotypes in *Hypothyris anastasia*, *Mechanitis messenoides*, *Melinaea menophilus* and *Melinaea mothone* were scored as mainly present or absent, with a small number of intermediates (Extended Data Figures 17-21). Hindwing orange/black in *Mechanitis messenoides*, *Melinaea menophilus, Melinaea marsaeus* and *Hypothyris anastasia* were scored as present or absent. The black in the wing tip and base of the forewing in *Mechanitis messenoides* was scored as present or absent (Extended Data Figure 22). The apical spot on the forewing of *Melinaea menophilus* was scored as present or absent (Extended Data Figure 14). Phenotype scores are shown in Supplementary Table S1.

### Quantitative phenotyping of *Mechantis messenoides* orange/black colouration

The dorsal side of hindwings of *Mec. messenoides* were photographed within a uniformly lit lightbox. The background was removed from the raw images, and, in cases where a part of the wing was missing, the corresponding wing from the other side was mirrored and used to fill in any gaps. Areas of the three predominant colours (black, yellow, orange) were then made uniform using CorelDraw. Wings were aligned and the pattern of black colouration was analysed using Patternize^53^. The extracted black colour pattern was analysed using PCA. PC1 explained 32% of variation in the hindwing black pattern, and the resulting eigenvectors were used as phenotypic values in a GWA analysis (see below).

### Reference genome assembly

For *Hypothyris anastasia* and *Chetone histrio histrio*, DNA was extracted from Ecuadorian flash-frozen females using a standard phenol-chloroform protocol and samples sequenced to ∼150✕ coverage using PromethION R9.4.1 flow cells (Oxford Nanopore Technologies). Genome assembly carried out using NextDenovo 2.5.2^54^ and polished with additional Illumina sequence using NextPolish1.4.1^55^. The *Hyp. anastasia* assembly was scaffolded with HiC data generated using the Arima Hi-C+ kit and sequenced on HiSeq X using YaHS^56^. To screen non-insect sequences in the assemblies, we divided the genome into 10 kb windows and performed BLASTN^57^ searches against the GenBank complete nt database^58^. Scaffolds with predominantly non-insect matches were removed from the assembly. Additionally, we trimmed scaffolds showing strong similarity to non-insect sequences. Both assemblies showed high contiguity, with scaffold N50 values of 5.8 Mb for *C. histrio histrio* and 37.2 Mb for *Hyp. anastasia*. Genome completeness was also high, with 97.4% and 98.1% of single-copy BUSCO genes^59^ found in *C. histrio histrio* and *Hyp. anastasia* respectively. Additional genome assemblies for *Melinaea marsaeus*^43,60^, *Mechanitis messenoides*^43^ and *Melinaea mothone*^43^ were used for mapping the respective genome sequence data.

### Whole-genome resequencing and genotype calling

lllumina adapter sequences were trimmed from the raw FASTQ files using Cutadapt 1.2.1^61^, with further trimming with a minimum window quality score of 20 using Sickle 1.2^62^. Reads shorter than 15 bp were removed. Trimmed reads were mapped with BWA v0.7.17^63^ using BWA mem and default options against the reference genome of their respective species. Mapped reads were sorted using Samtools v1.15^64^, and Picard 2.25.5 (http://broadinstitute.github.io/picard/) used to add read groups and mark duplicates. The SNP calling was performed using GATK v4.1.3^65^. The GATKHaplotypeCaller was used to generate GVCF files followed by genotyping using GATK GenotypeGVCFs. VCF files were filtered using Bcftools v1.19^64^ to extract biallelic SNPs with a variant quality score (QUAL) ≥ 10, a genotype quality (GQ) ≥ 10 and a depth of coverage ≥ 5. SNPs with more than 20% of missing data were removed. We imputed missing data and phased the VCF files for each species using SHAPEIT v4.2^66^ with default parameters.

### BUSCO phylogeny and divergence times

To infer the phylogeny and divergence time between the major groups analyzed in this study, we built a dated phylogenetic tree of 18 species: *Napeogenes inachia, Hypothyris anastasia, Ithomia salapia, Scada zibia, Methona curvifascia, Mechanitis messenoides*, *Melinaea mothone*, *Melinaea menophilus*, *Melinaea marsaeus*, *Danaus plexippus, Heliconius pardalinus, Heliconius melpomene*, *Heliconius numata*, *Heliconius erato*, *Bicyclus anynana*, *Chetone histrio*, *Biston betularia* and *Plutella xylostella*) using BUSCO genes.

For each reference genome, we ran BUSCO v5.4.3^59^ and extracted the BUSCO genes common to all species. Sequences for each gene were translated into amino acids and aligned using MUSCLE v3.8.31^67^. The alignment was then reverse-translated to nucleotides using PAL2NAL v14^68^, retaining only genes with fewer than 2% gaps, resulting in a dataset of 257 genes.

We inferred the maximum likelihood (ML) tree based on these concatenated common genes using RAxML v8.2.12^69,70^. We generated 100 bootstrap alignments using the *-f j* option in RAxML and optimized the model parameters and branch lengths of these bootstrapped trees based on the previously inferred ML tree using the *-f e* option. All trees were rooted using pxrr v1.3.1^70^ with *Plutella xylostella* as the outgroup.

Divergence time estimates were obtained using a penalized-likelihood-based approach implemented in TreePL v1.0^71^. The node separating Papilionidae from the moths was used as a calibration node, constraining it to range from 100 to 120 Mya^72^. TreePL was run on each bootstrapped tree to obtain age estimate ranges for each node. The priming step was performed on each of the 100 bootstrapped trees, cross-validation was run 10 times, and finally, for the dating step, the best smoothing parameters for each run were chosen based on the lowest χ2 value and the most common value out of the 10 runs. We used the TreeAnnotator utility from the BEAST package^73^ to calculate the 95% highest posterior density for the node ages using a burn-in of 10%.

### Assessing population structure

Population structure within the taxa involved in each genome-wide association analysis was assessed via Principal Component Analysis (PCA) in PLINK v1.9^74^ using an LD-pruned SNP dataset. LD pruning was performed in PLINK using a window size of 100 SNPs, a window shift of 10 SNPs and an r² value of 0.1.

### Genome wide association mapping

We investigated the genetic basis of phenotypic differences using a genome-wide association (GWA) approach^75^. To identify single-nucleotide polymorphisms (SNPs) associated with each trait, we applied linear univariate mixed models in GEMMA v0.98.5^76^. SNPs were filtered to retain only those with a minor allele frequency ≥10% and missingness <25%. To account for multiple testing, we applied a Bonferroni correction. Sample relatedness was controlled for by incorporating a pairwise relatedness matrix as a covariate in the model. All other parameters were set to default values.

To annotate the association peak regions, we predicted genes within a 250 kb interval around each peak using AUGUSTUS v3.5.0^77^, trained on the *Heliconius melpomene* annotation. We then performed BLASTP^57^ searches against the UniProt database^78^ to identify and annotate the genes. For SNPs exceeding the significance threshold, we calculated squared Spearman’s rank correlation coefficients (ρ²) to quantify the strength of association between SNP genotypes and phenotypic traits, as both were encoded categorically.

GWA analyses were performed for the presence/absence of a forewing yellow bar in *Melinaea mothone* (N = 49), *Melinaea menophilus* (N = 64), *Mechanitis messenoides* (N = 111), and *Hypothyris anastasia* (N = 61). For the hindwing, GWA was conducted in *Melinaea marsaeus* (N = 40), *Mechanitis messenoides* (N = 102), and *Hypothyris anastasia* (N = 61), comparing solid black vs. striped-black phenotypes. For *Melinaea menophilus* (N = 67), we compared three hindwing phenotypes: solid black (encoded as 1), striped-black (0.5), and completely orange (0). For *Mechanitis messenoides*, hindwing black pattern variation is somewhat more continuous rather than strictly discrete. We also performed an additional GWA using quantitative phenotype values obtained using Patternize^53^, producing results consistent with the manual classification (Extended Data Fig. 38). We also conducted targeted GWA in *Mechanitis messenoides* to investigate black-orange patterning at both the forewing base (N = 111) and tip (N = 111; Extended Data Fig. 10), as well as in *Melinaea menophilus* (N = 67) to assess the presence/absence of an apical spot on the forewing (Extended data Fig. 14).

### GWA peak alignments

For each pair of species, we aligned the regions around the GWA peaks to assess whether these association peaks fall within homologous genomic regions. We limited the alignment to the two genes flanking the GWA peaks. The alignments were performed using Nucmer from the MUMmer package v3.23^79^. For each genome pair, one genome was divided into non-overlapping sliding windows of 1000 bp. These windows were then individually aligned to the alternative genome. Due to the high divergence between genomes, we ran nucmer using the following flags: --mum -c 20 -b 500 -l 10 --maxgap 500.

### QTL mapping in Heliconius pardalinus

Crosses and sequencing of hybrids between *Hel. pardalinus butleri* and *Hel. pardalinus sergestus* are described in^38^. Dorsal surfaces of wings from 82 backcross hybrids were photographed in a standardized light box against a white background using a Canon EOS D1000 together with an X-rite ColorChecker® Mini to enable color calibration. Yellow and orange forewing patterning, along with orange hindwing patterning were quantified using a standardized patternize workflow as follows. A reference image was selected, and the RGB color signature of a key pattern element was extracted using the sampleRGB() function. Images were aligned to this reference to standardize spatial orientation using the patRegRGB() function, with a color offset (colOffset = 0.15) and background removal threshold (removebg = 100) to isolate the focal pattern. To quantify variation in color pattern distribution among individuals, we performed a PCA on the aligned pattern rasters using patPCA(). The resulting PCA scores were subsequently used for quantitative trait locus (QTL) mapping as described in^39^.

### Butterfly husbandry

Wild *Mechanitis messenoides messenoides* and *Mechanitis messenoides deceptus* individuals were caught with nets in the Napo province of Ecuador, and used to establish breeding stocks in outdoor insectaries at Ikiam Regional Universidad Amazonica. The adults were fed sucrose solution and had access to *Lantana* and Asteraceae flowers. *Solanum quitoense* was used for oviposition and rearing larvae.

### CRISPR-Cas9 genetic modification

*Ivory* and *optix* were annotated in the reference genome of *Mechanitis messenoides* (ilMecMess1.1.primary.fa^43^) and *ivory* in *Melinaea mothone* (ilMelMoth8.1.primary.fa^43^) based on manual curation of BLAST-hits with the corresponding genes from *Danaus plexippus and Heliconius erato*. RNA-guides (sgRNA) were designed against the annotated *ivory*- and *optix*-genes using Geneious (www.geneious.com) (Supplementary Table S7). Eggs were collected and arranged with non-toxic glue on a microscope slide. The eggs were injected with a 1:1 mixture of the sgRNA (Sigma Aldrich) and Cas9-protein (TrueCut Cas9 Protein V2, Invitrogen) at 1 ug/ul within 3-4 hours of laying following established protocols^80^.

### *In situ* hybridisation with HCR

Wing tissues were dissected at different developmental timepoints (5th instar caterpillar, day 1-4 after pupation). Caterpillars and pupae were anaesthetised on ice before dissection in cold phosphate-buffered saline (PBS). Dissected wing tissue was fixed for 30-40 minutes with formaldehyde (0.25 ml 37% formaldehyde with 0.75 ml PBS 2mM ethylene glycol tetraacetic acid), and subsequently dehydrated and stored in methanol at -20 °C following the protocol of ^81^ until the ‘post-fixation’ step. Subsequently, the HCR *in situ* protocol of Molecular Instruments (MI-Protocol-RNAFISH-GenericSolution) was followed. The wings were also stained with DAPI/HOECHST antibody to visualise the nucleus (Sigma-Aldrich; 10236276001), mounted on a slide in 60% glycerol, and imaged with a Leica SP8 confocal microscope.

### *Cortex* antibody staining

Dissected wing tissues were collected with a fixation (∼30-40 minutes) in 4% paraformaldehyde in PBS 2mM EGTA. The wings were stored in PT-BSA (PBS 0.1% Triton X-100 with 0.1% sodium azide and Bovine Serum Albumin (0.05g in 10ml)). The samples were subsequently washed and stained according to a protocol adapted from^82,83^. The primary antibodies against *Cortex* were made for *Heliconius* (the same antibodies as^83^; rabbit), with secondary antibodies goat anti-rabbit AlexaFluor-555 (ThermoFisher; A-21428). The wings were also stained with Wheat Germ Agglutinin (WGA; plasma membrane; Cambridge BioScience BT29022-1; CF®488A Conjugate) at 488 nm and DAPI/HOECHST for the nuclear DNA at 405 nm (Sigma-Aldrich; 10236276001). The tissues were imaged as for the HCR.

### RNA sequencing for annotation of *ivory*

RNA-seq nanopore long reads were generated for 12 *Melinaea menophilus* wing discs (Day 2 after pupation), comprising 3 forewing and 3 hindwing disc samples each from two subspecies: *ssp. nov. 1* and *hicetas*. Nanopore sequences were also generated from forewing disc tissues (Day 2 after pupation) for 3 samples each of *Mechanitis messenoides messenoides*, *Mechanitis messenoides deceptus* and *Melinaea mothone mothone*. Wing discs were dissected out and either flash-frozen (*Mechanitis messenoides* and *Melinaea mothone*) or stored in RNAlater (*Melinaea menophilus*) at -70°C until processing. Following extraction, RNA quality was assessed using the Agilent Bioanalyzer. Oxford Nanopore Technologies full-length cDNA sequencing libraries were prepared using the ONT cDNA-PCR Sequencing V14 Barcoding kit (SQK-PCB114.24). Barcoded libraries, with cDNA pooled at equimolar concentrations, were sequenced on R10 flow cells (FLO-PRO114) using an ONT PromethION sequencer running MinKNOW version 24.02.10. Super-accuracy basecalling was performed with ONT’s Dorado software version 7.3.9. The reads from each sample were aligned to their respective reference genomes using minimap2 v2.26^84^ with the command “minimap2 -ax splice -uf -k14”. The resulting BAM files were visualized in IGV^85^ with the junction track option, focusing on the *ivory* region. Splicing events supported by at least five reads and present in at least two individuals were retained to define the exons and isoforms of *ivory*.

### Differential gene expression in *Melinaea menophilus*

Nanopore reads were mapped to the *Melinaea menophilus* reference genome using Minimap2 v2.26^84^ with parameters optimized for long-read spliced alignment. The resulting BAM files were then used to assemble transcripts for each sample individually with StringTie v3.0.0^86^, using the -L option to account for long reads. These individual transcript assemblies were subsequently merged into a unified transcriptome to generate a consensus annotation. Read quantification, with parameters optimized for long-read data (-L -s 0 -M --fraction -O), was performed using featureCounts v2.0.4^87^ to count reads assigned to genes and isoforms inferred by StringTie. Because automated quantification of long non-coding RNA is often unreliable, the automatically assigned read counts for *ivory* were manually removed and replaced with curated counts obtained using IGV. Differential gene expression in *Melinaea menophilus* was analyzed by comparing forewing and hindwing datasets for both subspecies (*ssp* vs *hicetas*). Three replicates were included for each condition. Read count data were processed using DESeq2 v1.38.3^88^. Raw count data were normalized and differential expression was assessed using the Wald test with a Benjamini-Hochberg correction for multiple testing. Genes with an adjusted *p*-value (*padj*) < 0.05 were considered significantly differentially expressed. We then specifically examined whether *ivory* was differentially expressed between the yellow-barred and non-yellow-barred forewings.

### Differential gene expression in *Mechanitis messenoides*

Fore- and hindwings were dissected at six different timepoints (early 5th instar, late 5th instar, Day 1 after pupation to Day 4 after pupation), for three forewings and hindwings each for both *Mec. messenoides deceptus* and *messenoides* (6 ✕ 3 ✕ 3 ✕ 2 = 72 samples). Each wing was collected separately and flash frozen in liquid nitrogen. RNA was extracted with a MagMAX Mirvana Total RNA isolation kit, following the manufacturer’s protocol (Thermo Fisher Scientific: AM1830). Libraries were 150bp paired-end Illumina sequenced on two Novaseq X 10B lanes. Illumina reads were trimmed with FastP v0.23.2^89^ and RCorrector^90^, before using STAR without an annotation in ‘two pass mode’ v2.7.9a ^91^ to align them to the *Mec. messenoides* reference genome. Based on the STAR-alignments, a *de novo* transcriptome was assembled using Trinity v2.15.1^92^ with the ‘genome_guided_bam’ option. Read quantification was performed using Salmon v1.10.2^93^. Differential expression in the forewing and hindwing datasets was then assessed for both subspecies (*messenoides* vs *deceptus*) using the same approaches as for *Mel. menophilus* (see previous paragraph).

### Measuring genome-wide introgression using *f_4_* statistics

To investigate potential recent introgression events between sympatric *Melinaea, Mechanitis* and *Hypothyris* species, we employed the *f_4_*-statistics framework implemented in ADMIXTOOLS v2.0.8^94^. *f_4_*-statistics identify deviations in allele frequency correlations across populations or species, which can provide evidence for admixture or introgression. Specifically, we computed *f_4_*-statistics between pairs of populations of different species in the same location versus between pairs of populations of the same species in different locations. We designed the test so that a positive f*_4_*-statistic indicates excess allele sharing between sympatric species, consistent with gene flow. Statistical significance was assessed using a block-jackknife approach, dividing the genome into non-overlapping 500 kb blocks.

### Measuring introgression at colour genes

We tested for evidence of introgression at wing color pattern genes through inference of local genealogies consistent with introgression, and complemented this approach with visualization of genotypes at top GWA SNPs. These analyses were applied independently to *Mechanitis*, *Melinaea*, and *Hypothyris*, testing whether species with similar phenotypes (i.e., co-mimetic species) showed evidence of gene flow at color loci. F*_d_* and F*_dM_* statistics^95,96^ did not reveal any evidence of introgression at candidate color loci. We therefore focused on alternative methods that provide more localized or topology-based insights into gene flow in these regions.

To assess introgression at GWA peaks, we employed two complementary approaches to infer local genealogies: (1) marginal trees at individual SNPs using Relate^42^, and (2) Neighbor-Joining (NJ) trees built from non-overlapping 100-SNP sliding windows, using the ape R package^97^. Both methods were applied to a core set of quartets, and some additional quartets were analysed using the NJ approach (Extended data Fig. 26-28). The two analyses followed a common framework for summarizing topologies and assessing significance.

#### Quartet Definition and Taxon Selection

Analyses were conducted across four different taxon combinations (see Table 4). Each quartet ideally consisted of two species, each represented by two morphologically distinct subspecies (spA: sspA1/sspA2 and spB: sspB1/sspB2). We specifically focused on introgression between two distant species with similar wing phenotypes, particularly between sspA1 and sspB1. When this configuration was not possible, we modified the arrangement to include one taxon (sspB1) for introgression testing with sspA1 and a third outgroup (sspC1). In both quartet types, we interpreted clustering of sspA1 and sspB1 near the GWA peak as evidence of potential introgression between these species.

#### Relate based inference

Relate was run on 4–10 Mb regions centered on each GWA peak, comparing species involved in the GWA and closely related, phenotypically similar species (Extended data Fig. 26-28). VCFs were imputed and phased with SHAPEIT v4.2^66^, and ancestral alleles were inferred using an outgroup species - *Hyalyris antea* and *Hyalyris lactea* for *Hypothyris*; *Forbestra equicola, F. proceris* and *F. olivencia* for *Mechanitis*; and *Melinaea ludovica* for *Melinaea*. For SNPs missing in the outgroup, we used the most frequent allele across the genus. Relate was run with an effective population size of 1x10^7^ and a mutation rate of 2.9x10^-9^ per site per generation, based on estimates from *Heliconius* studies^98–100^. Because Relate requires a genetic map, we used a uniform recombination rate of 6 cM/Mb, generated with a script available at: https://github.com/joanam/scripts/blob/master/createuniformrecmap.r.

#### Sliding Window NJ Tree Inference

As an alternative to Relate, we inferred genealogies by constructing Neighbor-Joining (NJ) trees from non-overlapping 100 SNP sliding windows using the ape R package. This window-based approach offered a complementary view of local genealogies by summarizing phylogenetic signal across short genomic regions. For this analysis, we used separate VCFs that retained multiallelic SNPs. NJ tree inference was applied to the same taxon quartets used in the Relate analysis. As this test can be run using a small sample size, additional quartets based on non-GWA species were also analysed.

#### Summarizing Tree Topologies and Assessing Significance

The results from both Relate and the sliding window NJ tree inference were summarized using Twisst^41^. To assess whether a region exhibited a significant excess of introgression-compatible topologies, we performed a permutation test using block-shuffling^101^. This method disrupts potential signals of introgression whilst preserving the local genomic structure, by randomly shuffling 100 kb blocks across the entire region. The null distribution for introgression topology was derived by counting the number of windows with a Twisst weight of at least 0.95 for the introgressed topology (referred to hereafter as intro95). A p-value was calculated by comparing the observed intro95 count in the GWA peak region to the null distribution, based on 50,000 permutations. This approach was applied to both the Relate and sliding window NJ tree analyses to assess introgression significance consistently across both methods.

#### Genotypes matrices

We also created a genotype matrix for the most strongly associated GWA SNPs (Extended Data Table 2) to determine whether the same SNPs associated with wing color patterns in one species are also linked to similar phenotypic traits in co-mimetic species. This approach allows us to detect subtle signals of introgression, particularly in cases where the local genealogy analyses might not identify introgression due to the signal being present in a very narrow window.

### Balancing selection

To assess whether colour pattern genes are targets of long-term balancing selection, we examined patterns of polymorphism and allele frequency across *Mechanitis*, *Melinaea*, and *Hypothyris* species. To test for balancing selection at the colour genes of interest across species, we used three approaches: multispecies nucleotide diversity, detection of trans-species polymorphisms, and the MuteBaSS method^102^. For *Melinaea*, analyses were conducted at *ivory*, *optix*, and *antp*, whereas in *Mechanitis* and *Hypothyris*, analyses focused on *ivory* and *optix*.

#### Multispecies nucleotide diversity

To quantify polymorphism at colour genes and surrounding regions, we calculated multispecies nucleotide diversity by pooling data across species and subspecies. We used Pixy v1.2.5^103^ to compute nucleotide diversity in non-overlapping 50 kb sliding windows. Analyses were performed on repeat-masked VCF files including invariant sites as well as bi- and multiallelic SNPs. Repetitive elements in the Lepidoptera genomes were identified using RepeatModeler v2.0.4^104^, and subsequently masked with RepeatMasker v4.1.2^105^ using the generated repeat library. We calculated nucleotide diversity under two conditions:

1. Analysis including all species.
2. Pairwise comparisons between individual species/subspecies to identify cases where balancing selection signals is limited to a subset of taxa.

Detailed species groupings for *Melinaea*, *Mechanitis* and *Hypothyris* are provided in supplementary table 6.

#### Trans-Species Polymorphisms

To test for trans-species polymorphisms across multiple species, we analyzed genomic variation shared across multiple species using a custom python pipeline based on *pysam* v0.21.0. We used the same repeat-masked VCF-files as described in the ‘multispecies nucleotide diversity’ section.

A site was classified as transpolymorphic if it met the following criteria:

1. at least four species were polymorphic (when more than three species were present);
2. all species were polymorphic (if three or fewer species were present);
3. each allele was represented by at least six total copies across species.

#### MuteBaSS

To further test for local signals of balancing selection, we used MuteBaSS v1.0^102^, which detects balancing selection without relying on trans-species polymorphic sites. We used MuteBaSS to compute the multispecies NCD, NCDsub, and NCDopt statistics, as well as the trans-HKA test. To capture the potentially narrow signals of balancing selection, all statistics were calculated in 1 kb windows with a 500 bp sliding step. Ancestral states were inferred as described in the *Relate based inference* section. For *Melinaea* and *Mechanitis*, we assumed SNPs followed the phylogeny proposed by Van der Heijden et al. 2024^43^. For *Hypothyris*, we used the phylogeny from Chazot et al. 2019^106^.

### Transcription factor binding site analysis

In all cases excluding the inversion in *Chetone histrio*, the colour-pattern associated regions identified by GWA fall outside of the genes of interest. This may suggest that the associated regions control colour-pattern by influencing gene regulation, potentially by affecting transcription factor binding. As the peaks of association with the yellow-bar phenotype are narrow and contain only a few fixed SNPs, it may be possible to associate these SNPs with a specific transcription factor binding site, by identifying binding motifs which are enriched consequences of one colour pattern form compared to the other. To test this we focused on our best sampled species, *Mechanitis messenoides*. Multiple analyses were performed to uncover transcription factor (TF) binding sites within and around the *ivory* GWA peak region, defined here as the ∼1.5kb region between the weakly-associated SNPs which flank the associated region identified through GWA (Extended Data Figure 2). Only individuals homozygous for the fixed SNPs identified in the GWA analysis were selected. A fasta file for the region of interest was generated for each individual using the consensus commands in either samtools or bcftools. Homer^107^ was used to identify *de novo* DNA motifs which are enriched in one set of sequences compared to a control group, using the findMotifs.pl script and the insect known motif collection. The two forms (yellow-barred vs non yellow-barred) were alternately used as both control and target sequences.

To locate and identify known TF binding sites within the peak region, the FIMO algorithm in MEME Suite^108^ was run on all sequences, including both forms, for each species. A background file was generated based on the entire chromosome containing the peak region (SUPER6), using the fasta-get-markov command. The expression of detected motifs was checked by examining expression levels of the corresponding transcription factors within the *Mechanitis messenoides* RNAseq data.

## Data availability

DNA and RNA sequence data and genome assemblies will be uploaded to ENA. Phylogenies and wing images will be made available in a Dryad repository.

## Code availability

More detailed information and the associated scripts are available at the following link: https://github.com/yacinebenchehida/Ithominii_convergence/blob/main/

## Acknowledgements

This work was funded by NERC grant NE/T008121/1 to K.K.D.; NERC ACCE DTP PhD studentship to E.J.P.; E.v.H.; a Wellcome Trust award 220540/Z/20/A, a Branco Weiss Fellowship, a Royal Society University Research Fellowship (URF\R1\221041) and a Bateson Research Fellowship by St John’s College, Cambridge awarded to J.I.M.; a ANR grant (SPECREP - ANR-14-CE02-0011-01) awarded to M.E. The University of York Viking2 HPC and the Wellcome Sanger farm HPC were used for bioinformatics analyses. We thank SERFOR, the Peruvian Ministry of Agriculture and the Área de Conservación Regional Cordillera Escalera (0289-2014-MINAGRI-DGFFS/DGEFFS, 020-014/GRSM/PEHCBM/DMA/ACR-CE and 040–2015/GRSM/PEHCBM/DMA/ACR-CE) for collecting permits; the Ministerio del Ambiente and Museo Ecuatoriano de Ciencias Naturales in Ecuador (MAATE-DBI-CM-2021-176 and MAATE-DBI-CM-2023-0286) for collecting permits. Field collections in Colombia were conducted under permit no. 530 issued by the Autoridad Nacional de Licencias Ambientales of Colombia (ANLA). We thank Mathieu Joron and Stephen Montgomery for providing samples, Ismael Aldas for support with fieldwork, Nicol Rueda for help with the Relate analysis and Chris Thomas for comments on the manuscript.

## Author contributions

Y.B.C., E.v.H., and E.J.P. contributed equally to this study. K.K.D., N.J.N., and J.I.M. designed the study. Y.B.C. wrote and implemented most bioinformatics pipelines and carried out the *Chetone* analysis. E.v.H. produced and analysed the *Mechanitis* datasets (with help from M.R.) and led the functional analysis (with help from M.S.P. K.G.G.C. and M.J.S.C). E.J.P. produced and analysed the *Hypothyris* dataset and performed transcription factor binding site analysis. Field collections and insect husbandry in Ecuador were carried out by P.S.C, K.G.G.C, M.J.S.C, F.C,. A.P.A.C, and C.N.B., M.E., J.M., C.S., M.M., G.L., C.J. and N.R. contributed to sample collection. N.R. led the QTL analysis (with the help of K.K.D.). Y.B.C., E.v.H., E.J.P., K.K.D., N.J.N., J.I.M., J.M., C.S. and M.E. wrote and finalized the paper with contributions from all authors.

## References

1. Schluter, D. & Nagel, L. M. Parallel speciation by natural selection. Am. Nat. 146, 292–301 (1995).

2. Blount, Z. D., Lenski, R. E. & Losos, J. B. Contingency and determinism in evolution: Replaying life’s tape. Science 362, eaam5979 (2018).

3. Gould, S. J. Wonderful Life: The Burgess Shale and the Nature of History. (W. W. Norton & Company, 1990).

4. Nadeau, N. J. & Jiggins, C. D. A golden age for evolutionary genetics? Genomic studies of adaptation in natural populations. Trends in Genetics 26, 484–492 (2010).

5. Hsia, C. C. W., Schmitz, A., Lambertz, M., Perry, S. F. & Maina, J. N. Evolution of Air Breathing: Oxygen Homeostasis and the Transitions from Water to Land and Sky. in Comprehensive Physiology vol. 3 849–915 (John Wiley & Sons, Ltd, 2013).

6. Thompson, M., Steichen, J. C. & ffrench-Constant, R. H. Conservation of cyclodiene insecticide resistance-associated mutations in insects. Insect Mol. Biol. 2, 149–154 (1993).

7. Tenaillon, O. et al. Tempo and mode of genome evolution in a 50,000-generation experiment. Nature 536, 165–170 (2016).

8. Stern, D. L. & Orgogozo, V. Is genetic evolution predictable? Science 323, 746–751 (2009).

9. Bohutínská, M. & Peichel, C. L. Divergence time shapes gene reuse during repeated adaptation. Trends Ecol. Evol. 39, 396–407 (2024).

10. Jones, F. C. et al. The genomic basis of adaptive evolution in threespine sticklebacks. Nature 484, 55–61 (2012).

11. Stern, D. L. The genetic causes of convergent evolution. Nat Rev Genet 14, 751–764 (2013).

12. Jones, M. R. et al. Adaptive introgression underlies polymorphic seasonal camouflage in snowshoe hares. Science 360, 1355–1358 (2018).

13. Song, Y. et al. Adaptive introgression of anticoagulant rodent poison resistance by hybridization between old world mice. Curr. Biol. 21, 1296–1301 (2011).

14. Wilson, J. S. et al. North American velvet ants form one of the world’s largest known Müllerian mimicry complexes. Curr Biol 25, R704–6 (2015).

15. Twomey, E. et al. Phenotypic and Genetic Divergence among Poison Frog Populations in a Mimetic Radiation. PLoS One 8, e55443 (2013).

16. Alexandrou, M. A. et al. Competition and phylogeny determine community structure in Müllerian co-mimics. Nature 469, 84–88 (2011).

17. Müller, F. Ituna and Thyridia; a remarkable case of mimicry in butterflies. Proceedings of the Entomological Society of London xx–xxix (1879).

18. Rodríguez, A., Mundy, N. I., Ibáñez, R. & Pröhl, H. Being red, blue and green: the genetic basis of coloration differences in the strawberry poison frog (Oophaga pumilio). BMC Genomics 21, 1–16 (2020).

19. Rubio, A. O. et al. What makes a mimic? Orange, red, and black color production in the mimic poison frog (Ranitomeya imitator). Genome Biol. Evol. 16, (2024).

20. Palmer, D. H. & Kronforst, M. R. A shared genetic basis of mimicry across swallowtail butterflies points to ancestral co-option of doublesex. Nature Communications 11, 1–10 (2020).

21. Nadeau, N. J. Genes controlling mimetic colour pattern variation in butterflies. Curr. Opin. Insect Sci. 17, 24–31 (2016).

22. Joron, M. et al. A conserved supergene locus controls colour pattern diversity in Heliconius butterflies. PLoS Biol. 4, e303 (2006).

23. Heliconius Genome Consortium. Butterfly genome reveals promiscuous exchange of mimicry adaptations among species. Nature 487, 94–98 (2012).

24. Zhang, W., Dasmahapatra, K. K., Mallet, J., Moreira, G. R. P. & Kronforst, M. R. Genome-wide introgression among distantly related Heliconius butterfly species. Genome Biol 17, 25 (2016).

25. Wallbank, R. W. R. et al. Evolutionary Novelty in a Butterfly Wing Pattern through Enhancer Shuffling. PLOS Biology 14, e1002353 (2016).

26. Joron, M. et al. Chromosomal rearrangements maintain a polymorphic supergene controlling butterfly mimicry. Nature 477, 203–206 (2011).

27. Doré, M. et al. Anthropogenic pressures coincide with Neotropical biodiversity hotspots in a flagship butterfly group. Divers. Distrib. 28, 2912–2930 (2022).

28. Beccaloni, G. Ecology, natural history and behaviour of Ithomiine butterflies and their mimics in Ecuador (Lepidoptera: Nymphalidae: Ithomiinae). Tropical Lepidoptera 8, 103–124 (1995).

29. van’t Hof, A. E., Reynolds, L. A., Yung, C. J., Cook, L. M. & Saccheri, I. J. Genetic convergence of industrial melanism in three geometrid moths. Biology Letters (2019) doi:10.1098/rsbl.2019.0582.

30. Hof, A. E. V. et al. The industrial melanism mutation in British peppered moths is a transposable element. Nature 534, 102–105 (2016).

31. Wang, S. et al. The evolution and diversification of oakleaf butterflies. Cell 185, 3138–3152.e20 (2022).

32. Fandino, R. A. et al. The lncRNA regulates seasonal color patterns in buckeye butterflies. Proc Natl Acad Sci U S A 121, e2403426121 (2024).

33. Tian, S. et al. A microRNA is the effector gene of a classic evolutionary hotspot locus. Science 386, 1135–1141 (2024).

34. Nadeau, N. J. et al. The gene cortex controls mimicry and crypsis in butterflies and moths. Nature 534, 106–110 (2016).

35. Van Belleghem, S. M. et al. Complex modular architecture around a simple toolkit of wing pattern genes. Nature Ecology & Evolution 1, 1–12 (2017).

36. Tendolkar, A. et al. Cis-regulatory modes of Ultrabithorax inactivation in butterfly forewings. (2024) doi:10.7554/eLife.90846.

37. Matsuoka, Y. & Monteiro, A. Hox genes are essential for the development of eyespots in Bicyclus anynana butterflies. Genetics 217, 1–9 (2021).

38. Rosser, N. et al. Complex basis of hybrid female sterility and Haldane’s rule in Heliconius butterflies: Z-linkage and epistasis. Mol. Ecol. 31, 959–977 (2022).

39. Rosser, N. et al. Hybrid speciation driven by multilocus introgression of ecological traits. Nature 628, 811–817 (2024).

40. Saenko, S. V. et al. Unravelling the genes forming the wing pattern supergene in the polymorphic butterfly. Evodevo 10, 16 (2019).

41. Martin, S. H. & Van Belleghem, S. M. Exploring Evolutionary Relationships Across the Genome Using Topology Weighting. Genetics 206, 429–438 (2017).

42. Speidel, L. et al. Inferring Population Histories for Ancient Genomes Using Genome-Wide Genealogies. Mol Biol Evol 38, 3497–3511 (2021).

43. van der Heijden, E. S. M. et al. Genomics of Neotropical biodiversity indicators: two butterfly radiations with rampant chromosomal rearrangements and hybridisation. bioRxiv 2024.07.07.602206 (2024) doi:10.1101/2024.07.07.602206.

44. Rueda-M, N. et al. Genomic evidence reveals three W-autosome fusions in Heliconius butterflies. PLoS Genet. 20, e1011318 (2024).

45. Brown, K. S., Jr, Von Schoultz, B. & Suomalainen, E. Chromosome evolution in Neotropical Danainae and Ithomiinae (Lepidoptera). Hereditas 141, 216–236 (2004).

46. McClure, M., Dutrillaux, B., Dutrillaux, A.-M., Lukhtanov, V. & Elias, M. Heterozygosity and Chain Multivalents during Meiosis Illustrate Ongoing Evolution as a Result of Multiple Holokinetic Chromosome Fusions in the Genus Melinaea (Lepidoptera, Nymphalidae). Cytogenet Genome Res 153, 213–222 (2018).

47. Zhang, L., Mazo-Vargas, A. & Reed, R. D. Single master regulatory gene coordinates the evolution and development of butterfly color and iridescence. Proceedings of the National Academy of Sciences 114, 10707–10712 (2017).

48. Livraghi, L. et al. A long noncoding RNA at the cortex locus controls adaptive coloration in butterflies. Proceedings of the National Academy of Sciences 121, e2403326121 (2024).

49. Loh, L. S. et al. Lepidopteran scale cells derive from sensory organ precursors through a canonical lineage. Development 152, (2025).

50. Korlević, P. et al. A Minimally Morphologically Destructive Approach for DNA Retrieval and Whole-Genome Shotgun Sequencing of Pinned Historic Dipteran Vector Species. Genome Biol Evol 13, (2021).

51. Kučka, M. & Frank Chan, Y. HMW DNA Extraction Using Magnetic Beads. https://www.protocols.io/view/hmw-dna-extraction-using-magnetic-beads-b46bq zan (2022) doi:10.17504/protocols.io.b46bqzan.

52. Picelli, S. et al. Tn5 transposase and tagmentation procedures for massively scaled sequencing projects. Genome Res 24, 2033–2040 (2014).

53. Van Belleghem, S. M. et al. patternize: An R package for quantifying colour pattern variation. Methods Ecol Evol 9, 390–398 (2018).

54. Hu, J. et al. NextDenovo: an efficient error correction and accurate assembly tool for noisy long reads. Genome Biology 25, 1–19 (2024).

55. Hu, J., Fan, J., Sun, Z. & Liu, S. NextPolish: a fast and efficient genome polishing tool for long-read assembly. Bioinformatics 36, 2253–2255 (2019).

56. Zhou, C., McCarthy, S. A. & Durbin, R. YaHS: yet another Hi-C scaffolding tool. Bioinformatics 39, btac808 (2022).

57. Camacho, C. et al. BLAST+: architecture and applications. BMC Bioinformatics 10, 421 (2009).

58. Sayers, E. W. et al. GenBank 2023 update. Nucleic Acids Res 51, D141–D144 (2023).

59. Manni, M., Berkeley, M. R., Seppey, M., Simão, F. A. & Zdobnov, E. M. BUSCO Update: Novel and Streamlined Workflows along with Broader and Deeper Phylogenetic Coverage for Scoring of Eukaryotic, Prokaryotic, and Viral Genomes. Mol Biol Evol 38, 4647–4654 (2021).

60. Gauthier, J. et al. First chromosome scale genomes of ithomiine butterflies (Nymphalidae: Ithomiini): Comparative models for mimicry genetic studies. Mol Ecol Resour 23, 872–885 (2023).

61. Martin, M. Cutadapt removes adapter sequences from high-throughput sequencing reads. EMBnet.journal 17, 10–12 (2011).

62. Joshi, N. A. & Fass, J. N. Sickle: A Sliding-Window, Adaptive, Quality-Based Trimming Tool for FastQ Files. (2011).

63. Li, H. & Durbin, R. Fast and accurate short read alignment with Burrows-Wheeler transform. Bioinformatics 25, 1754–1760 (2009).

64. Danecek, P. et al. Twelve years of SAMtools and BCFtools. Gigascience 10, giab008 (2021).

65. Van Der Auwera, G. O. & Connor, B. D. Genomics in the Cloud: Using Docker, GATK, and WDL in Terra. (O’Reilly Media, 2020).

66. Delaneau, O., Zagury, J.-F., Robinson, M. R., Marchini, J. L. & Dermitzakis, E. T. Accurate, scalable and integrative haplotype estimation. Nature Communications 10, 1–10 (2019).

67. Edgar, R. C. MUSCLE: multiple sequence alignment with high accuracy and high throughput. Nucleic Acids Res 32, 1792–1797 (2004).

68. Suyama, M., Torrents, D. & Bork, P. PAL2NAL: robust conversion of protein sequence alignments into the corresponding codon alignments. Nucleic Acids Res 34, W609–12 (2006).

69. Stamatakis, A. RAxML version 8: a tool for phylogenetic analysis and post-analysis of large phylogenies. Bioinformatics 30, 1312–1313 (2014).

70. Brown, J. W., Walker, J. F. & Smith, S. A. Phyx: phylogenetic tools for unix. Bioinformatics 33, 1886–1888 (2017).

71. Smith, S. A. & O’Meara, B. C. treePL: divergence time estimation using penalized likelihood for large phylogenies. Bioinformatics 28, 2689–2690 (2012).

72. Kawahara, A. Y. et al. Phylogenomics reveals the evolutionary timing and pattern of butterflies and moths. Proc Natl Acad Sci U S A 116, 22657–22663 (2019).

73. Bouckaert, R. et al. BEAST 2: A Software Platform for Bayesian Evolutionary Analysis. PLOS Computational Biology 10, e1003537 (2014).

74. Purcell, S. et al. PLINK: a tool set for whole-genome association and population-based linkage analyses. Am J Hum Genet 81, 559–575 (2007).

75. Uffelmann, E. et al. Genome-wide association studies. Nature Reviews Methods Primers 1, 1–21 (2021).

76. Zhou, X. & Stephens, M. Genome-wide efficient mixed-model analysis for association studies. Nat Genet 44, 821–824 (2012).

77. Keller, O., Kollmar, M., Stanke, M. & Waack, S. A novel hybrid gene prediction method employing protein multiple sequence alignments. Bioinformatics 27, 757–763 (2011).

78. The UniProt Consortium. UniProt: a worldwide hub of protein knowledge. Nucleic Acids Res 47, D506–D515 (2018).

79. Kurtz, S. et al. Versatile and open software for comparing large genomes. Genome Biology 5, 1–9 (2004).

80. Zhang, L. & Reed, R. D. Genome editing in butterflies reveals that spalt promotes and Distal-less represses eyespot colour patterns. Nature Communications 7, 1–7 (2016).

81. Martin, A. & Reed, R. D. Wnt signaling underlies evolution and development of the butterfly wing pattern symmetry systems. Dev Biol 395, 367–378 (2014).

82. Martin, A. et al. Multiple recent co-options of Optix associated with novel traits in adaptive butterfly wing radiations. Evodevo 5, 7 (2014).

83. Livraghi, L. et al. -regulatory switches establish scale colour identity and pattern diversity in. Elife 10, (2021).

84. Li, H. Minimap2: pairwise alignment for nucleotide sequences. Bioinformatics 34, 3094–3100 (2018).

85. Robinson, J. T. et al. Integrative genomics viewer. Nature Biotechnology 29, 24–26 (2011).

86. Kovaka, S. et al. Transcriptome assembly from long-read RNA-seq alignments with StringTie2. Genome Biology 20, 1–13 (2019).

87. Liao, Y., Smyth, G. K. & Shi, W. featureCounts: an efficient general purpose program for assigning sequence reads to genomic features. Bioinformatics 30, 923–930 (2013).

88. Love, M. I., Huber, W. & Anders, S. Moderated estimation of fold change and dispersion for RNA-seq data with DESeq2. Genome Biology 15, 1–21 (2014).

89. Chen, S., Zhou, Y., Chen, Y. & Gu, J. fastp: an ultra-fast all-in-one FASTQ preprocessor. Bioinformatics 34, i884–i890 (2018).

90. Song, L. & Florea, L. Rcorrector: efficient and accurate error correction for Illumina RNA-seq reads. Gigascience 4, s13742–015–0089–y (2015).

91. Dobin, A. et al. STAR: ultrafast universal RNA-seq aligner. Bioinformatics 29, 15–21 (2012).

92. Grabherr, M. G. et al. Full-length transcriptome assembly from RNA-Seq data without a reference genome. Nature Biotechnology 29, 644–652 (2011).

93. Patro, R., Duggal, G., Love, M. I., Irizarry, R. A. & Kingsford, C. Salmon provides fast and bias-aware quantification of transcript expression. Nat Methods 14, 417–419 (2017).

94. Maier, R. et al. On the limits of fitting complex models of population history to f-statistics. (2023) doi:10.7554/eLife.85492.

95. Martin, S. H., Davey, J. W. & Jiggins, C. D. Evaluating the Use of ABBA–BABA Statistics to Locate Introgressed Loci. Mol Biol Evol 32, 244–257 (2014).

96. Malinsky, M. et al. Genomic islands of speciation separate cichlid ecomorphs in an East African crater lake. Science 350, 1493–1498 (2015).

97. Paradis, E. & Schliep, K. ape 5.0: an environment for modern phylogenetics and evolutionary analyses in R. Bioinformatics 35, 526–528 (2018).

98. Van Belleghem, S. M. et al. Evolution at two time frames: Polymorphisms from an ancient singular divergence event fuel contemporary parallel evolution. PLOS Genetics 14, e1007796 (2018).

99. Keightley, P. D. et al. Estimation of the Spontaneous Mutation Rate in Heliconius melpomene. Mol Biol Evol 32, 239–243 (2014).

100. Tobler, A. et al. First-generation linkage map of the warningly colored butterfly Heliconius erato. Heredity (Edinb) 94, 408–417 (2005).

101. Anderson M. Winkler, Matthew A. Webster, Diego Vidaurre, Thomas E. Nichols, Stephen M. Smith. Multi-level block permutation. NeuroImage 123, 253–268 (2015).

102. Cheng, X. & DeGiorgio, M. Detection of Shared Balancing Selection in the Absence of Trans-Species Polymorphism. Mol Biol Evol 36, 177–199 (2019).

103. Korunes, K. L. & Samuk, K. pixy: Unbiased estimation of nucleotide diversity and divergence in the presence of missing data. Molecular Ecology Resources 21, 1359–1368 (2021).

104. Flynn, J. M. et al. RepeatModeler2 for automated genomic discovery of transposable element families. Proceedings of the National Academy of Sciences 117, 9451–9457 (2020).

105. Smit, A. F. A., Hubley, R. & Green, P. RepeatMasker Open-4.0. (2013-2015).

106. Chazot, N. et al. Renewed diversification following Miocene landscape turnover in a Neotropical butterfly radiation. Global Ecology and Biogeography 28, 1118–1132 (2019).

107. Heinz, S. et al. Simple combinations of lineage-determining transcription factors prime cis-regulatory elements required for macrophage and B cell identities. Mol Cell 38, 576–589 (2010).

108. Grant, C. E. & Bailey, T. L. FIMO: scanning for occurrences of a given motif. Bioinformatics 27, 1017–1018 (2011).

