## Extended Data Figures 1-38 for "Genetic parallelism underpins convergent mimicry coloration across Lepidoptera"

Extended Data Figure 1: Key to butterfly images used in Figure 1.
Extended Data Figure 2: Genome-wide associations for forewing yellow bar and hindwing orange/black wing phenotypes
Extended Data Figure 3: Genotypic variation at the *Mechanitis messenoides ivory* GWA peak in *Mechanitis* species
Extended Data Figure 4: Genotypic variation at the *Melinaea mothone ivory* GWA peak in *Melinaea* species
Extended Data Figure 5: Genotypic variation at the *Melinaea menophilus ivory* GWA peak in *Melinaea* species
Extended Data Figure 6: Genotypic variation at the *Hypothyris anastasia ivory* GWA peak in *Hypothyris* species
Extended Data Figure 7: Genotypic variation at the *Melinaea marsaeus optix* GWAS peak in *Melinaea* species
Extended Data Figure 8: Genotypic variation at the *Melinaea menophilus optix* GWA peak in *Hypothyris* species
Extended Data Figure 9: Genotypic variation at the *Hypothyris anastasia optix* GWA peak in *Hypothyris* species
Extended Data Figure 10: Genome-wide associations for different black-orange wing phenotypes in *Mechanitis messenoides*
Extended Data Fig. 11: Genotypic variation at the *Mechanitis messenoides optix* forewing base GWA peak in *Mechanitis* species
Extended Data Fig. 12: Genotypic variation at the *Mechanitis messenoides optix* forewing tip GWA peak in *Mechanitis* species
Extended Data Fig. 13: Genotypic variation at the *Mechanitis messenoides optix* hindwing GWA peak in *Mechanitis* species
Extended Data Fig. 14 | Genome wide association for the forewing apical spot in *Melinaea menophilus*
Extended Data Figure 15: Genotypic variation at the *Melinaea menophilus antennapedia* GWA peak in *Melinaea* species
Extended Data Figure 16: QTL mapping intervals for forewing and hindwing pattern variation in *Heliconius pardalinus*
Extended Data Figure 17: *Hypothyris anastasia*: Taxon distribution, wing phenotypes and genetic PCA
Extended Data Figure 18: *Melinaea menophilus*: Taxon distribution, wing phenotypes and genetic PCA
Extended Data Figure 19: *Melinaea marseus*: Taxon distribution, wing phenotypes and genetic PCA
Extended Data Figure 20: *Melinaea mothone*: Taxon distribution, wing phenotypes and genetic PCA
Extended Data Figure 21: *Mechanitis messenoides* (forewing yellow bar and hindwing orange/black): Taxon distribution, wing phenotypes and genetic PCA
Extended Data Figure 22: *Mechanitis messenoides* (forewing base and tip orange/black): Taxon distribution, wing phenotypes and genetic PCA
Extended Data Figure 23: *Chetone histrio*: Taxon distribution, wing phenotypes and genetic PCA
Extended Data Figure 24: Inversion breakpoint analysis in *Chetone histrio*
Extended Data Figure 25: $f_4$ statistics for allele sharing between sympatric species of *Hypothyris*, *Mechanitis, and Melinaea*
Extended Data Figure 26: Testing for introgression across pairs of *Mechanitis* species at *ivory*

Extended Data Figure 27: Testing for introgression across pairs of *Hypothyris* species at *ivory* and *optix*
Extended Data Figure 28: Testing for introgression across pairs of *Melinaea* species at *ivory, optix* and *antennapedia*
Extended Data Figure 29: Genomic signals of introgression at the *optix* locus among four pairs of *Melinaea* species
Extended Data Figure 30: Multispecies balancing selection tests in *Melinaea* at *antennapedia*, *ivory,* and *optix*
Extended Data Figure 31: Multispecies balancing selection tests in *Hypothyris* at *ivory* and *optix.*
Extended Data Figure 32: Multispecies balancing selection tests in *Mechanitis* at *ivory* and *optix*
Extended Data Figure 33: Additional *Mechanitis messenoides* CRISPR *ivory* and *optix* mutants
Extended Data Figure 34: *Mechanitis messenoides in situ* hybridisation for *ivory*
Extended Data Figure 35: *Cortex* expression is not associated with the forewing yellow bar phenotype
Extended Data Figure 36: Differential gene expression analysis in pupal wing discs
Extended Data Figure 37: Transcription factor binding site analysis in *Mechanitis messenoides*
Extended Data Figure 38: *Mechanitis messenoides* GWA for HW black/orange using categorical grouping vs patternize scoring for phenotyping

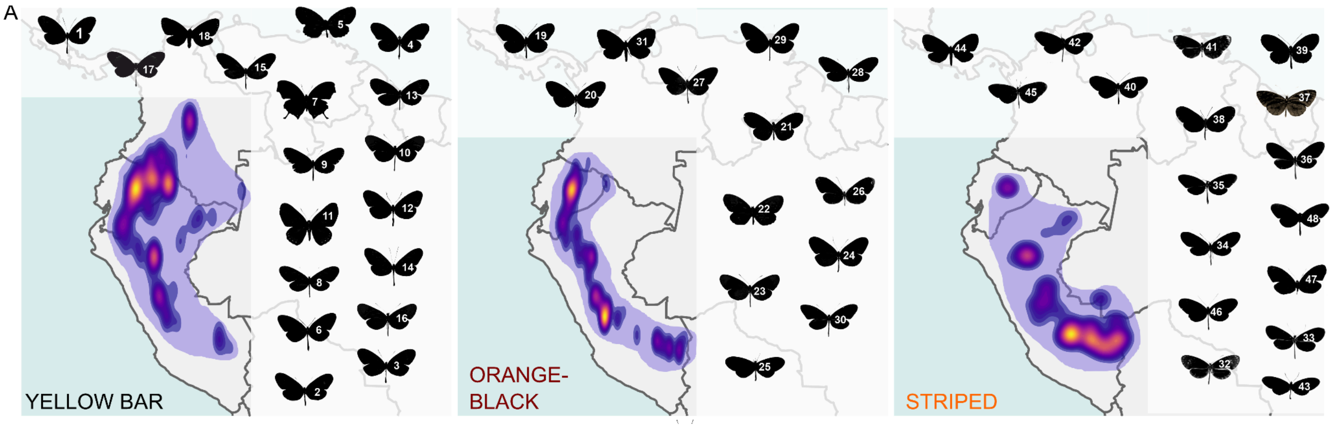


**Extended Data Fig. 1 | Key to butterfly images used in Figure 1a.** 1). *Melinaea menophilus zaneka* 2). *Hypothyris moebiusi moebiusi* 3). *Hypothyris mamercus mamercus* 4). *Heliconius numata euphone* 5). *Eresia pelonia f. callonia* 6). *Melinaea isocomma isocomma* 7). *Consul fabius bogatanus* 8). *Mechanitis mazeus fallax* 9). *Forbestra equicola* 10). *Hypothyris mansuetus amica* 11). *Themone paid trivittata* 12). *Hypothyris cantobrica zamorita* 13). *Hypothyris semifulva ssp.* 14). *Melinaea mnasias abitagua* 15). *Mechanitis messenoides messenoides* 16). *Hypothyris euclea pyrippe* 17). *Melinaea marseus messenina* 18). *Chetone histrio histrio* 19). *Heliconius numata bicoloratus* 20). *Hypothyris semifulva semifulva* 21). *Eueides lampeto* 22). *Eresia pelonia f. Ithomiola* 23). *Napeogenes rhezia acaea* 24). *Hypothyris anastasia aureata* 25). *Hypothyris mansuetus meterus* 26). *Hyposcada anchiala mendax* 27). *Melinaea mothone* 28). *Mechanitis messenoides deceptus* 29). *Hypothyris anastasia bicolora* 30). *Melinaea isocomma simulator* 31). *Chetone histrio hydra* 32). *Hypothyris euclea callanga* 33). *Hypothyris fluonia seminigra* 34). *Forbestra olivencia olivencia* 35). *Melinaea menophilus orestes* 36). *Melinaea mnasias romualdo* 37). *Tithorea harmonia brunnea 38). Napeogenes zurippa deucalion* 39). *Heliconius pardalinus butleri* 40). *Melinaea marseus phasiana* 41). *Hypothyris anastasia anastasina* 42). *Melinaea menophilus hicetas* 43). *Mechanitis mazeus mazeus* 44). *Chetone histrio histrio* 45). *Hypothyris anastasia acreana* 46). *Melinaea satevis lamasi* 47). *Athyrtis mechanitis salvini 48). Melinaea marseus clara*. Some butterfly images courtesy of https://www.butterfliesofamerica.com/ (Andrew Warren); http://www.sangay.eu/esdex.php/ (Jean-Claude Petit).

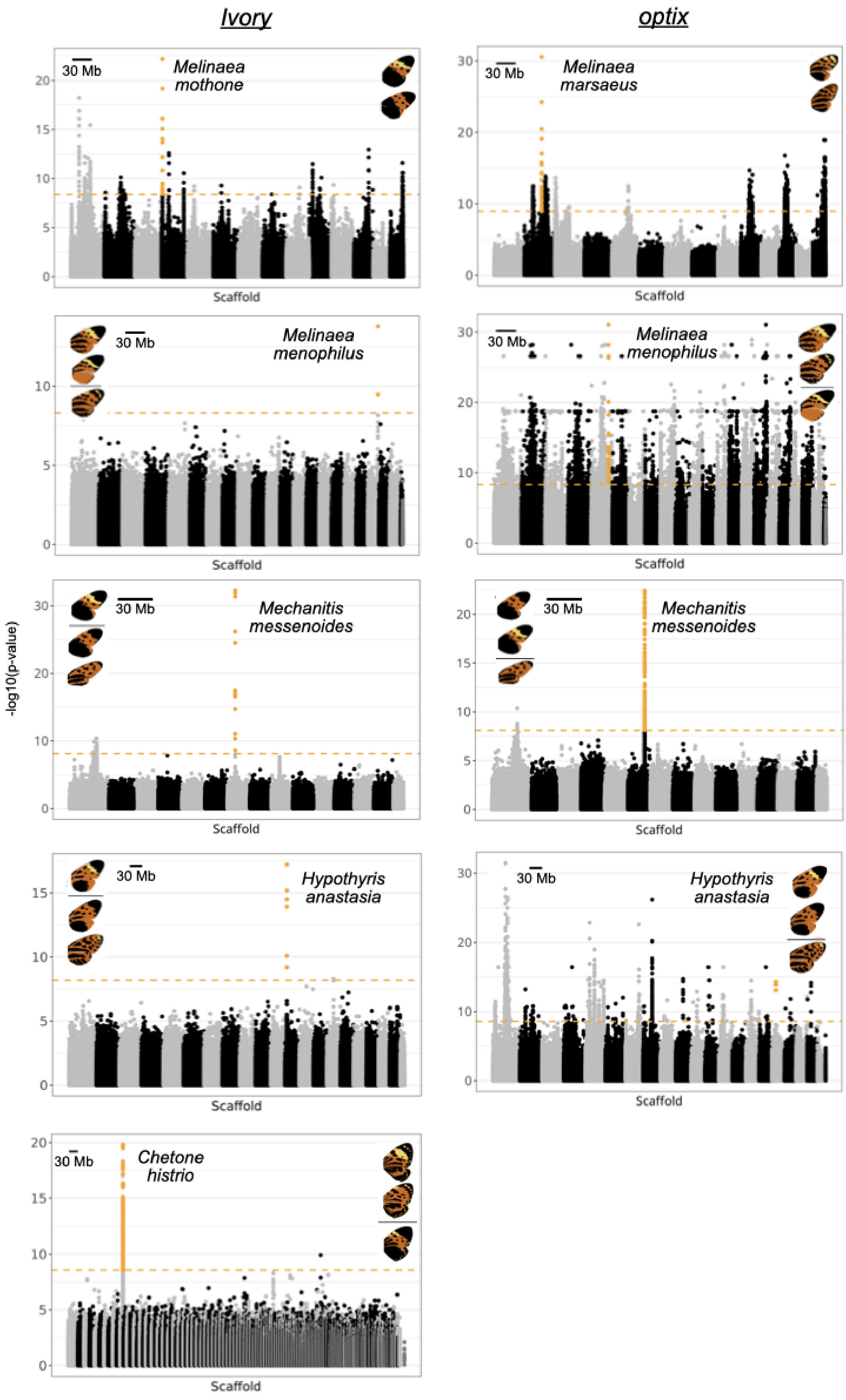

Ivory
optix
30 Mb
Melinaea mothone
Melinaea marsaeus
Melinaea menophilus
Mechanitis messenoides
Hypothyris anastasia
Chetone histrio
-log10(p-value)
Scaffold

**Extended Data Fig. 2 | Genome-wide associations for forewing yellow bar and hindwing orange/black wing phenotypes.** SNPs above the Bonferroni-corrected significance threshold (horizontal dashed line) in the main peak of association are highlighted in orange. The wing images on the right denote the phenotype compared in each analysis. Zoomed in plots of the peaks are shown in Figure 2. Additional associated SNPs in some of the species are likely a result of population structure correlated with the phenotypes (Extended Data Figs. 17-20).

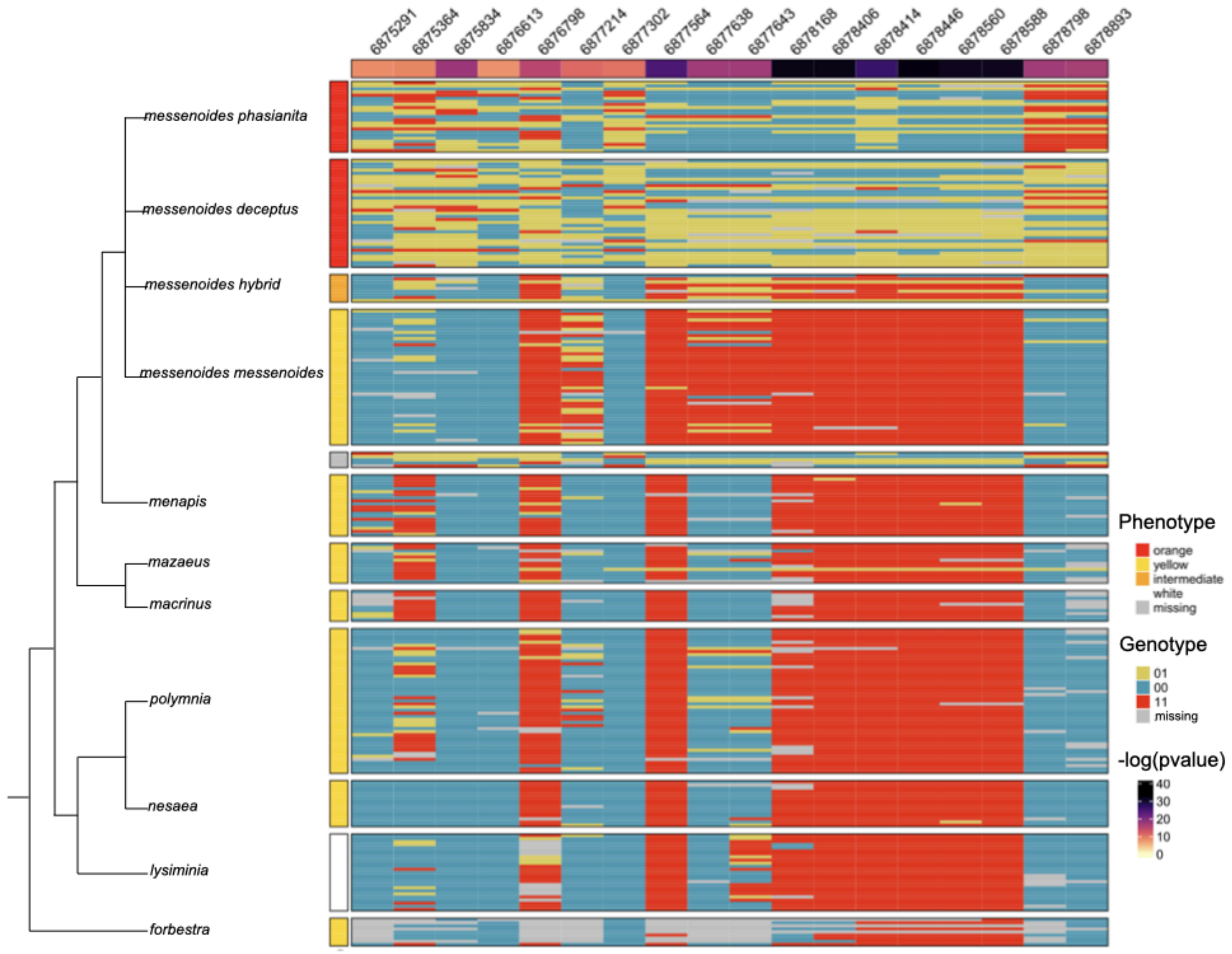


**Extended Data Fig. 3 | Genotypic variation at the *Mechanitis messenoides ivory* GWA peak in *Mechanitis* species.** This genotype matrix shows the genotypic states at the top SNPs from the GWA at the *ivory* locus, with individuals grouped by phylogenetic relationships (left dendrogram). Each row corresponds to an individual and each column to a SNP. The top panels (down to *messenoides messenoides*) include the focal species used in the GWAS. The remaining taxa are shown to illustrate the lack of association between genotype and phenotype at these SNPs across a broader phylogenetic context. Phenotype information (wing colour patterns) is represented by coloured boxes to the left of the genotype matrix. The numbers displayed along the top of the figure are the genomic positions of the SNPs. The GWA -log(p-value) values are displayed beneath the genomic positions, with colours ranging from yellow (low significance) to dark purple (high significance).

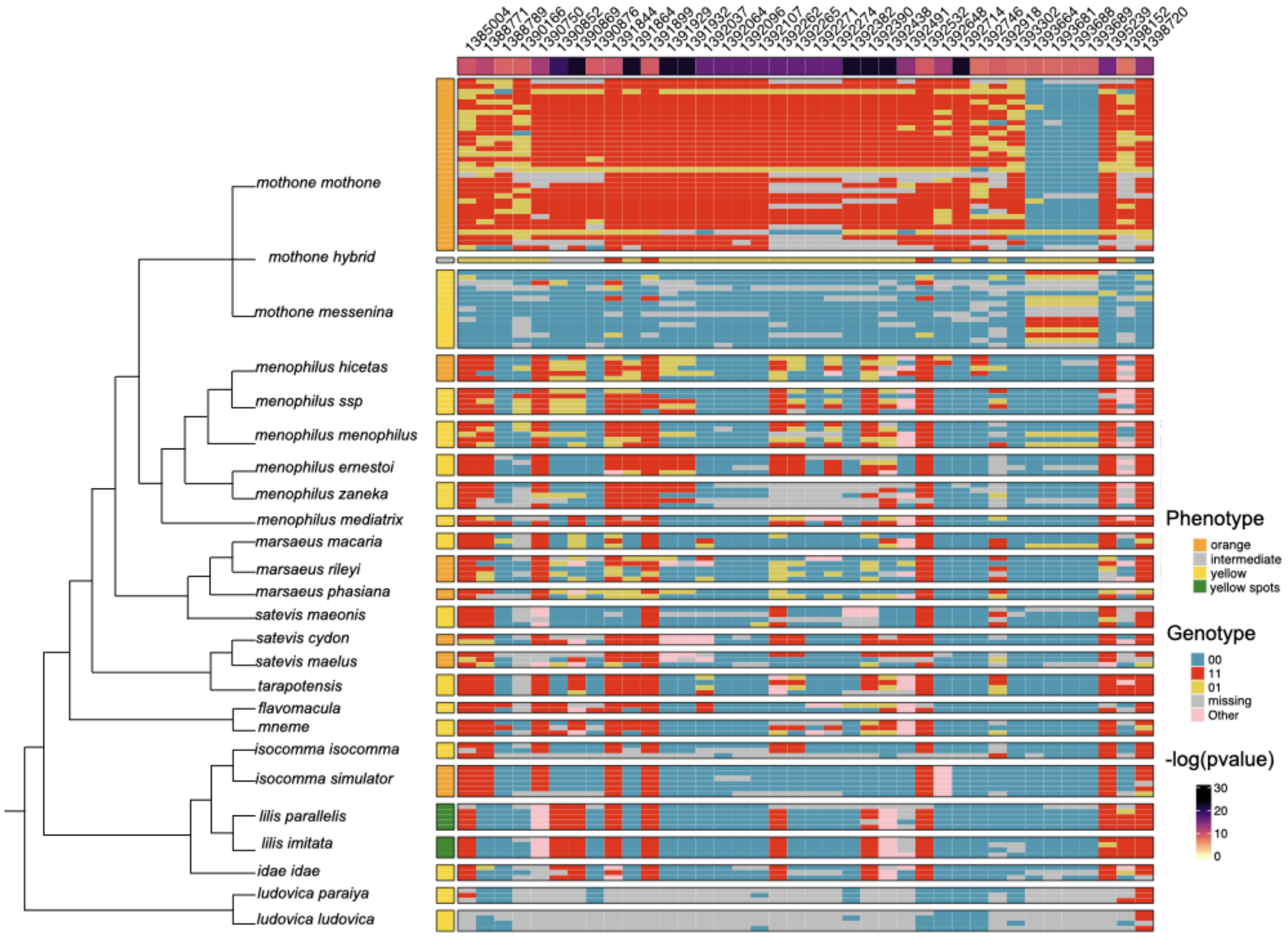


**Extended Data Fig. 4 | Genotypic variation at the *Melinaea mothone ivory* GWA peak in *Melinaea* species.** This genotype matrix shows the genotypic states at the top SNPs from the GWA at the *ivory* locus, with individuals grouped by phylogenetic relationships (left dendrogram). Each row corresponds to an individual and each column to a SNP. The top panels (down to *mothone messenina*) include the focal species used in the GWAS. The remaining taxa are shown to illustrate the lack of association between genotype and phenotype at these SNPs across a broader phylogenetic context. Phenotype information (wing colour patterns) is represented by coloured boxes to the left of the genotype matrix. The numbers displayed along the top of the figure are the genomic positions of the SNPs. The GWA -log(p-value) values are displayed beneath the genomic positions, with colours ranging from yellow (low significance) to dark purple (high significance).

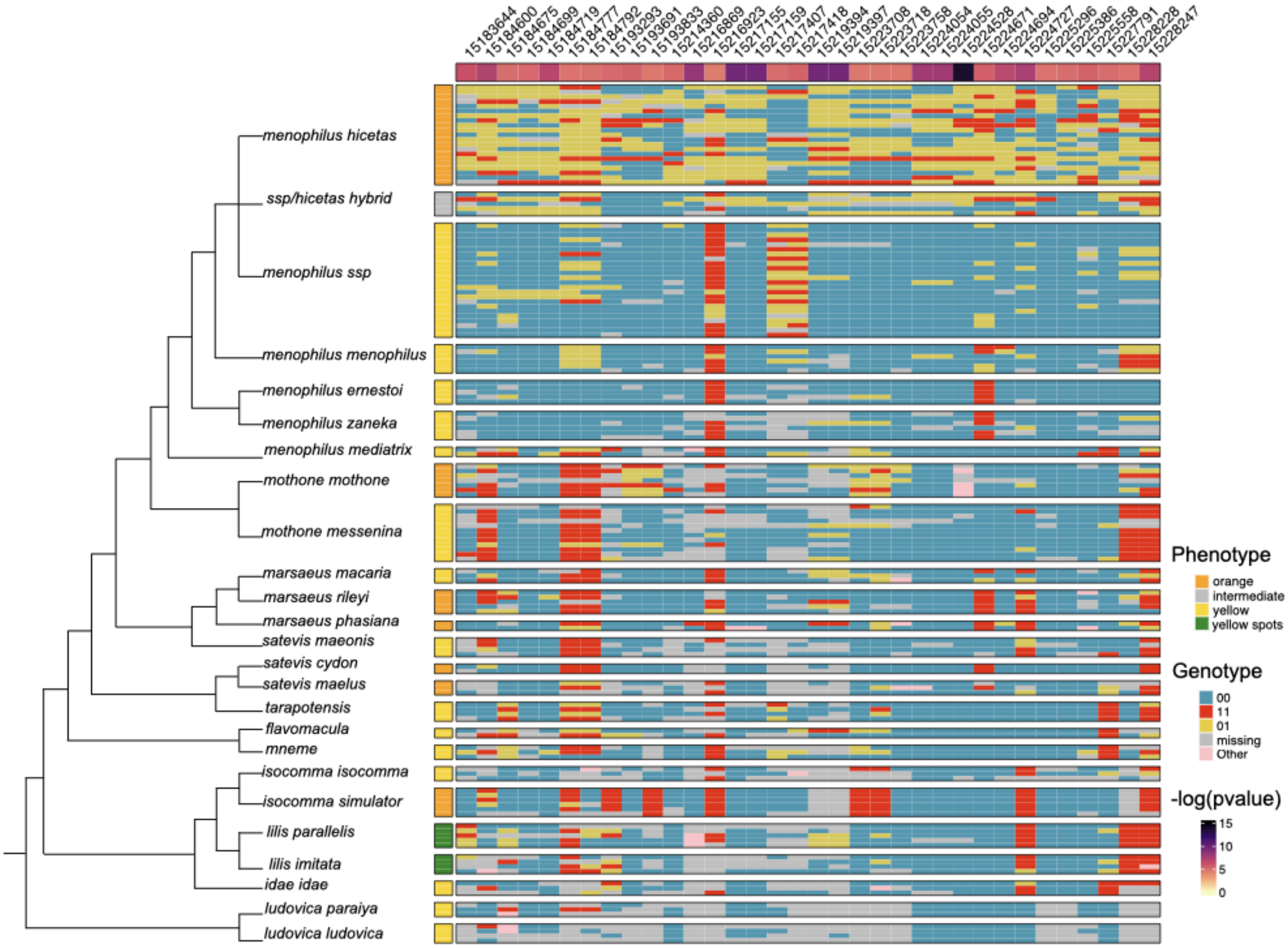


**Extended Data Fig. 5 | Genotypic variation at the *Melinaea menophilus ivory* GWA peak in *Melinaea* species.** This genotype matrix shows the genotypic states at the top SNPs from the GWA at the *ivory* locus, with individuals grouped by phylogenetic relationships (left dendrogram). Each row corresponds to an individual and each column to a SNP. The top panels (down to *menophilus zaneka*) include the focal species used in the GWAS. The remaining taxa are shown to illustrate the lack of association between genotype and phenotype at these SNPs across a broader phylogenetic context. Phenotype information (wing colour patterns) is represented by coloured boxes to the left of the genotype matrix. The numbers displayed along the top of the figure are the genomic positions of the SNPs. The GWA -log(p-value) values are displayed beneath the genomic positions, with colours ranging from yellow (low significance) to dark purple (high significance).

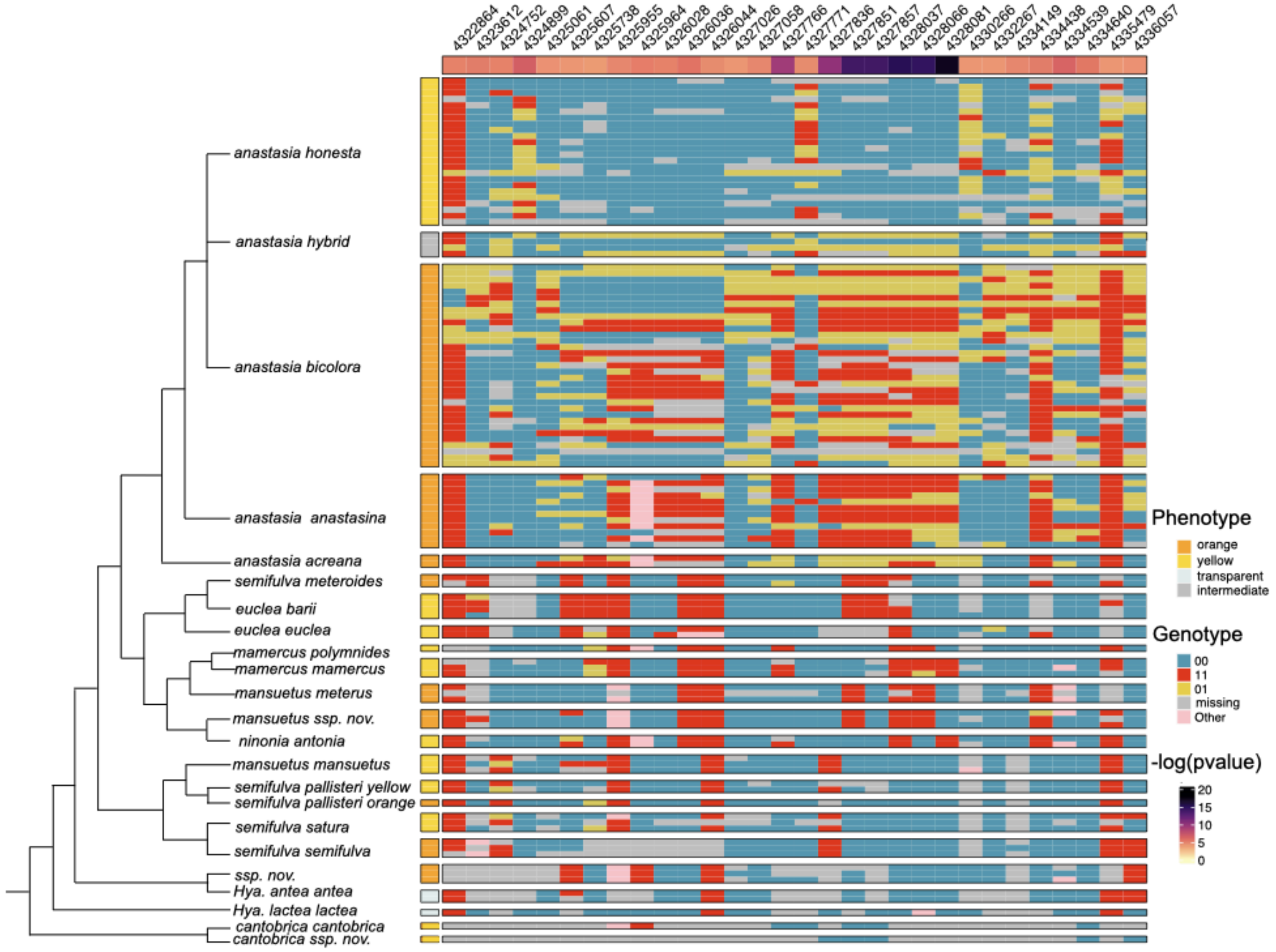


**Extended Data Fig. 6 | Genotypic variation at the *Hypothyris anastasia ivory* GWA peak in *Hypothyris* species.** This genotype matrix shows the genotypic states at the top SNPs from the GWA at the *ivory* locus, with individuals grouped by phylogenetic relationships (left dendrogram). Each row corresponds to an individual and each column to a SNP. The top panels (down to *anastasia bicolora*) include the focal species used in the GWAS. The remaining taxa are shown to illustrate the lack of association between genotype and phenotype at these SNPs across a broader phylogenetic context. Phenotype information (wing colour patterns) is represented by coloured boxes to the left of the genotype matrix. The numbers displayed along the top of the figure are the genomic positions of the SNPs. The GWA -log(p-value) values are displayed beneath the genomic positions, with colours ranging from yellow (low significance) to dark purple (high significance).

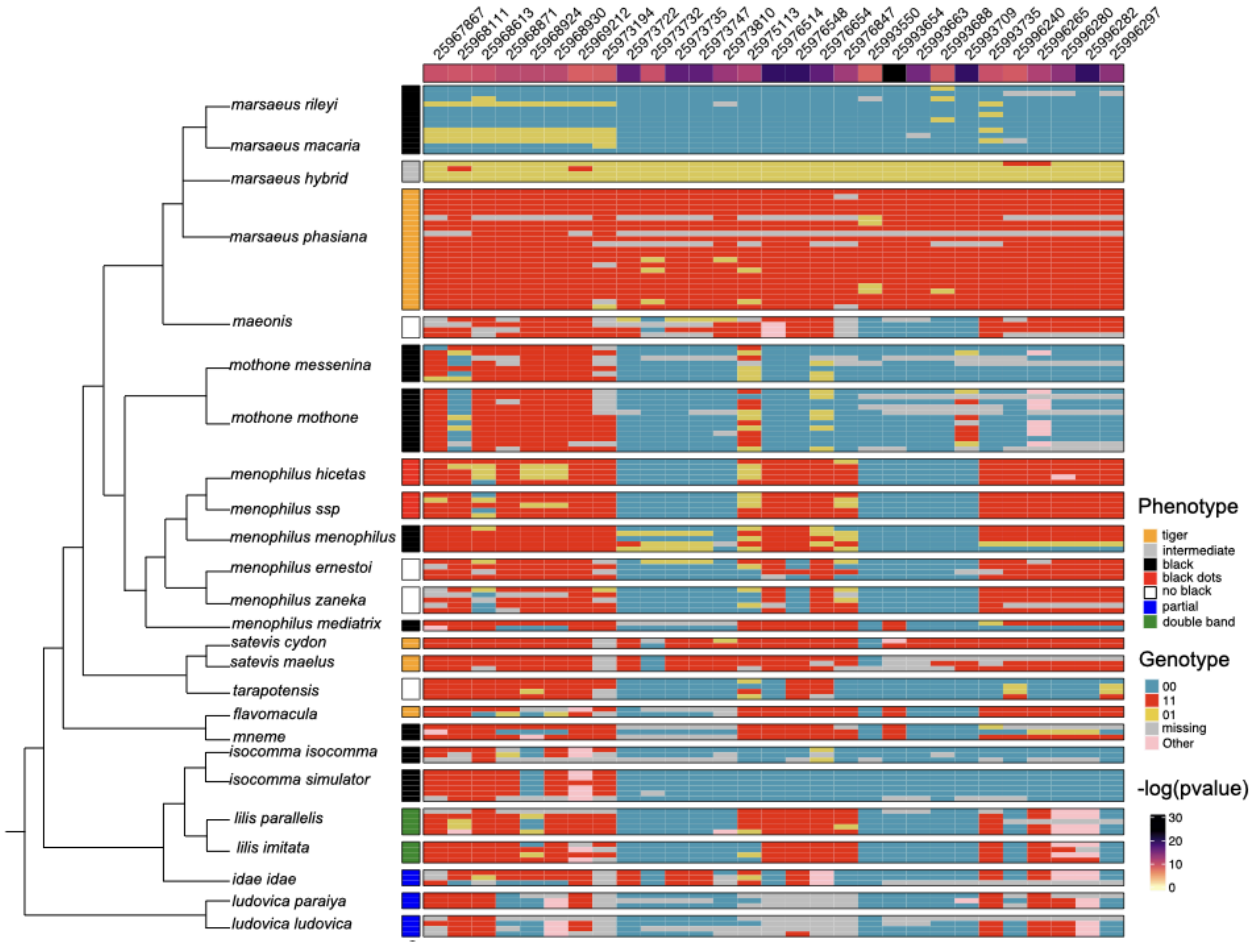


**Extended Data Fig. 7 | Genotypic variation at the *Melinaea marsaeus optix* GWA peak in *Melinaea* species.** This genotype matrix shows the genotypic states at the top SNPs from the GWA at the *optix* locus, with individuals grouped by phylogenetic relationships (left dendrogram). Each row corresponds to an individual and each column to a SNP. The top panels (down to *marsaeus phasiana*) include the focal species used in the GWAS. The remaining taxa are shown to illustrate the association between genotype and phenotype at these SNPs across a broader phylogenetic context. Phenotype information (wing colour patterns) is represented by coloured boxes to the left of the genotype matrix. The numbers displayed along the top of the figure are the genomic positions of the SNPs. The GWA -log(p-value) values are displayed beneath the genomic positions, with colours ranging from yellow (low significance) to dark purple (high significance).

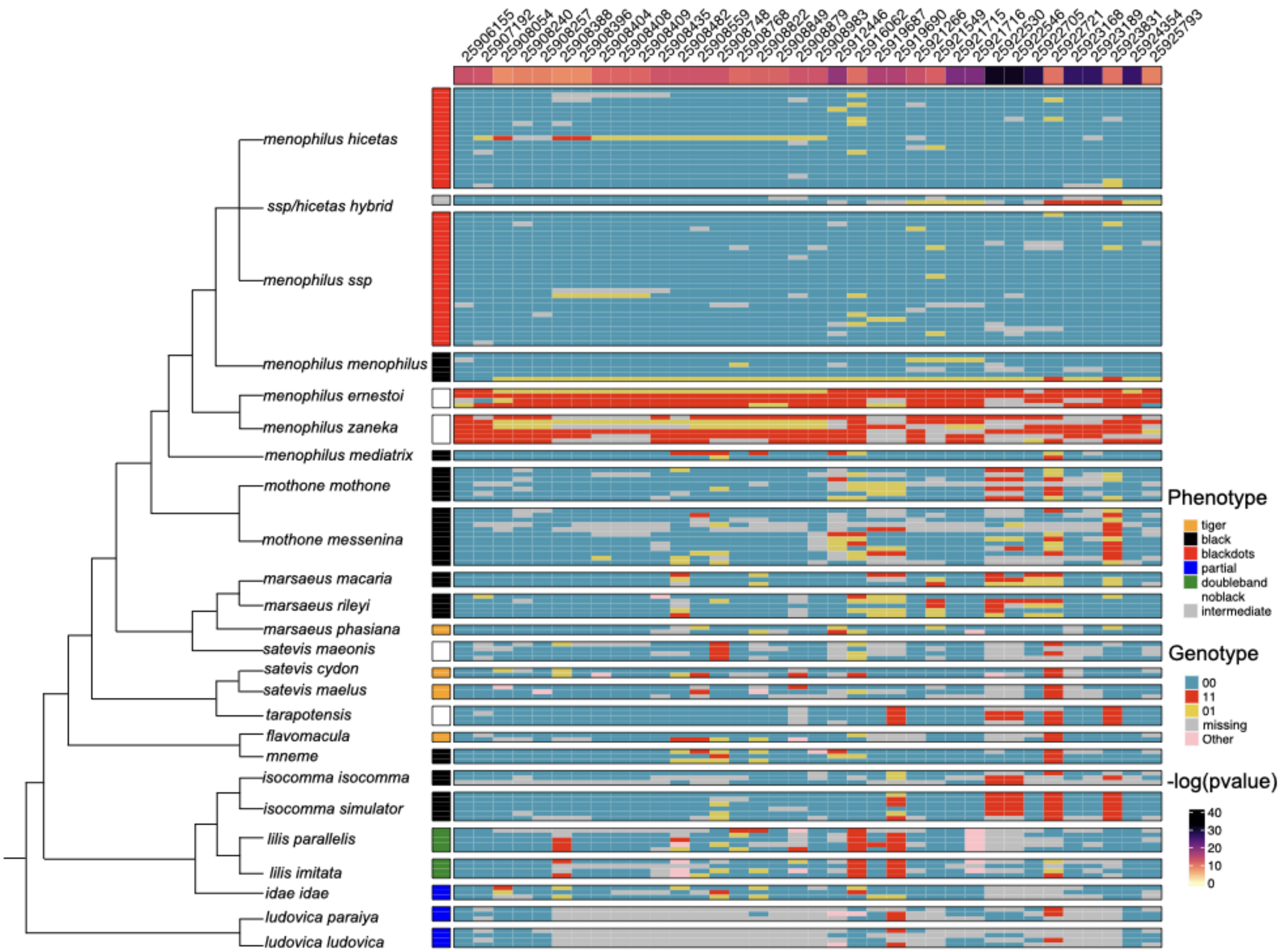


**Extended Data Fig. 8 | Genotypic variation at the *Melinaea menophilus optix* GWA peak in *Melinaea* species.** This genotype matrix shows the genotypic states at the top SNPs from the GWA at the *optix* locus, with individuals grouped by phylogenetic relationships (left dendrogram). Each row corresponds to an individual and each column to a SNP. The top panels (down to *menophilus zaneka*) include the focal species used in the GWAS. The remaining taxa are shown to illustrate the lack of association between genotype and phenotype at these SNPs across a broader phylogenetic context. Phenotype information (wing colour patterns) is represented by coloured boxes to the left of the genotype matrix. The numbers displayed along the top of the figure are the genomic positions of the SNPs. The GWA -log(p-value) values are displayed beneath the genomic positions, with colours ranging from yellow (low significance) to dark purple (high significance).

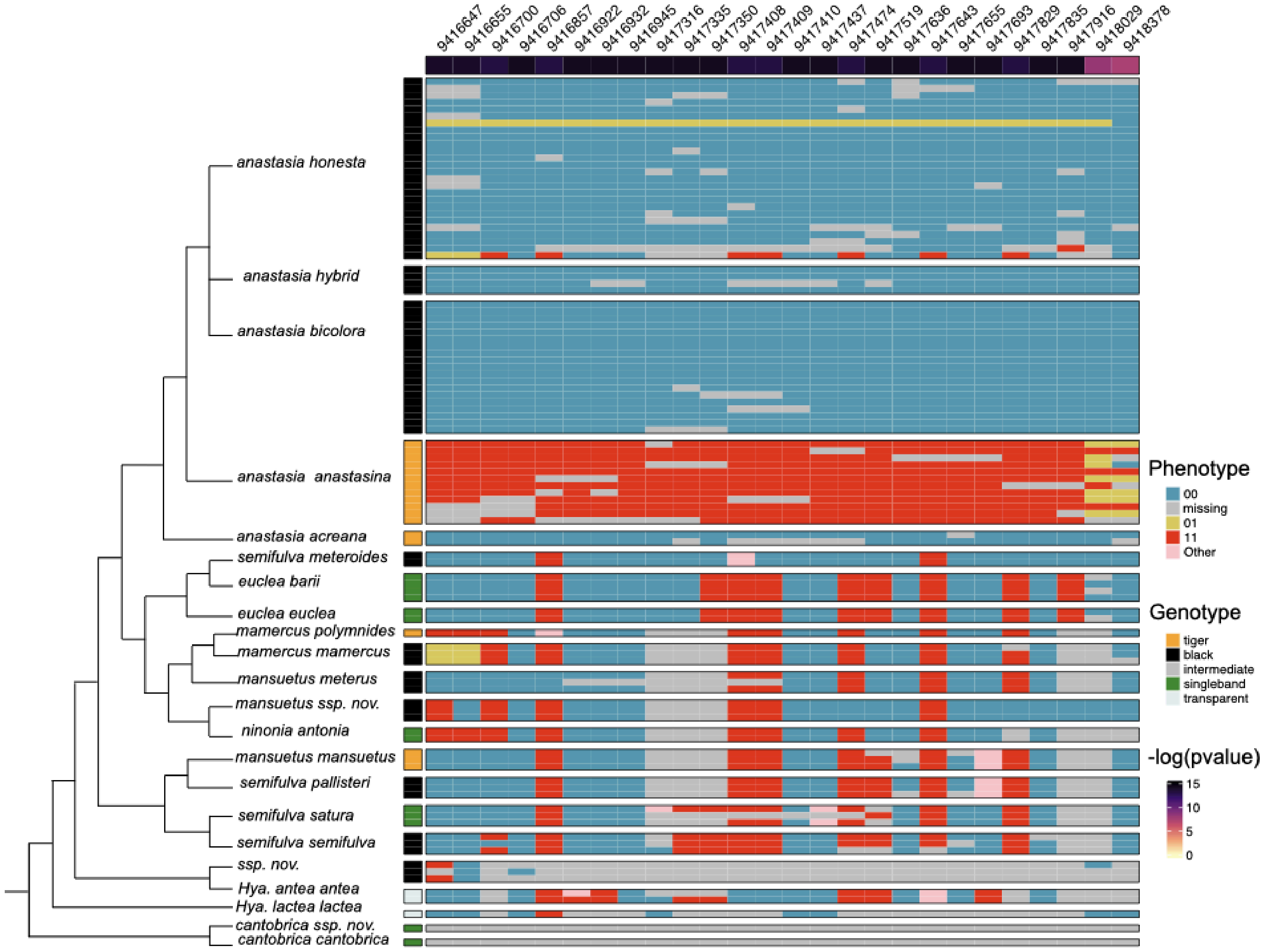


**Extended Data Fig. 9 | Genotypic variation at the *Hypothyris anastasia optix* GWA peak in *Hypothyris* species.** This genotype matrix shows the genotypic states at the top SNPs from the GWA at the *optix* locus, with individuals grouped by phylogenetic relationships (left dendrogram). Each row corresponds to an individual and each column to a SNP. The top panels (down to *anastasia acreana*) include the focal species used in the GWAS. The remaining taxa are shown to illustrate the lack of association between genotype and phenotype at these SNPs across a broader phylogenetic context. Phenotype information (wing colour patterns) is represented by coloured boxes to the left of the genotype matrix. The numbers displayed along the top of the figure are the genomic positions of the SNPs. The GWA -log(p-value) values are displayed beneath the genomic positions, with colours ranging from yellow (low significance) to dark purple (high significance).

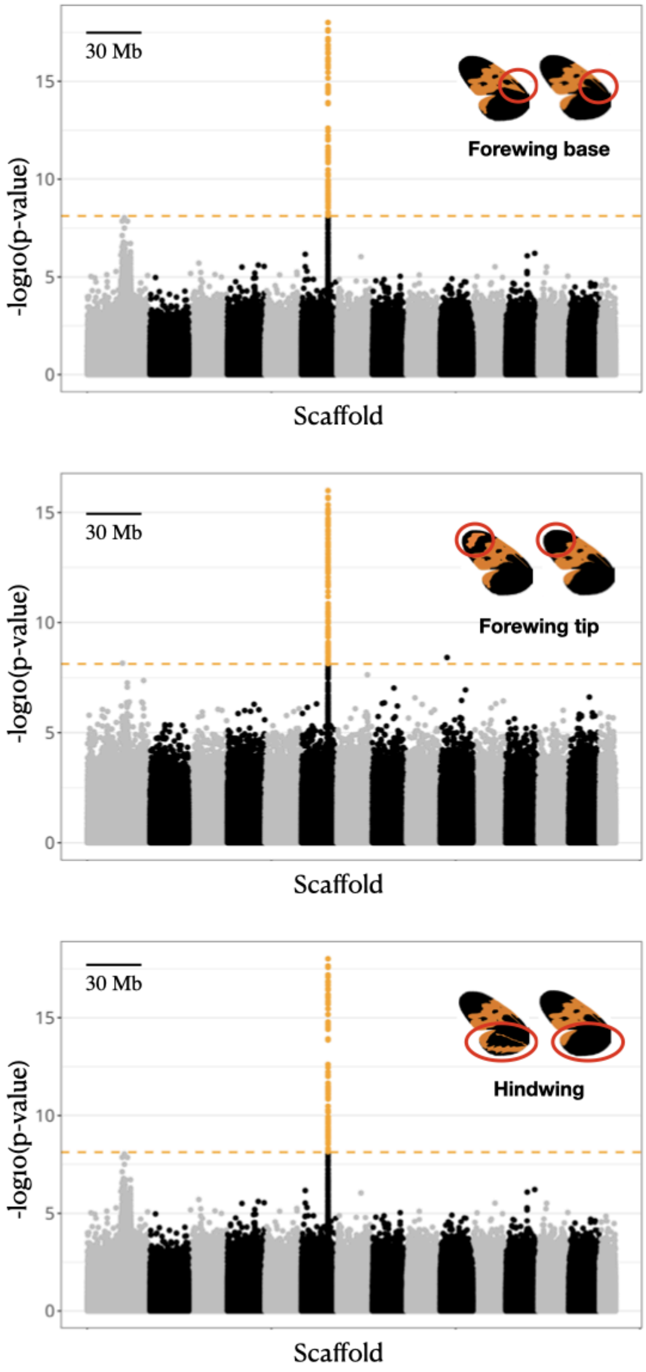


**Extended Data Fig. 10 | Genome-wide associations for different black-orange wing phenotypes in *Mechanitis messenoides*.** Genome-wide associations for black and orange wing patterns are shown for three different wing regions: forewing base, forewing tip, and hindwing melanization (wing images show the phenotype compared in each analysis). SNPs above the Bonferroni-corrected significance threshold (horizontal dashed line) in the main peak of association are highlighted in orange, which in all cases lies near *optix*. A zoomed in plot of the peak is shown in Figure 4a.

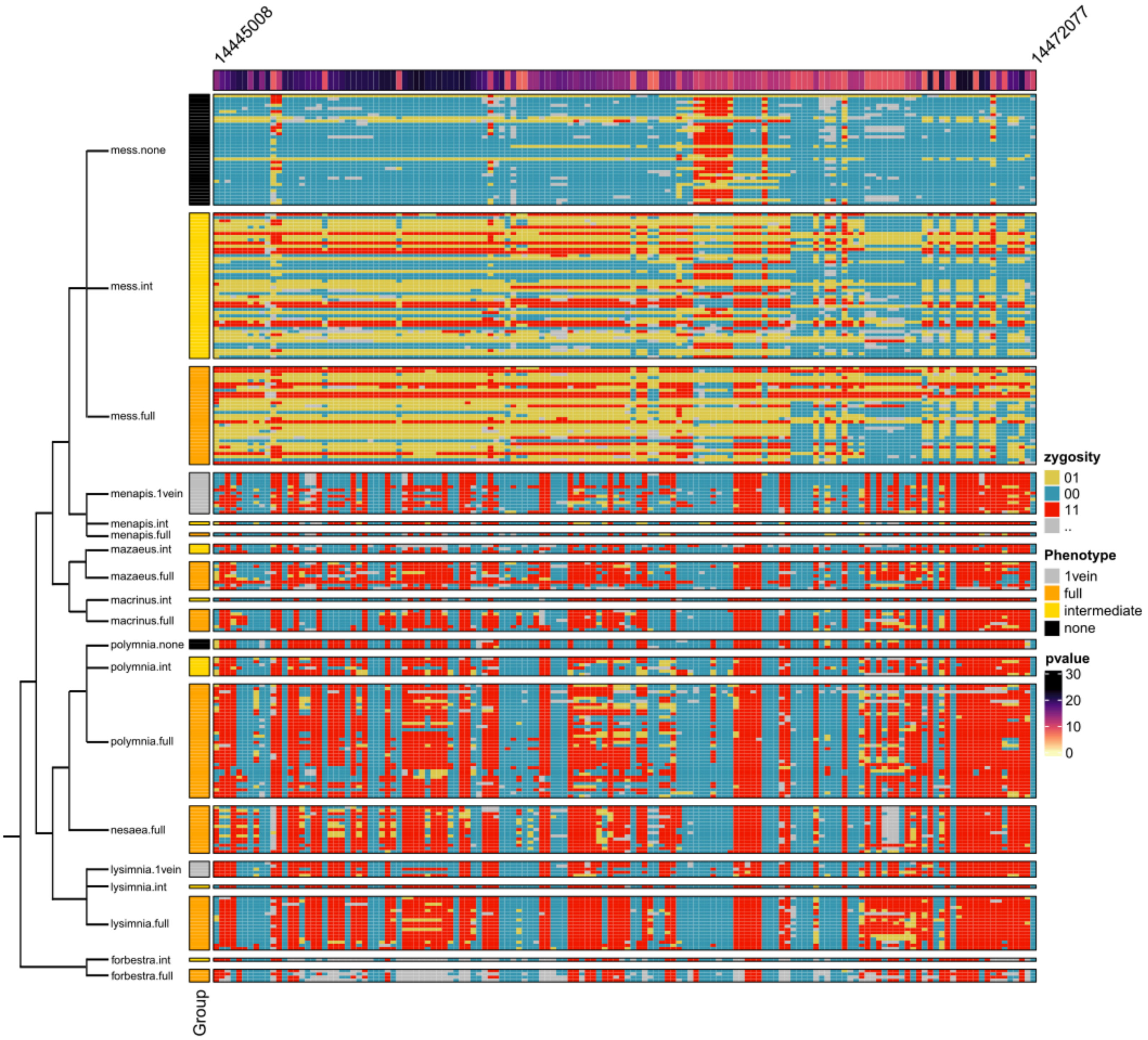


**Extended Data Fig. 11 | Genotypic variation at the *Mechanitis messenoides optix* forewing base GWA peak in *Mechanitis* species.** This genotype matrix shows the genotypic states at the top SNPs from the GWA at the *optix* locus, with individuals grouped by phylogenetic relationships (left dendrogram). Each row corresponds to an individual and each column to a SNP. The top three panels include the focal species used in the GWAS. The remaining taxa are shown to illustrate the lack of association between genotype and phenotype at these SNPs across a broader phylogenetic context. Phenotype information (wing colour patterns) is represented by coloured boxes to the left of the genotype matrix. The numbers displayed along the top of the figure are the genomic positions of the SNPs. The GWA -log(p-value) values are displayed beneath the genomic positions, with colours ranging from yellow (low significance) to dark purple (high significance).

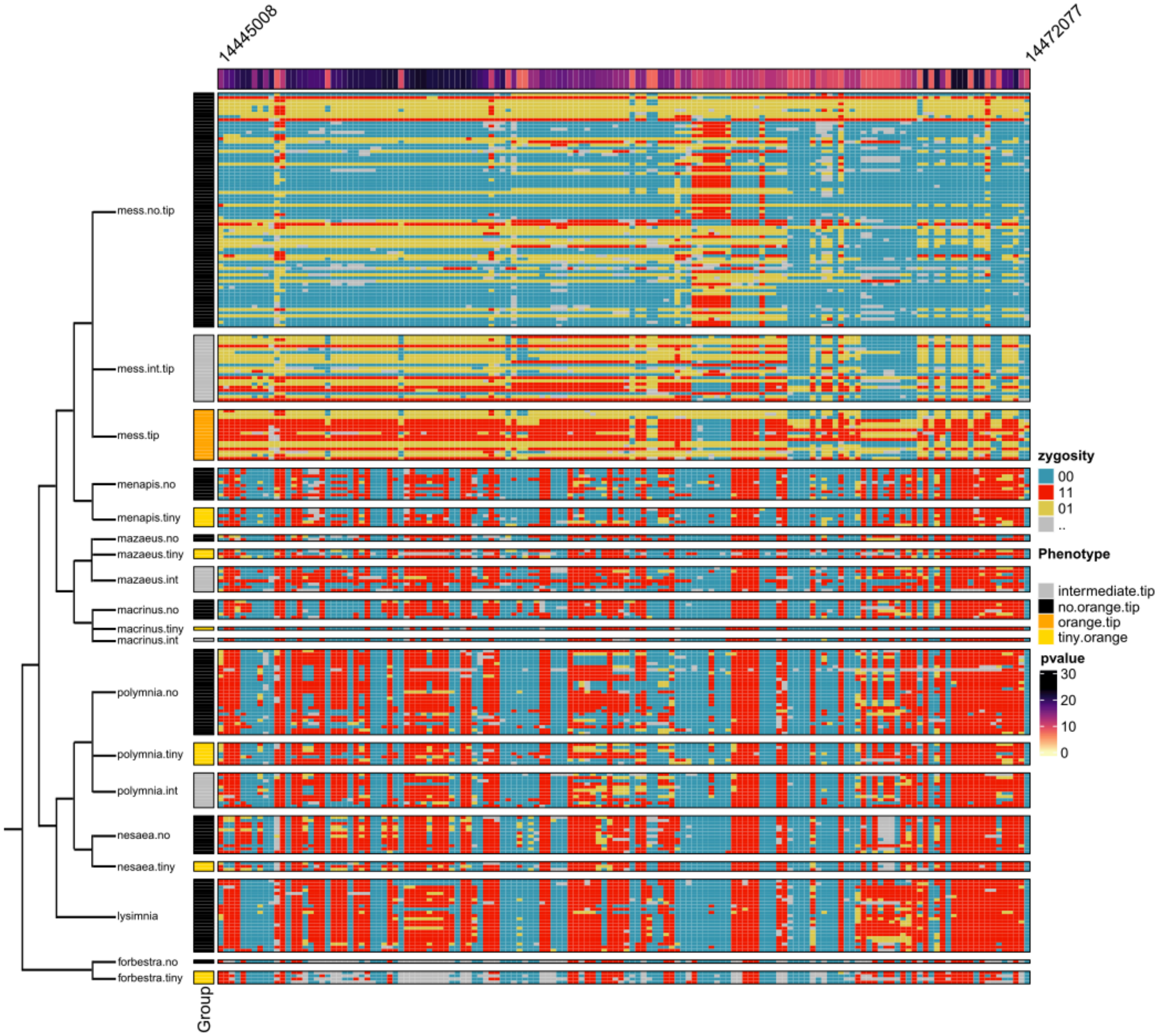


**Extended Data Fig. 12 | Genotypic variation at the *Mechanitis messenoides optix* forewing tip GWA peak in *Mechanitis* species.** This genotype matrix shows the genotypic states at the top SNPs from the GWA at the *optix* locus, with individuals grouped by phylogenetic relationships (left dendrogram). Each row corresponds to an individual and each column to a SNP. The top three panels include the focal species used in the GWAS. The remaining taxa are shown to illustrate the lack of association between genotype and phenotype at these SNPs across a broader phylogenetic context. Phenotype information (wing colour patterns) is represented by coloured boxes to the left of the genotype matrix. The numbers displayed along the top of the figure are the genomic positions of the SNPs. The GWA -log(p-value) values are displayed beneath the genomic positions, with colours ranging from yellow (low significance) to dark purple (high significance).

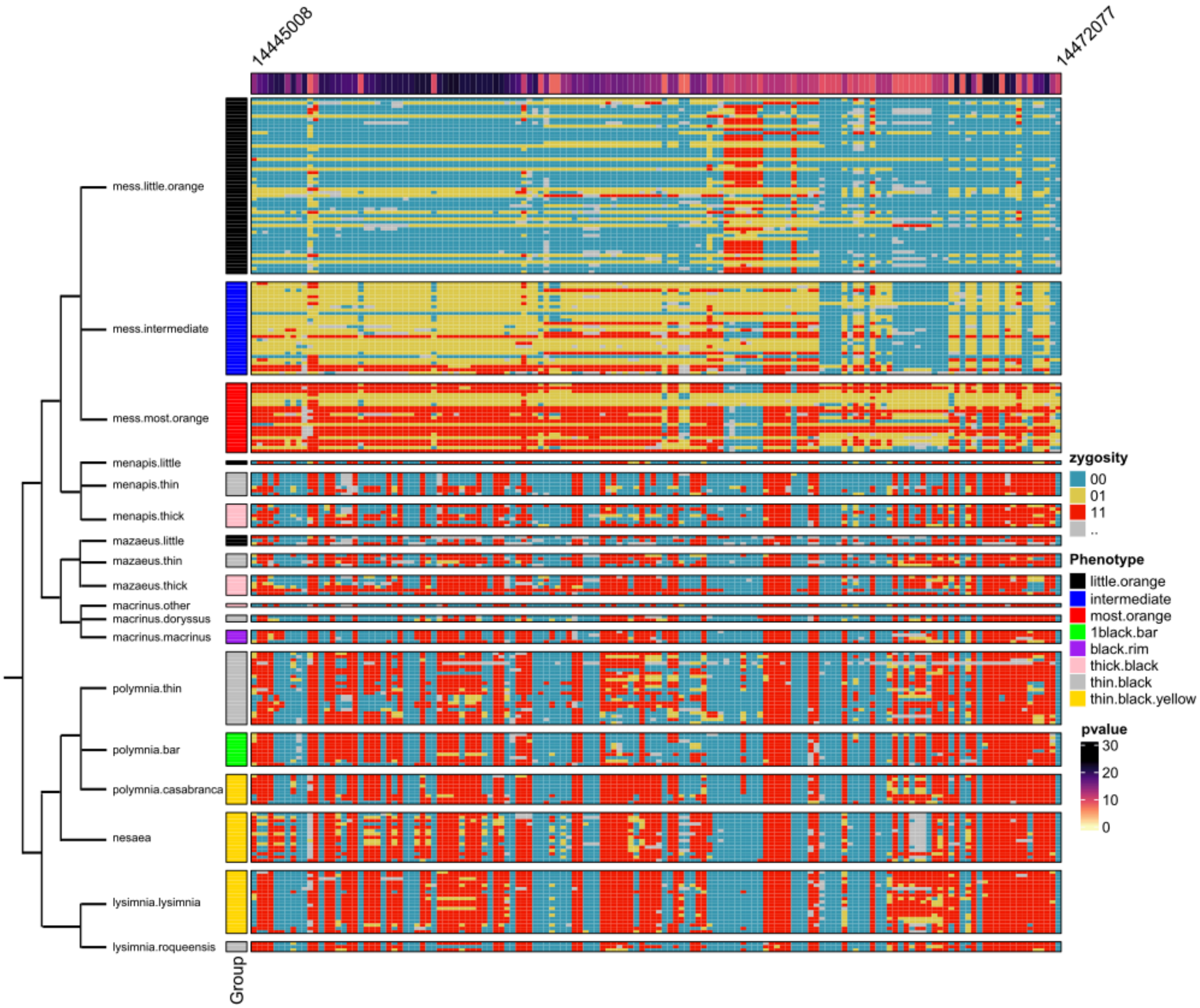


**Extended Data Fig. 13 | Genotypic variation at the *Mechanitis messenoides optix* hindwing GWA peak in *Mechanitis* species.** This genotype matrix shows the genotypic states at the top SNPs from the GWA at the *optix* locus, with individuals grouped by phylogenetic relationships (left dendrogram). Each row corresponds to an individual and each column to a SNP. The top three panels include the focal species used in the GWAS. The remaining taxa are shown to illustrate the lack of association between genotype and phenotype at these SNPs across a broader phylogenetic context. Phenotype information (wing colour patterns) is represented by coloured boxes to the left of the genotype matrix. The numbers displayed along the top of the figure are the genomic positions of the SNPs. The GWA -log(p-value) values are displayed beneath the genomic positions, with colours ranging from yellow (low significance) to dark purple (high significance).

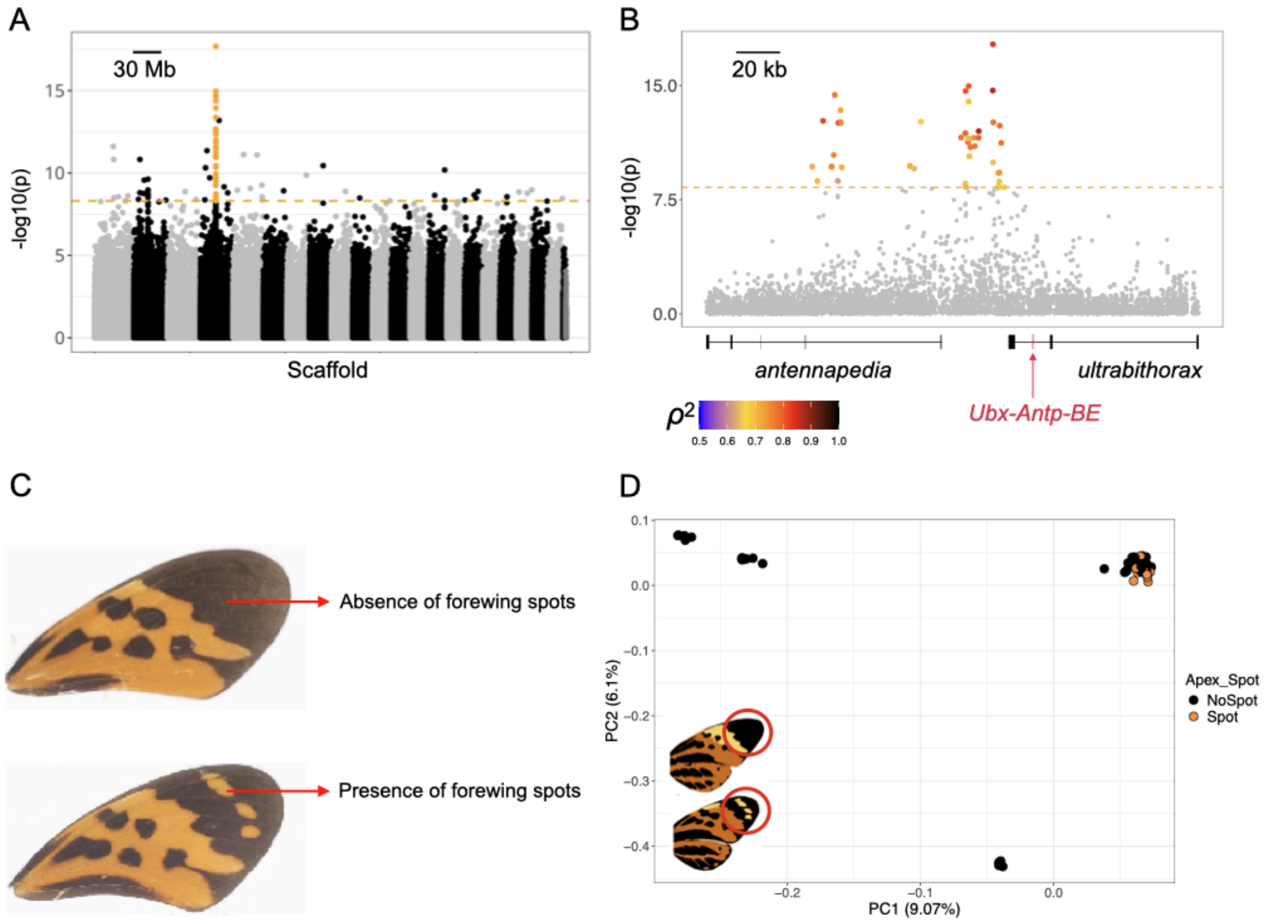


**Extended Data Fig. 14 | Genome wide association for the forewing apical spot in *Melinaea menophilus*. a,** Manhattan plot for the whole genome. The orange dashed line represents the threshold of significance. SNPs in the main peak of association are highlighted in orange. **b,** Zoom on the main peak of association (SUPER_4:16233326-16458737) with locations of annotated genes. SNPs above the Bonferroni-corrected significance threshold (dashed orange line) are coloured according to the squared Spearman's rank correlation coefficient ($\rho^2$), which indicates the strength of association between genotype and phenotype. Red arrow: position of Topologically Associated Domain boundary element *Antp-Ubx_BE* (Tendolkar et al. 2024). **c,** The wing phenotypes compared in the analysis. **d,** Principal component analysis of 412,584 LD pruned biallelic SNPs showing the population structure among the sampled individuals. Points are coloured by apex spot phenotype.

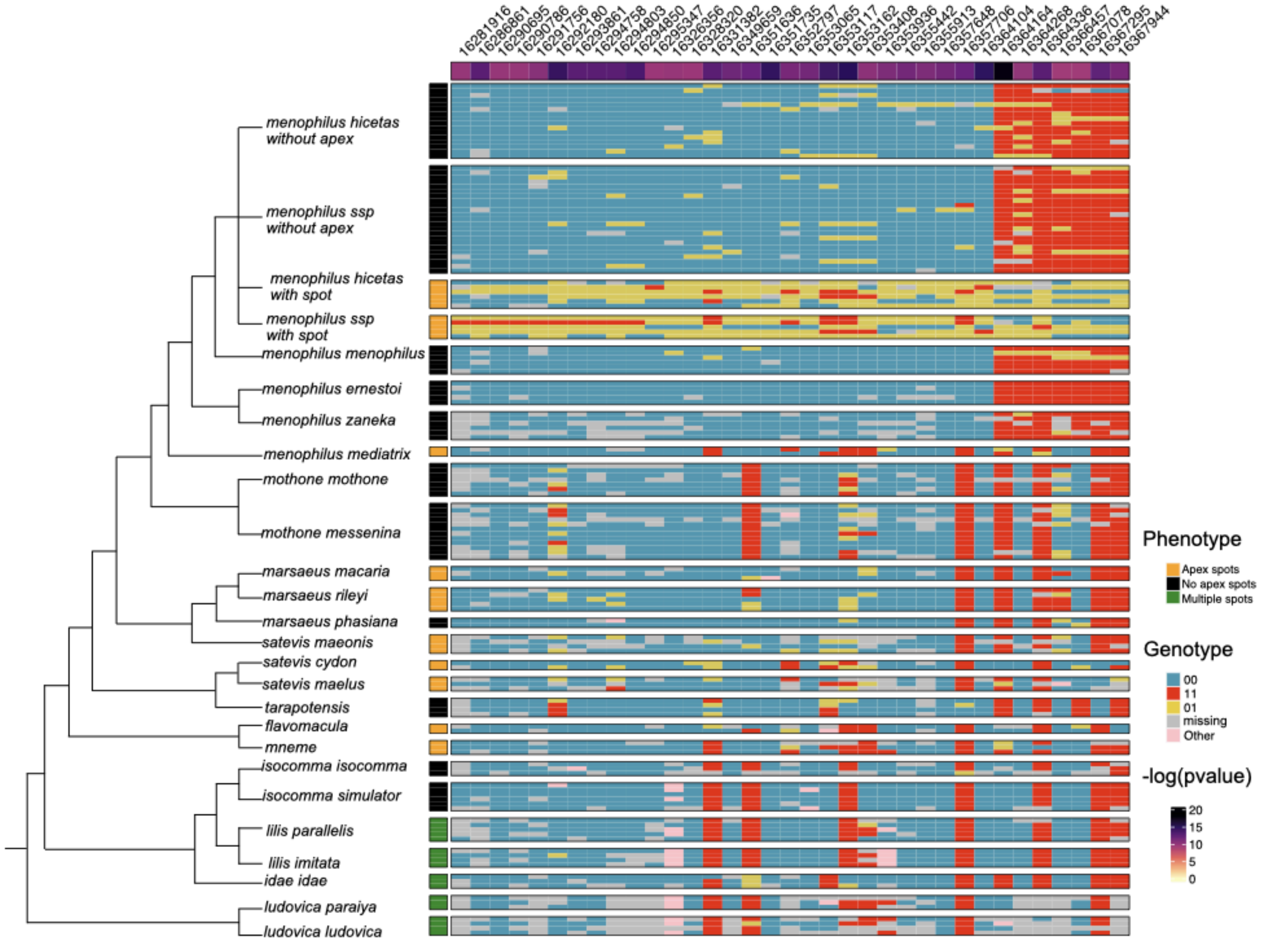


**Extended Data Fig. 15 | Genotypic variation at the *Melinaea menophilus antennapedia* GWA peak in *Melinaea* species.** This genotype matrix shows the genotypic states at the top SNPs from the GWA at the *antennapedia* locus, with individuals grouped by phylogenetic relationships (left dendrogram). Each row corresponds to an individual and each column to a SNP. The top panels (down to *menophilus zaneka*) include the focal species used in the GWAS. The remaining taxa are shown to illustrate the lack of association between genotype and phenotype at these SNPs across a broader phylogenetic context. Phenotype information (wing colour patterns) is represented by coloured boxes to the left of the genotype matrix. The numbers displayed along the top of the figure are the genomic positions of the SNPs. The GWA -log(p-value) values are displayed beneath the genomic positions, with colours ranging from yellow (low significance) to dark purple (high significance).

**a**

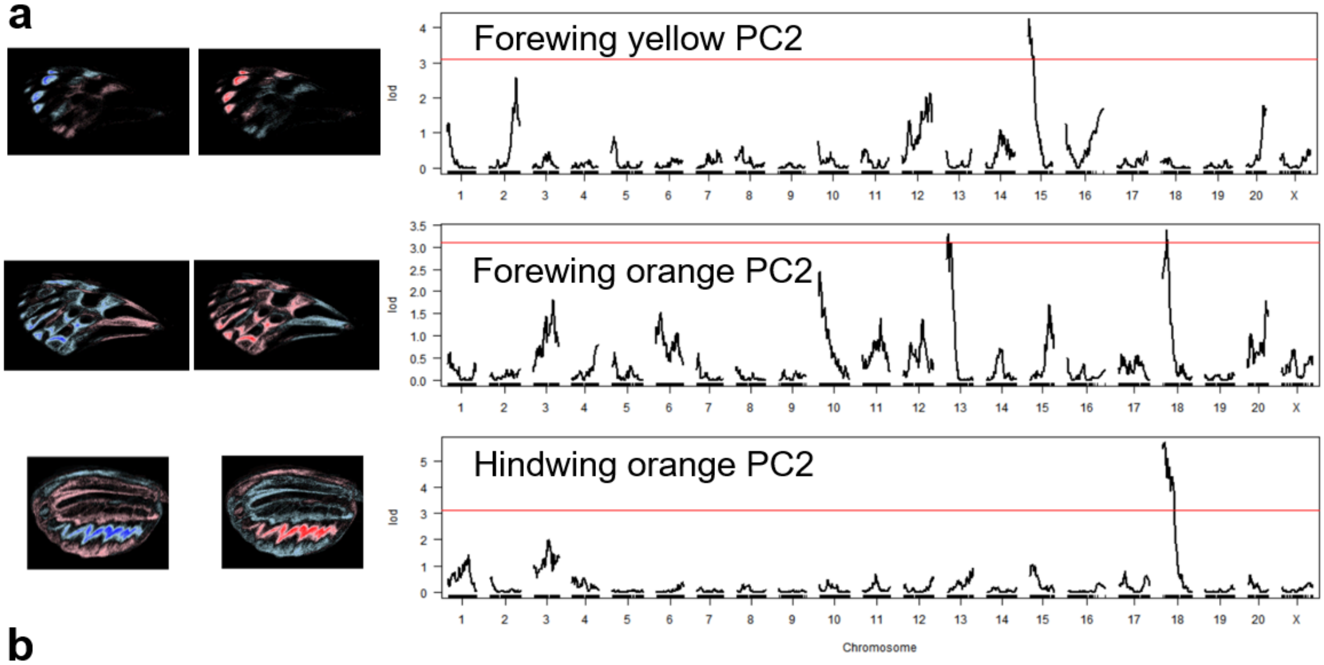


**b**

| Phenotype | LOD max | Chrom max | physical peak | Physical limits | $R^2$ |
|---|---|---|---|---|---|
| Forewing yellow PC2 | 4.27 | 15 | Hmel215001o:6078-Hmel215003o:344661 | Hmel215001o:2349-Hmel215003o:4230382 | 0.21 |
| Forewing orange PC2 | 3.39 | 18 | Hmel218003o:3001552-3078881 | Hmel218001o:7968-Hmel218003o:3843642 | 0.17 |
| Forewing orange PC2 | 3.3 | 13 | Hmel213001o:2293499-3149684 | Hmel213001o:21488-Hmel213001o:5242923 | 0.17 |
| Hindwing orange PC2 | 5.7 | 18 | Hmel218001o:8057-Hmel218003o:1845297 | Hmel218001o:7968-Hmel218003o:4324414 | 0.28 |

**Extended Data Fig. 16 | QTL mapping intervals for forewing and hindwing pattern variation in *Heliconius pardalinus*. a,** Mapping intervals for yellow and orange forewing and hindwing pattern variation. Heatmap images to the left indicate the phenotypes exhibited by the individuals at the extremes of each PC axis; red = presence of the colour, blue = absence of the colour. Note that the forewing has only yellow/white, orange/red and black/brown colours, and the hindwing only orange/red and black/brown. Since the black/brown pattern variation is a complement of the other colours, we do not show the results of the black/brown variation. **b,** Details of the QTL mapping intervals.

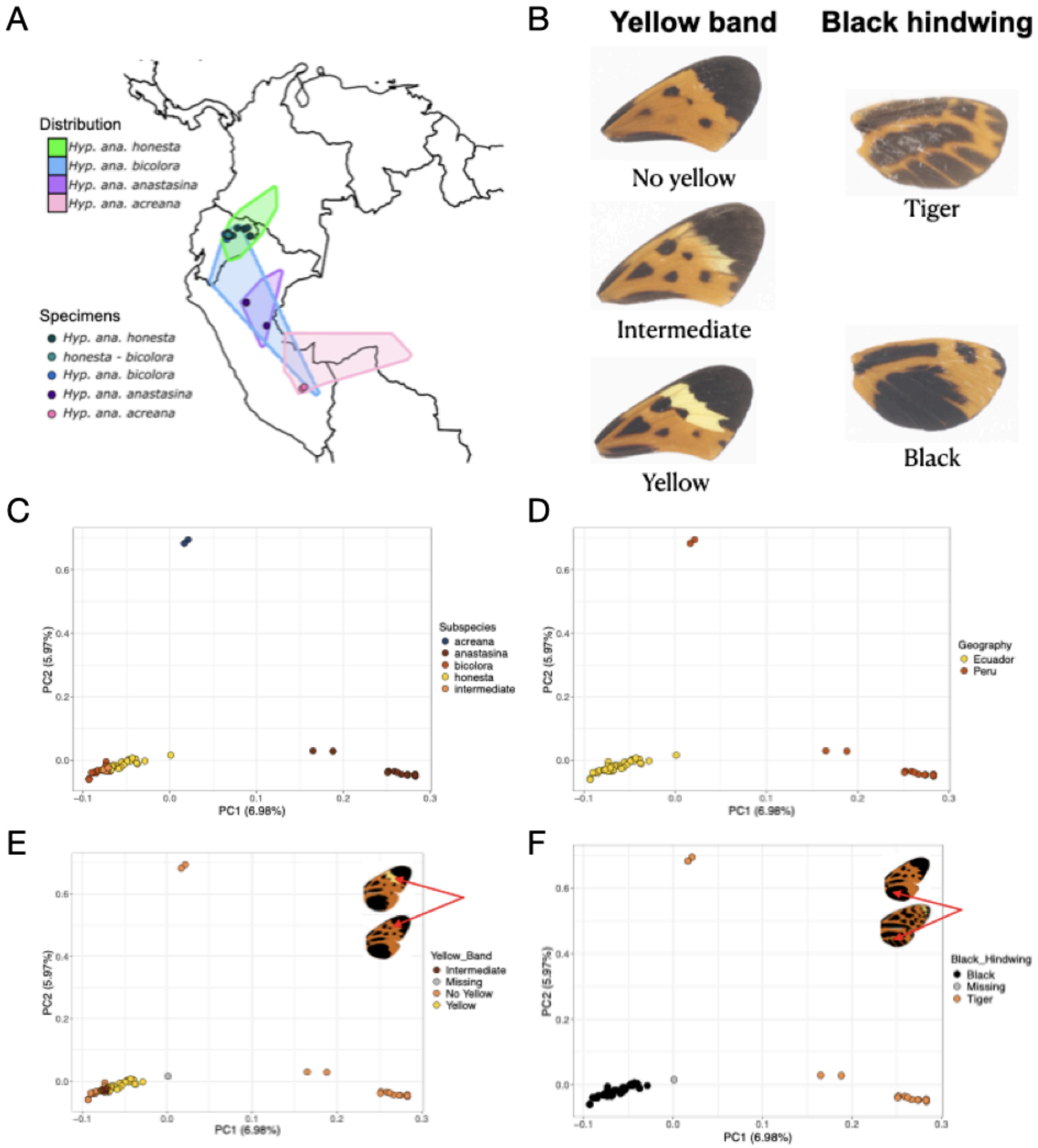


**Extended Data Fig. 17 | *Hypothyris anastasia*: taxon distribution, wing phenotypes and genetic PCA**. (A) Geographical distribution of *Hypothyris anastasia* subspecies, with specimen collection locations indicated. (B) Phenotypes used in the GWA. (C to F): Principal component analysis (PCA) showing the genetic distance between the sampled *Hypothyris anastasia* individuals using a dataset consisting of 400,452 LD pruned biallelic SNPS. The scatter plots correspond to the first two principal components (PCs). Points are coloured by subspecies (C), geography (D), yellow bar phenotype (E), and hindwing black phenotype (D). The wing images highlight the colour-coded phenotypes.

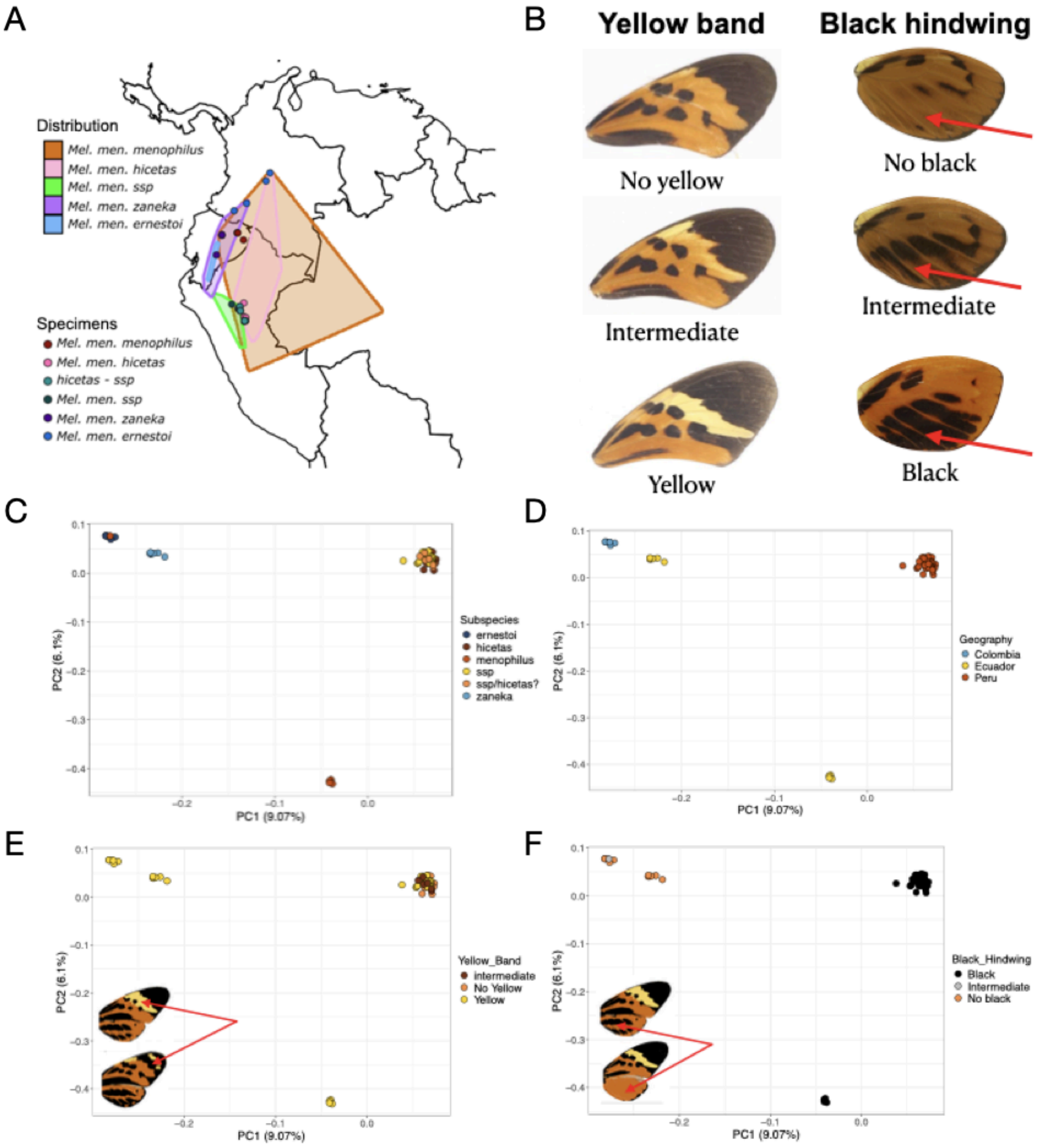


**Extended Data Fig. 18 | *Melinaea menophilus*: taxon distribution, wing phenotypes and genetic PCA*.*** (A) Geographical distribution of *Melinaea menophilus* subspecies, with specimen collection locations indicated. (B) Phenotypes used in the GWA. (C to F): Principal Principal component analysis (PCA) showing the genetic distance between the sampled *Melinaea menophilus* individuals using a dataset consisting of 412,584 LD pruned biallelic SNPS. The scatter plots correspond to the first two principal components (PCs). Points are coloured by subspecies (C), geography (D), yellow bar phenotype (E) and hindwing black phenotype (F). The wing images highlight the colour-coded phenotypes.

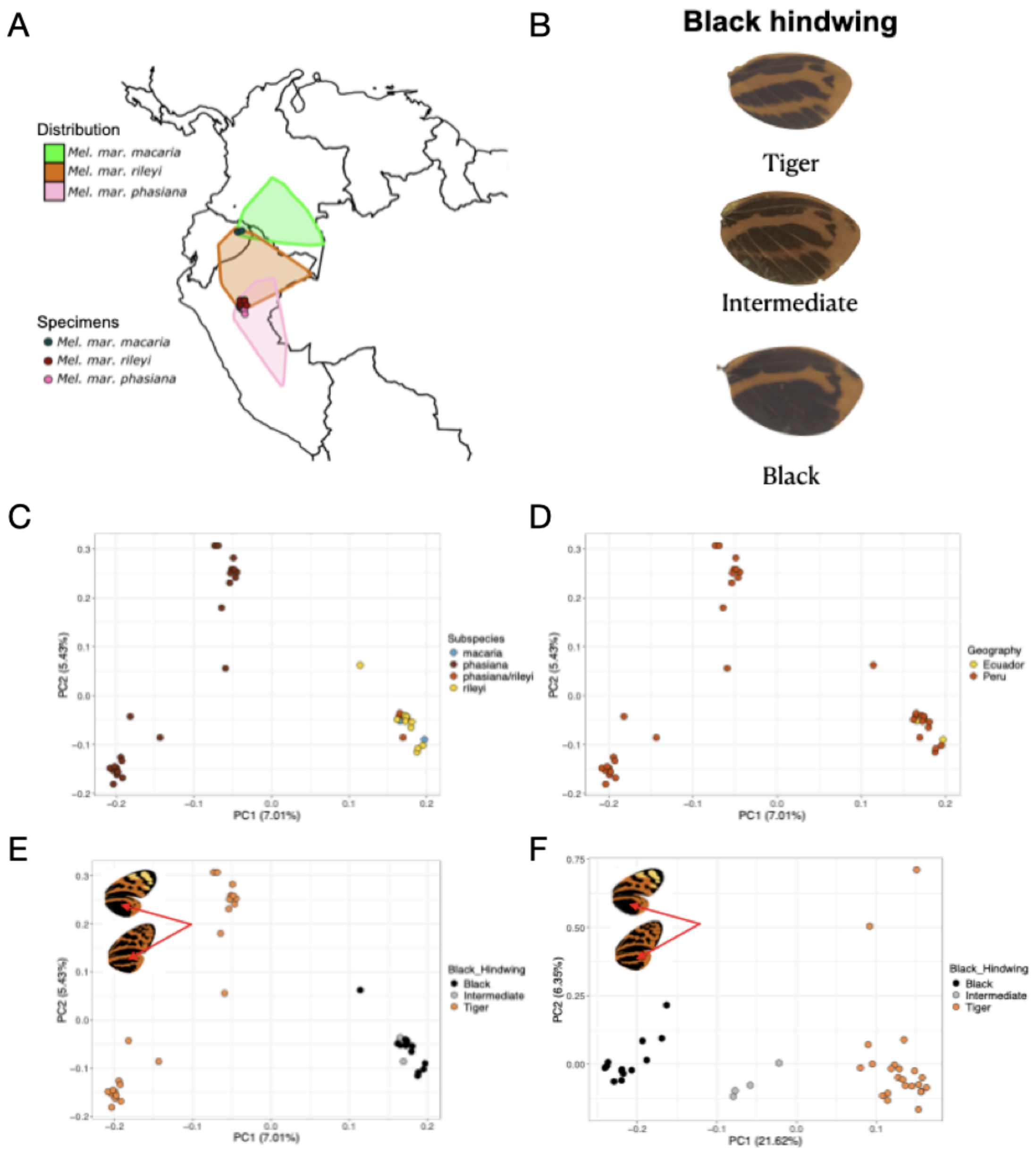


**Extended Data Fig. 19 | *Melinaea marseus*: taxon distribution, wing phenotypes and genetic PCA*.*** (A) Geographical distribution of *Melinaea marsaeus* subspecies, with specimen collection locations indicated. (B) Phenotypes used in the GWA. (C to F): Principal component analysis (PCA) showing the genetic distance between the sampled *Melianea marsaeus* individuals using a dataset consisting of 205,438 LD pruned biallelic SNPs (C-E) or 1,686 no LD pruned SNPs around the *Optix* region (F). The scatter plots correspond to the first two principal components (PCs). Points are coloured by subspecies (C), geography (D), hindwing black phenotype (E,F). The wing images highlight the colour-coded phenotypes.

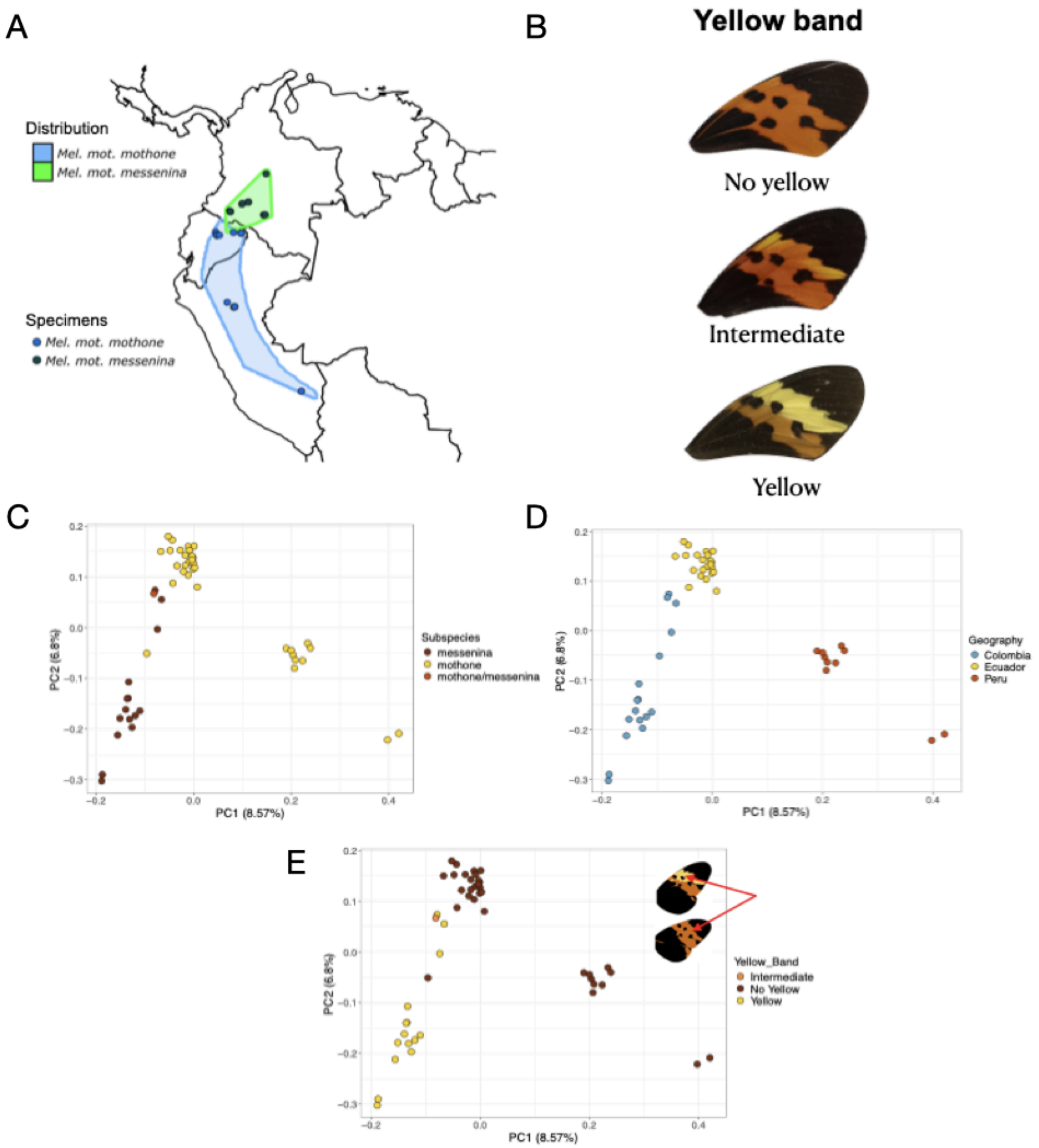


**Extended Data Fig. 20 | *Melinaea mothone*: taxon distribution, wing phenotypes and genetic PCA.** (A) Geographical distribution of *Melinaea mpothone* subspecies, with specimen collection locations indicated. (B) Phenotypes used in the GWA. (C to E): Principal component analysis (PCA) showing the genetic distance between the sampled *Melinaea mothone* individuals using a dataset consisting of 13,652 LD pruned biallelic SNPs. The scatter plots correspond to the first two principal components (PCs). Points are coloured by subspecies (C), geography (D) and yellow bar phenotype (E). The wing images highlight the colour-coded phenotypes.

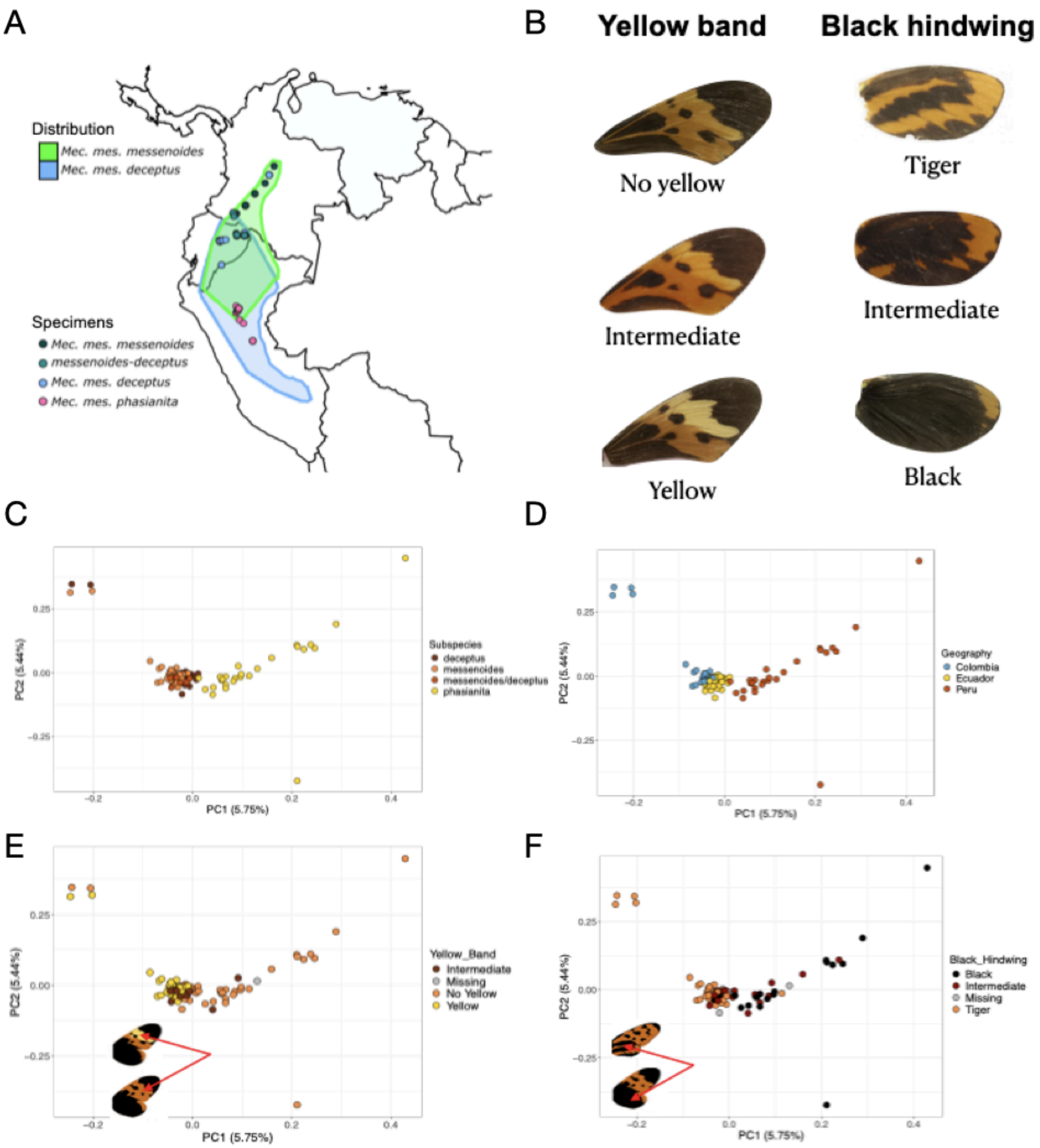


**Extended Data Fig. 21 | *Mechanitis messenoides* (forewing yellow bar and hindwing orange/black): taxon distribution, wing phenotypes and genetic PCA.** (A) Geographical distribution of *Mechanitis messenoides* subspecies, with specimen collection locations indicated. (B) Phenotypes used in the GWA. (C to F): Principal component analysis (PCA) showing the genetic distance between the sampled *Mechanitis messenoides* individuals using a dataset consisting of 86,312 LD pruned biallelic SNPS. The scatter plots correspond to the first two principal components (PCs). Points are coloured by subspecies (C), geography (D), yellow bar phenotype (E), and hindwing black phenotype (F). The wing images highlight the colour-coded phenotypes.

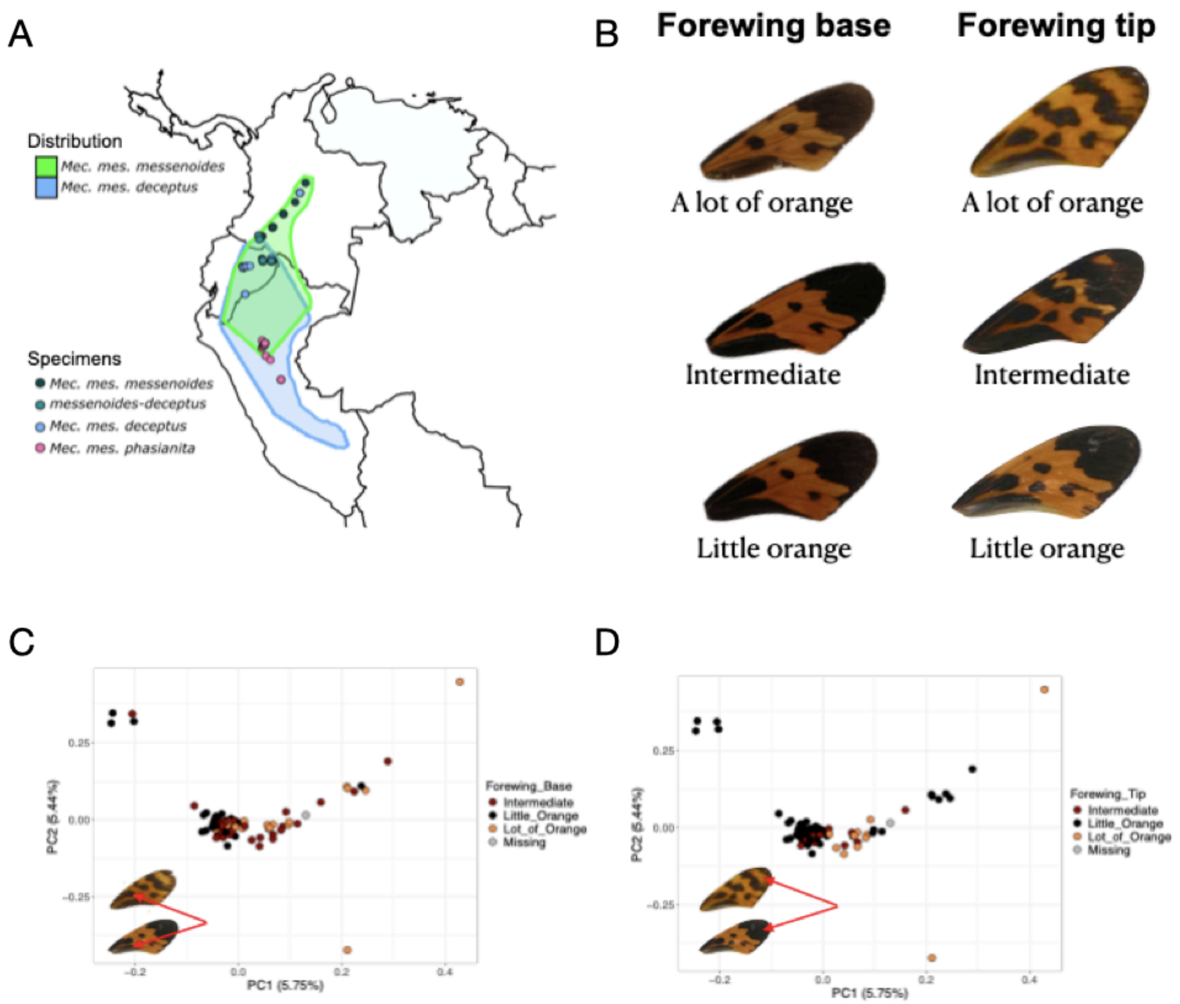


**Extended Data Fig. 22 | *Mechanitis messenoides* (forewing base and tip orange/black): taxon distribution, wing phenotypes and genetic PCA.** (A) Geographical distribution of *Mechanitis messenoides* subspecies, with specimen collection locations indicated. (B) Phenotypes used in the GWA. (C to D): Principal component analysis (PCA) showing the genetic distance between the sampled *Mechanitis messenoides* individuals using a dataset consisting of 86,312 LD pruned biallelic SNPS. The scatter plots correspond to the first two principal components (PCs). Points are coloured by forewing base phenotype (C) and forewing tip phenotype (D). The wing images highlight the colour-coded phenotypes.

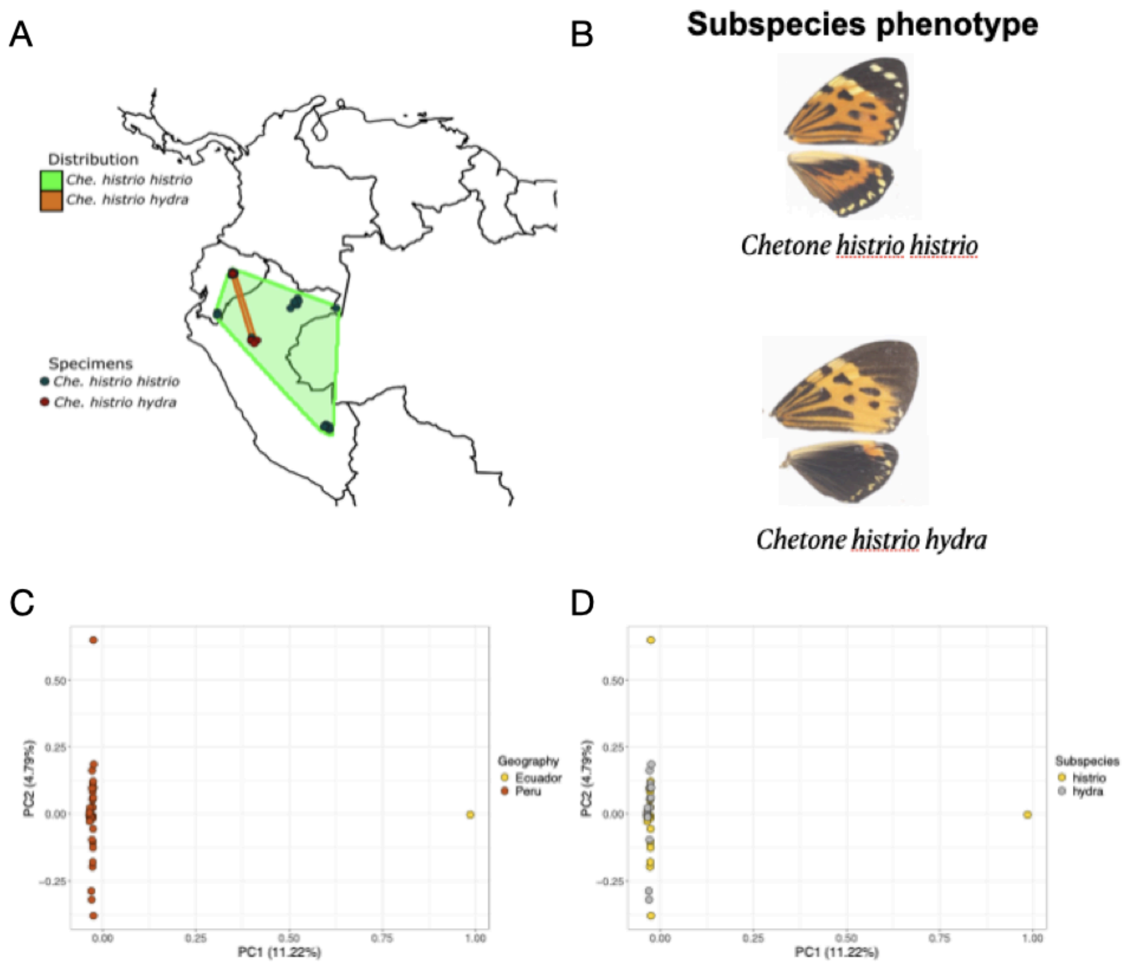


**Extended Data Fig. 23 | *Chetone histrio*: taxon distribution, wing phenotypes and genetic PCA.** **a,** Geographical distribution of *Chetone histrio* subspecies, with specimen collection locations indicated. **b,** Phenotypes used in the GWA. Principal Component Analysis (PCA) of 1,159,948 LD-pruned biallelic SNPs in *Chetone histrio* with points coloured by **c,** geography and **d,** subspecies.

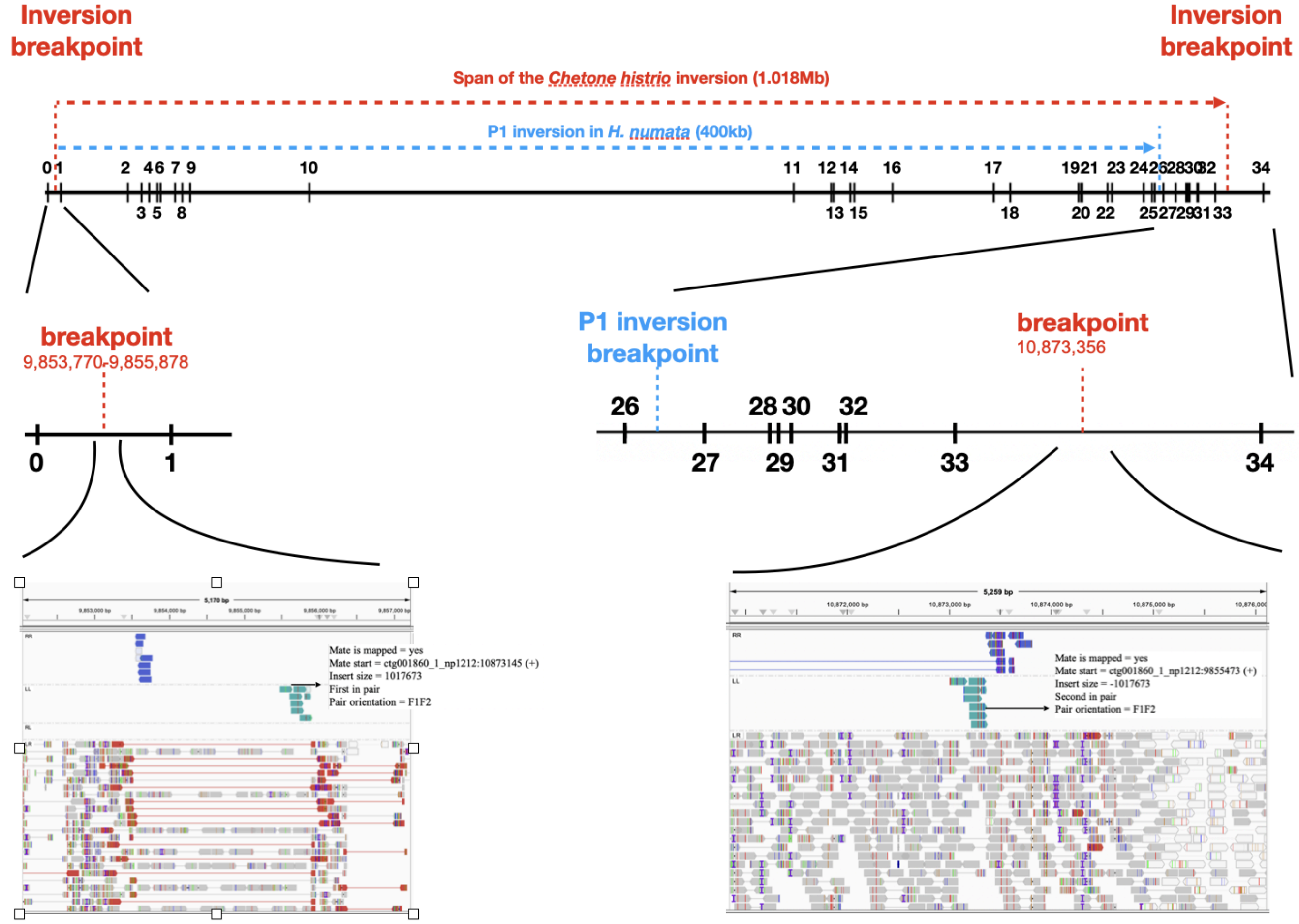


**Extended Data Fig. 24 | Inversion breakpoint analysis in *Chetone histrio*.** Examination of Illumina read-pair orientation and mapped insert sizes using IGV allows localisation of the inversion breakpoints in *Chetone histrio.* The IGV screenshots are for individual NR15-519, which is heterozygous for the inversion. a) Comparison of the locations of the *Chetone histrio* and *Heliconius numata* P1 inversion based on the locations of genes labelled 0-34. b) The left breakpoint in *Chetone histrio* is located between genes 0 and 1 at ctg001860_1_np2121:9,853,770-9,855,878. A region of poor mapping prevents more accurate inference of the breakpoint. c) The right breakpoint in *Chetone histrio* is located between genes 33 and 34 at ctg001860_1_np2121:10,873,356.

Gene 0-34 refer to: 0: *beta-fructofuranosidase*; 1: *glutaminyl-peptidecyclotransferase*; 2: *HMEL000021*; 3: *enoyl-CoA hydratase*; 4: *Cancer-related nucleoside-triphosphatase*; 5: *Sur-8/LRR*; 6: *HMEL032678*; 7: *HMEL002023g1*; 8: *HMEL002023g2*; 9: *HMEL002024*; 10: *cortex*; 11: *parn*; 12: *HMEL000027*; 13: *ARP-like*; 14: *ATP synthase subunit f*; 15: *proteasome 26S non ATPasesubunit 4*; 16: *zinc phosphodiesterase*; 17: *serine/threonine-proteinkinase*; 18: *WD repeat-containing protein 19*; 19: *HMEL013472*; 20: *WAS protein family homologue 1*; 21: *Domeless*; 22: *HMEL032681*; 23: *HMEL032683*; 24: *mitogen-activated protein kinase*; 25: *DNA excision repair protein ERCC-6*; 26: *penguin*; 27: *thymidylate kinase*; 28: *caspase-activated DNase*; 29: *ribosome biogenesis regulatory protein*; 30: *INO80 complex subunit C*; 31: *uncharacterized WD repeat-containing protein C2E1P5.05*; 32: *Sr protein*; 33: *HMEL000048*; 34: *HMEL000049*.

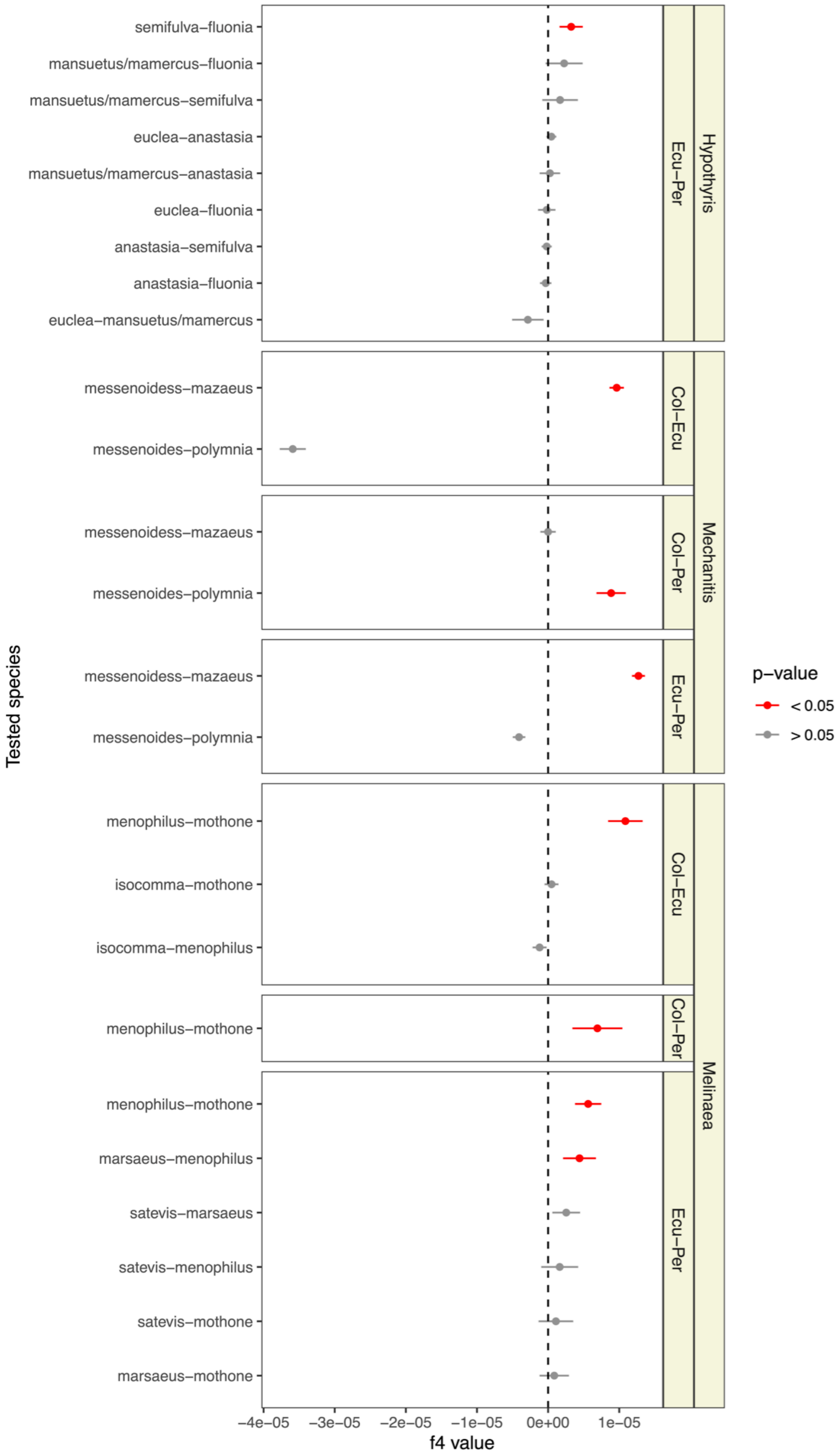


**Extended Data Fig. 25 | $f_4$ statistics testing for allele sharing between sympatric species of *Hypothyris*, *Mechanitis*, and *Melinaea*.** Comparisons are shown across different geographic regions (Colombia-Ecuador, Colombia-Peru and Ecuador-Peru). Each point represents an $f_4$ value, with standard errors. Red points are significantly different from zero ($p < 0.05$). Significantly positive $f_4$ value suggests excess allele sharing between the tested species, consistent with interspecific gene flow. Details of the taxa tested are shown in Table S5.

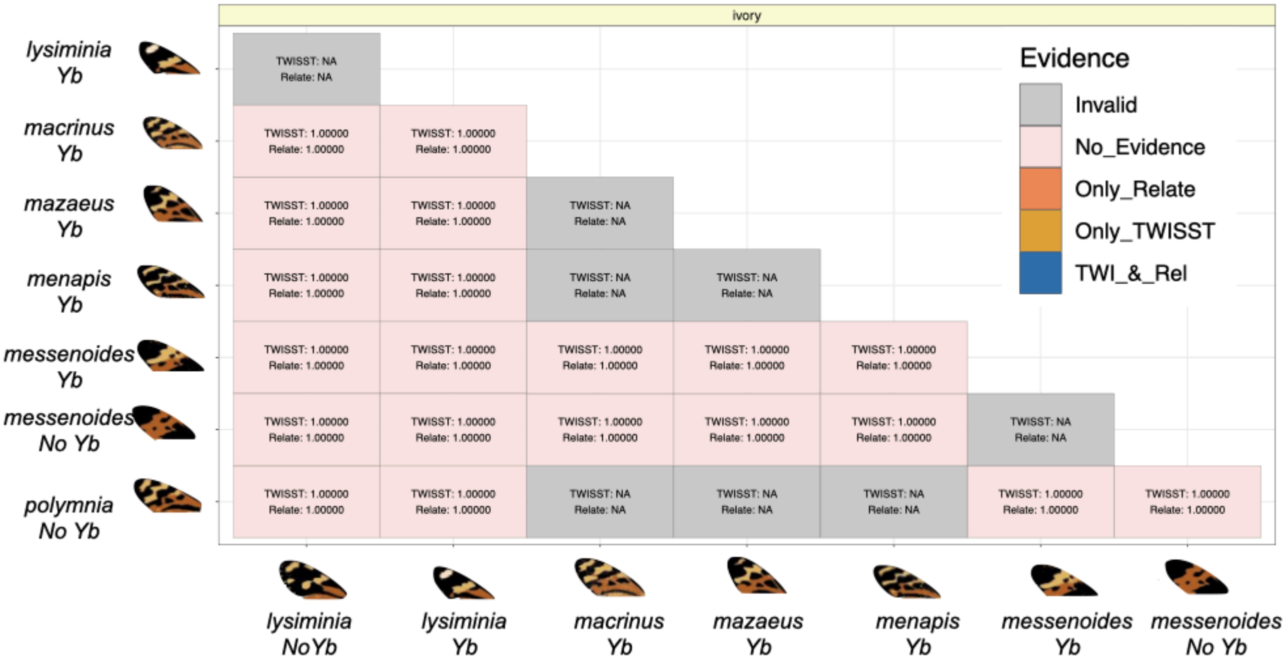
ivory
Evidence
Invalid
No_Evidence
Only_Relate
Only_TWISST
TWI_&_Rel
lysiminia Yb
macrinus Yb
mazaeus Yb
menapis Yb
messenoides Yb
messenoides No Yb
polymnia No Yb
lysiminia NoYb
lysiminia Yb
macrinus Yb
mazaeus Yb
menapis Yb
messenoides Yb
messenoides No Yb

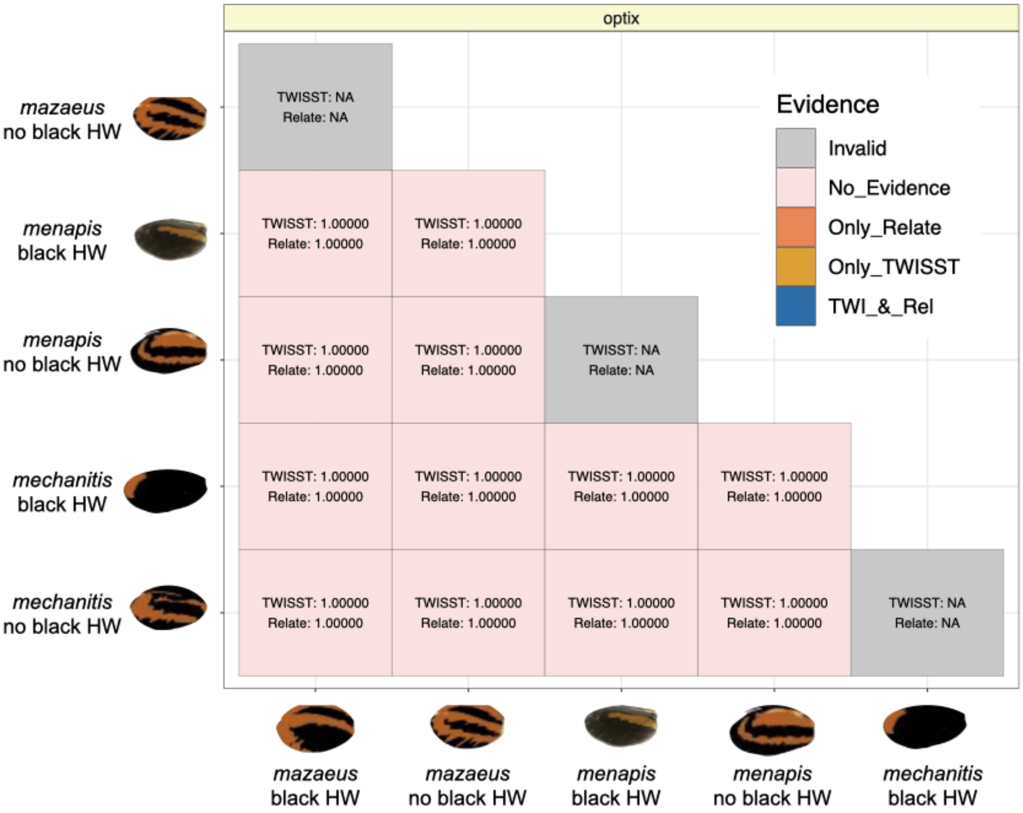
optix
Evidence
Invalid
No_Evidence
Only_Relate
Only_TWISST
TWI_&_Rel
mazaeus no black HW
menapis black HW
menapis no black HW
mechanitis black HW
mechanitis no black HW
mazaeus black HW
mazaeus no black HW
menapis black HW
menapis no black HW
mechanitis black HW

**Extended Data Fig. 26 | Testing for introgression across pairs of *Mechanitis* species at *ivory* and *optix.*** Comparisons are shown between species with different forewing yellow bar (top) and hindwing orange/black (bottom) phenotypes.. Each cell in the matrix represents a comparison between a pair of species, with rows and columns labelled by species and wing phenotype. The matrix is colour-coded to indicate the type of evidence detected: no evidence of introgression (pink), evidence from Relate only (orange), evidence from Twisst only (ochre), or evidence from both Relate and Twisst (blue). The Twisst and Relate p-values, based on a block permutation test, are displayed in each cell. "NA" indicates intraspecific or invalid comparisons. P-values of 1.00000 indicate that no introgression-compatible topologies were observed within the GWAS peak region, making it impossible to compute a p-value. For Relate, “low” indicates that the analyses included fewer than 20 samples and could not be run. Species and wing phenotypes are depicted along the matrix's margins with corresponding butterfly illustrations. Details of the taxa tested are shown in Table S4.

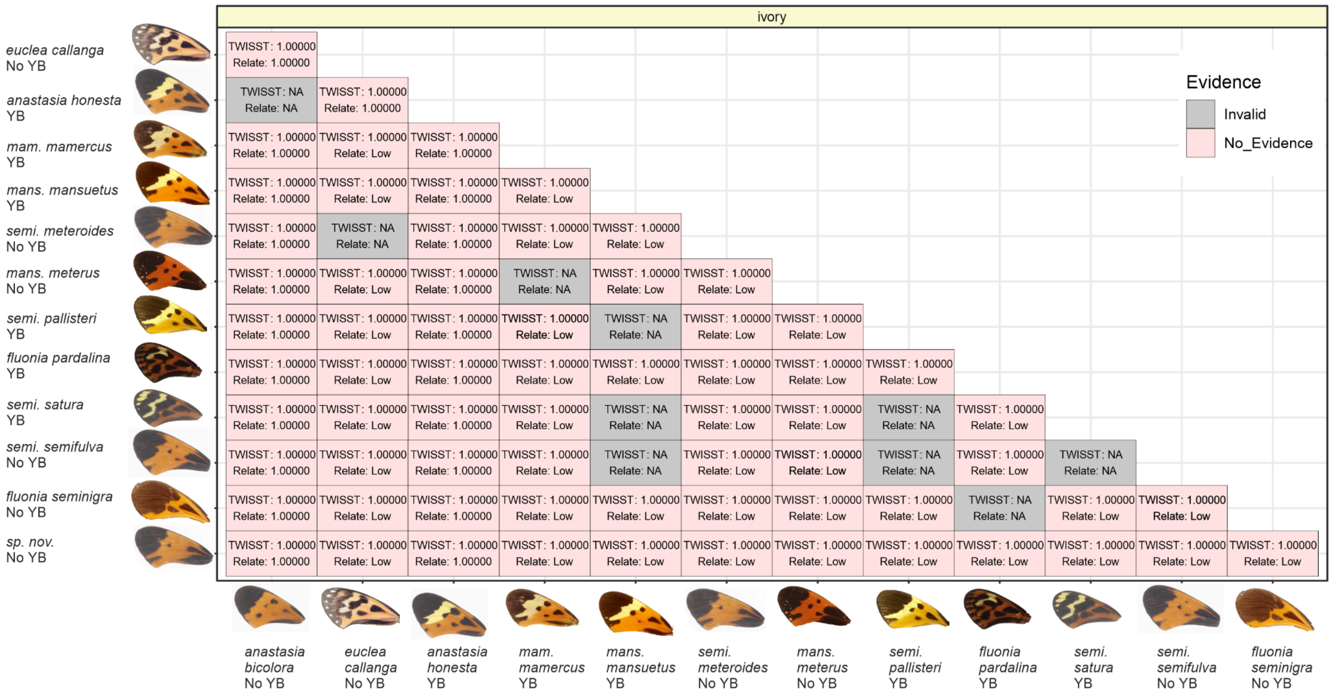


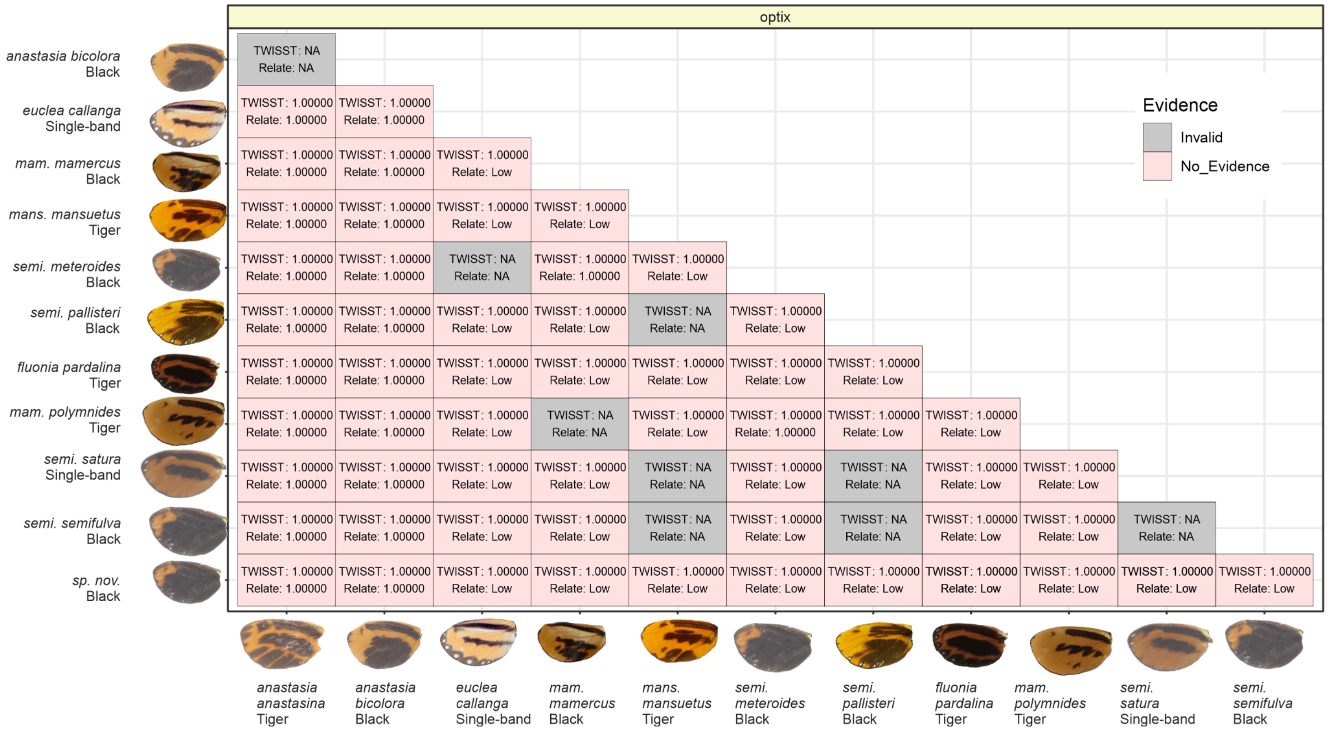


**Extended Data Fig. 27 | Testing for introgression across pairs of *Hypothyris* species at *ivory* and *optix*.** Comparisons are shown between species with different forewing yellow bar (top) and hindwing orange/black (bottom) phenotypes. Each cell in the matrix represents a comparison between a pair of species, with rows and columns labelled by species and wing phenotype. No evidence of introgression is found in any comparison. The Twisst and Relate p-values, based on a block permutation test, are displayed in each cell. "NA" indicates intraspecific or invalid comparisons. P-values of 1.00000 indicate that no introgression-compatible topologies were observed within the GWAS peak region, making it impossible to compute a p-value. For Relate, “low” indicates that the analyses included fewer than 20 samples and could not be run. Species and wing phenotypes are depicted along the matrix's margins with corresponding wing photographs. Details of the taxa tested are shown in Table S4.

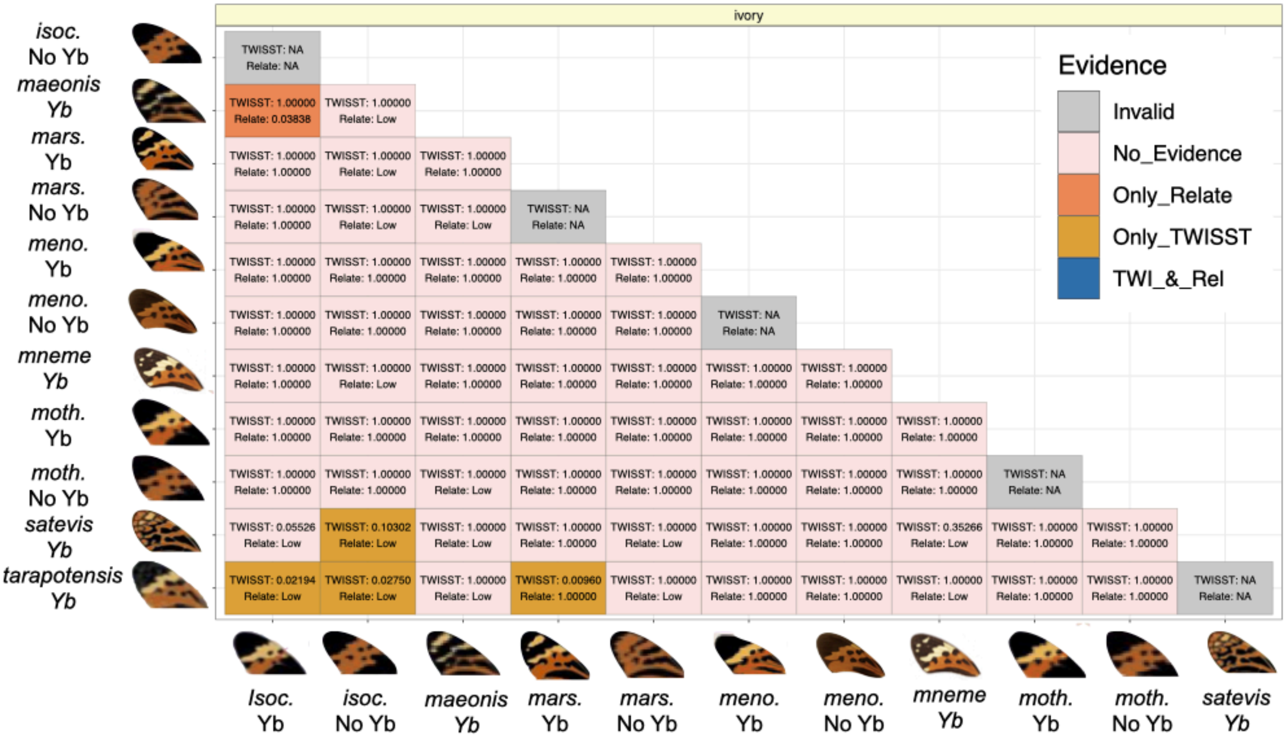
ivory
Evidence
Invalid
No_Evidence
Only_Relate
Only_TWISST
TWI_&_Rel
isoc. No Yb
maeonis Yb
mars. Yb
mars. No Yb
meno. Yb
meno. No Yb
mneme Yb
moth. Yb
moth. No Yb
satevis Yb
tarapotensis Yb
TWISST: NA Relate: NA
TWISST: 1.00000 Relate: 0.03838
TWISST: 1.00000 Relate: Low
TWISST: 1.00000 Relate: 1.00000
TWISST: 1.00000 Relate: Low
TWISST: 1.00000 Relate: 1.00000
TWISST: 1.00000 Relate: 1.00000
TWISST: 1.00000 Relate: Low
TWISST: 1.00000 Relate: Low
TWISST: NA Relate: NA
TWISST: 1.00000 Relate: 1.00000
TWISST: 1.00000 Relate: 1.00000
TWISST: 1.00000 Relate: 1.00000
TWISST: 1.00000 Relate: 1.00000
TWISST: 1.00000 Relate: 1.00000
TWISST: 1.00000 Relate: 1.00000
TWISST: 1.00000 Relate: 1.00000
TWISST: 1.00000 Relate: 1.00000
TWISST: 1.00000 Relate: 1.00000
TWISST: 1.00000 Relate: 1.00000
TWISST: NA Relate: NA
TWISST: 1.00000 Relate: 1.00000
TWISST: 1.00000 Relate: Low
TWISST: 1.00000 Relate: 1.00000
TWISST: 1.00000 Relate: 1.00000
TWISST: 1.00000 Relate: 1.00000
TWISST: 1.00000 Relate: 1.00000
TWISST: 1.00000 Relate: 1.00000
TWISST: 1.00000 Relate: 1.00000
TWISST: 1.00000 Relate: 1.00000
TWISST: 1.00000 Relate: 1.00000
TWISST: 1.00000 Relate: 1.00000
TWISST: 1.00000 Relate: 1.00000
TWISST: 1.00000 Relate: 1.00000
TWISST: 1.00000 Relate: 1.00000
TWISST: 1.00000 Relate: 1.00000
TWISST: 1.00000 Relate: 1.00000
TWISST: 1.00000 Relate: 1.00000
TWISST: 1.00000 Relate: Low
TWISST: 1.00000 Relate: 1.00000
TWISST: 1.00000 Relate: 1.00000
TWISST: 1.00000 Relate: 1.00000
TWISST: 1.00000 Relate: 1.00000
TWISST: 1.00000 Relate: 1.00000
TWISST: 1.00000 Relate: 1.00000
TWISST: NA Relate: NA
TWISST: 0.05526 Relate: Low
TWISST: 0.10302 Relate: Low
TWISST: 1.00000 Relate: Low
TWISST: 1.00000 Relate: 1.00000
TWISST: 1.00000 Relate: Low
TWISST: 1.00000 Relate: 1.00000
TWISST: 1.00000 Relate: 1.00000
TWISST: 0.35266 Relate: Low
TWISST: 1.00000 Relate: 1.00000
TWISST: 1.00000 Relate: 1.00000
TWISST: 0.02194 Relate: Low
TWISST: 0.02750 Relate: Low
TWISST: 1.00000 Relate: Low
TWISST: 0.00960 Relate: 1.00000
TWISST: 1.00000 Relate: Low
TWISST: 1.00000 Relate: 1.00000
TWISST: 1.00000 Relate: 1.00000
TWISST: 1.00000 Relate: Low
TWISST: 1.00000 Relate: 1.00000
TWISST: 1.00000 Relate: 1.00000
TWISST: NA Relate: NA
Isoc. Yb
isoc. No Yb
maeonis Yb
mars. Yb
mars. No Yb
meno. Yb
meno. No Yb
mneme Yb
moth. Yb
moth. No Yb
satevis Yb

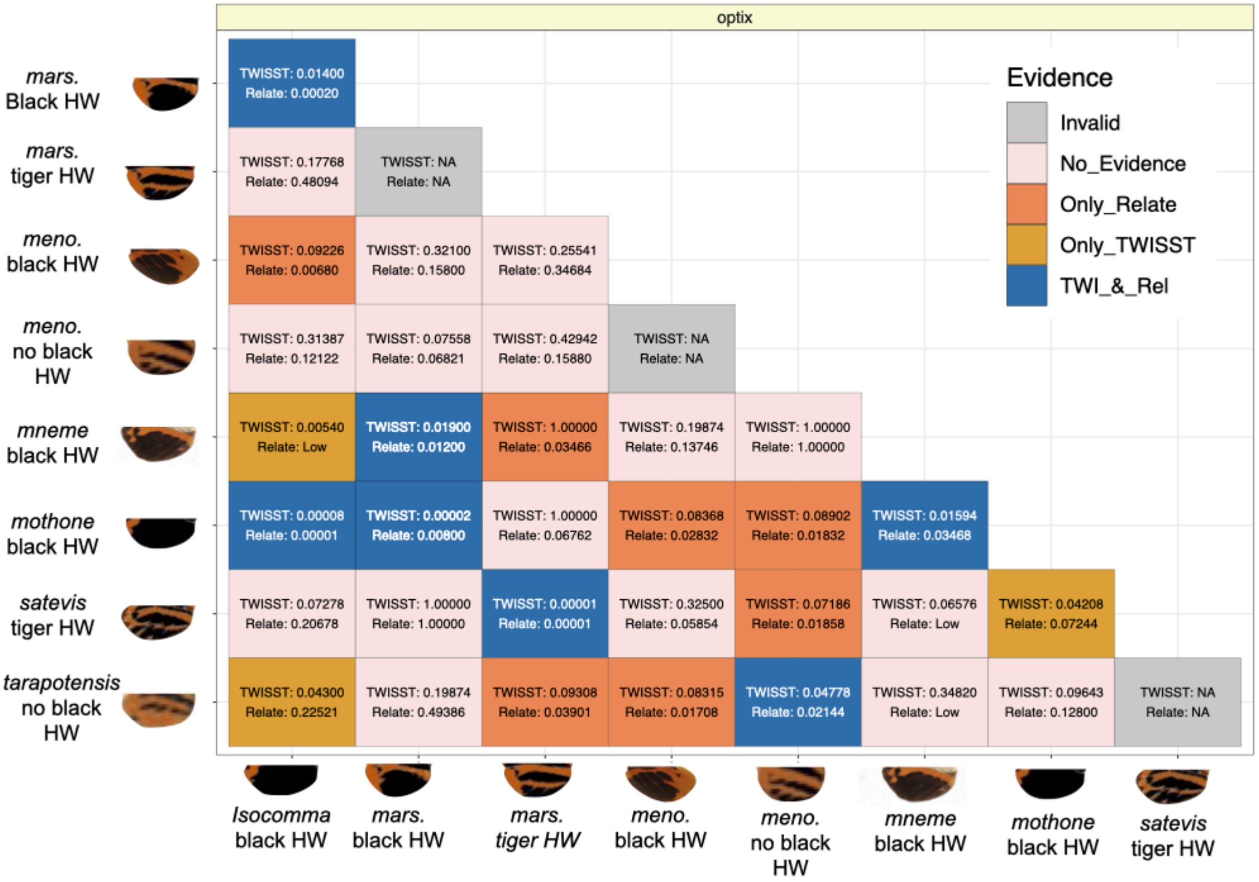
optix
Evidence
Invalid
No_Evidence
Only_Relate
Only_TWISST
TWI_&_Rel
mars. Black HW
mars. tiger HW
meno. black HW
meno. no black HW
mneme black HW
mothone black HW
satevis tiger HW
tarapotensis no black HW
TWISST: 0.01400 Relate: 0.00020
TWISST: 0.17768 Relate: 0.48094
TWISST: NA Relate: NA
TWISST: 0.09226 Relate: 0.00680
TWISST: 0.32100 Relate: 0.15800
TWISST: 0.25541 Relate: 0.34684
TWISST: 0.31387 Relate: 0.12122
TWISST: 0.07558 Relate: 0.06821
TWISST: 0.42942 Relate: 0.15880
TWISST: NA Relate: NA
TWISST: 0.00540 Relate: Low
TWISST: 0.01900 Relate: 0.01200
TWISST: 1.00000 Relate: 0.03466
TWISST: 0.19874 Relate: 0.13746
TWISST: 1.00000 Relate: 1.00000
TWISST: 0.00008 Relate: 0.00001
TWISST: 0.00002 Relate: 0.00800
TWISST: 1.00000 Relate: 0.06762
TWISST: 0.08368 Relate: 0.02832
TWISST: 0.08902 Relate: 0.01832
TWISST: 0.01594 Relate: 0.03468
TWISST: 0.07278 Relate: 0.20678
TWISST: 1.00000 Relate: 1.00000
TWISST: 0.00001 Relate: 0.00001
TWISST: 0.32500 Relate: 0.05854
TWISST: 0.07186 Relate: 0.01858
TWISST: 0.06576 Relate: Low
TWISST: 0.04208 Relate: 0.07244
TWISST: 0.04300 Relate: 0.22521
TWISST: 0.19874 Relate: 0.49386
TWISST: 0.09308 Relate: 0.03901
TWISST: 0.08315 Relate: 0.01708
TWISST: 0.04778 Relate: 0.02144
TWISST: 0.34820 Relate: Low
TWISST: 0.09643 Relate: 0.12800
TWISST: NA Relate: NA
Isocomma black HW
mars. black HW
mars. tiger HW
meno. black HW
meno. no black HW
mneme black HW
mothone black HW
satevis tiger HW

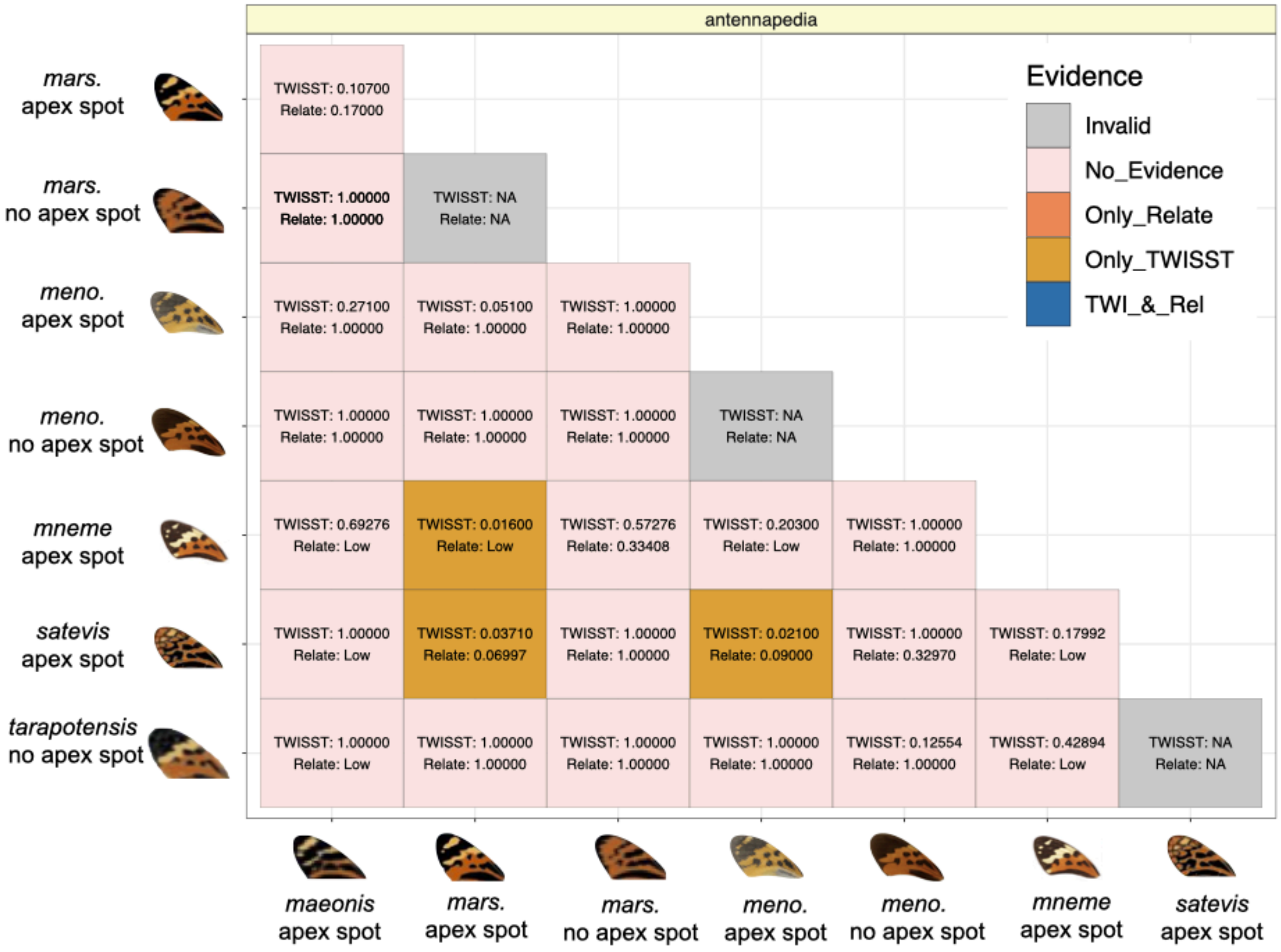


**Extended Data Fig. 28 | Testing for introgression across pairs of *Melinaea* species at *ivory, optix* and *antennapedia.*** Comparisons are shown between species with different forewing yellow bar (top), hindwing orange/black (middle) and forewing apex spot (bottom) phenotypes. Each cell in the matrix represents a comparison between a pair of species, with rows and columns labelled by species and wing phenotype. The matrix is colour-coded to indicate the type of evidence detected: no evidence of introgression (pink), evidence from Relate only (orange), evidence from Twisst only (ochre), or evidence from both Relate and Twisst (blue). The Twisst and Relate p-values, based on a block permutation test, are displayed in each cell. "NA" indicates intraspecific or invalid comparisons. P-values of 1.00000 indicate that no introgression-compatible topologies were observed within the GWAS peak region, making it impossible to compute a p-value. For Relate, “low” indicates that the analyses included fewer than 20 samples and could not be run. Species and wing phenotypes are depicted along the matrix's margins with corresponding butterfly illustrations. Abbreviations for species names are as follows: *meno*.: *menophius*; *mars*.: *marsaeus*; *isoc*.: *isocomma*; *moth*.: *mothone*. Details of the taxa tested are shown in Table S4.

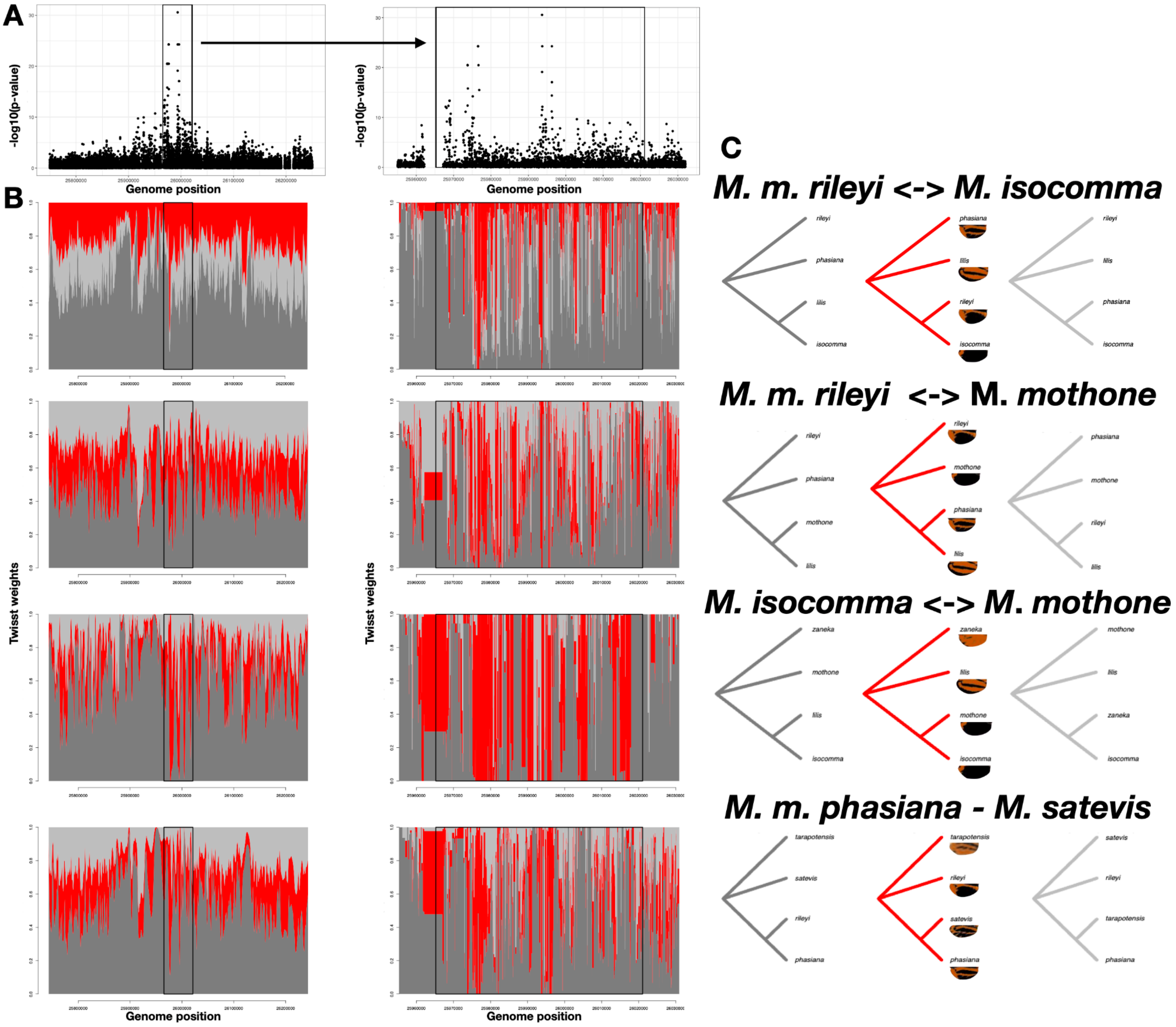


**Extended Data Fig. 29 | Genomic signals of introgression at the *optix* locus among four pairs of *Melinaea* species. a,** *Melinaea marsaeus* GWA Manhattan plot near *optix*. The black rectangle highlights the strongest association peak, which is shown zoomed in on the right panel. **b,** Twisst weight bar plots displaying local phylogenetic topology support across the highlighted region in **a**. Each row corresponds to a different species pair, with the y-axis indicating the Twisst weight, which quantifies how congruent the local genealogy is with one of the three topologies shown in **c,** The left bar plots provide a broader genomic context and include a smoothing function to reduce noise, while the right bar plots shows a zoom in of the region without smoothing. Higher red values indicate greater support for an introgressed topology, while the two shades of gray correspond to the two alternative topologies expected for unrooted trees based on four taxa. (**C**) Most likely marginal coalescent trees for each species pair tested, inferred using *RELATE*.

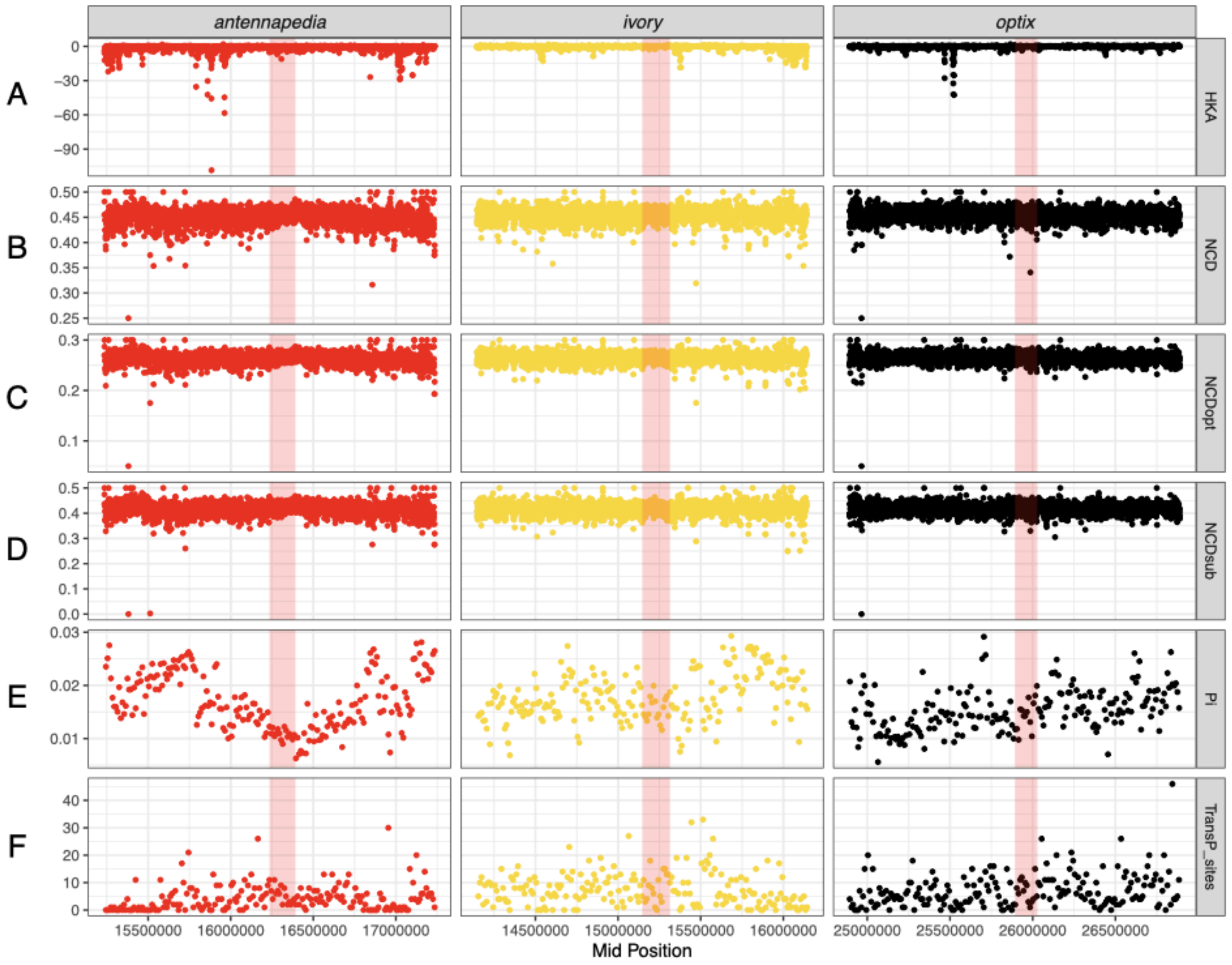


**Extended Data Fig. 30 | Multispecies balancing selection tests in *Melinaea* at *antennapedia*, *ivory* and *optix* (left to right).** These analyses included *Melinaea isocomma, Melinaea mothone, Melinaea marsaeus, Melinaea menophilus, Melinaea satevis and Melinaea tarapotensis*. The red-shaded regions indicate the location of the GWA peaks in the genes *antennapedia*, *ivory* and *optix*. A) HKAtrans statistics computed in 1 kb windows with a 500 bp sliding step. *Melinaea ludovica* was used to determine the ancestral state. Positive HKAtrans values indicate an excess of shared polymorphisms relative to divergence, which is consistent with ancient trans-species balancing selection. B) $NCD2_{trans}$ statistics computed in 1 kb windows with a 500 bp sliding step. This statistic measures allele frequency deviations from neutral expectations across multiple species. Values closer to 0 suggest ancient trans-species balancing selection. C) $NCD2_{trans\text{-}opt}$ statistics computed in 1 kb windows with a 500 bp sliding step. This statistic is a variant of $NCD2_{trans}$ that optimizes the target frequency at neutrality. Values closer to 0 suggest ancient trans-species balancing selection. D) $NCD2_{trans\text{-}sub}$ statistics computed in 1 kb windows with a 500 bp sliding step. A variant of $NCD2_{trans}$ that treats substitutions and polymorphisms separately. Values closer to 0 suggest ancient trans-species balancing selection. E) Multispecies nucleotide diversity (π) computed in 10 kb windows. Larger values are indicative of ancient trans-species balancing selection. F) Number of transpolymorphic sites computed in 10 kb windows. Larger values are indicative of ancient trans-species balancing selection.

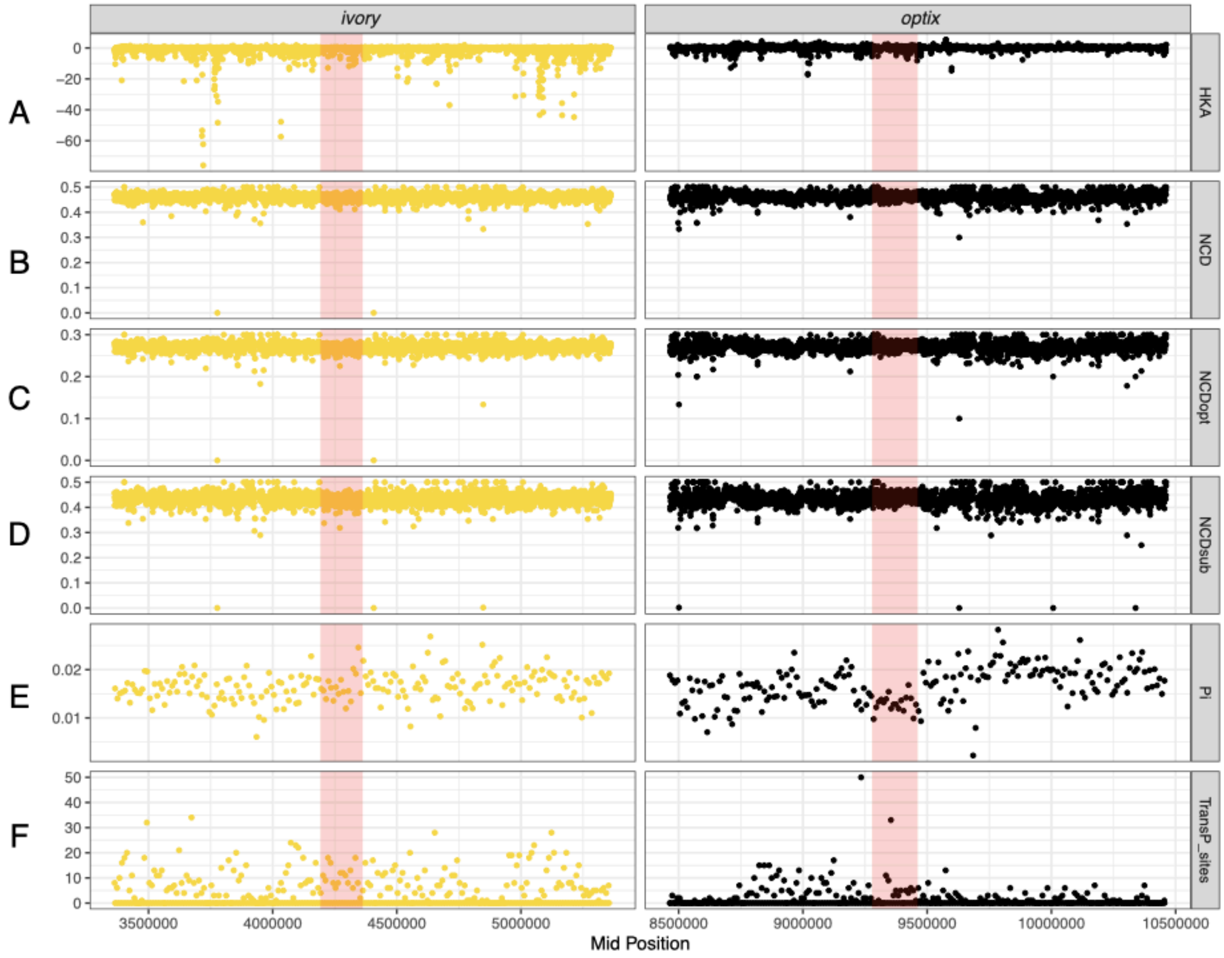


**Extended Data Fig. 31 | Multispecies balancing selection tests in *Hypothyris* at *ivory* and *optix* (left to right).** These analyses included *Hypothyris anastasia, Hypothyris euclea, Hypothyris ninonia and Hypothyris semifulva*. The red-shaded regions indicate the location of the GWA peaks in the genes *ivory* and *optix*. A) HKAtrans statistics computed in 1 kb windows with a 500 bp sliding step. *Hyalyris antea* was used to determine the ancestral state. Positive HKAtrans values indicate an excess of shared polymorphisms relative to divergence, which is consistent with ancient trans-species balancing selection. B) $NCD2_{trans}$ statistics computed in 1 kb windows with a 500 bp sliding step. This statistic measures allele frequency deviations from neutral expectations across multiple species. Values closer to 0 suggest ancient trans-species balancing selection. C) $NCD2_{trans\text{-}opt}$ statistics computed in 1 kb windows with a 500 bp sliding step. This statistic is a variant of $NCD2_{trans}$ that optimizes the target frequency at neutrality. Values closer to 0 suggest ancient trans-species balancing selection. D) $NCD2_{trans\text{-}sub}$ statistics computed in 1 kb windows with a 500 bp sliding step. A variant of $NCD2_{trans}$ that treats substitutions and polymorphisms separately. Values closer to 0 suggest ancient trans-species balancing selection. E) Multispecies nucleotide diversity (π) computed in 10 kb windows. Larger values are indicative of ancient trans-species balancing selection. F) Number of transpolymorphic sites computed in 10 kb windows. Larger values are indicative of ancient trans-species balancing selection.

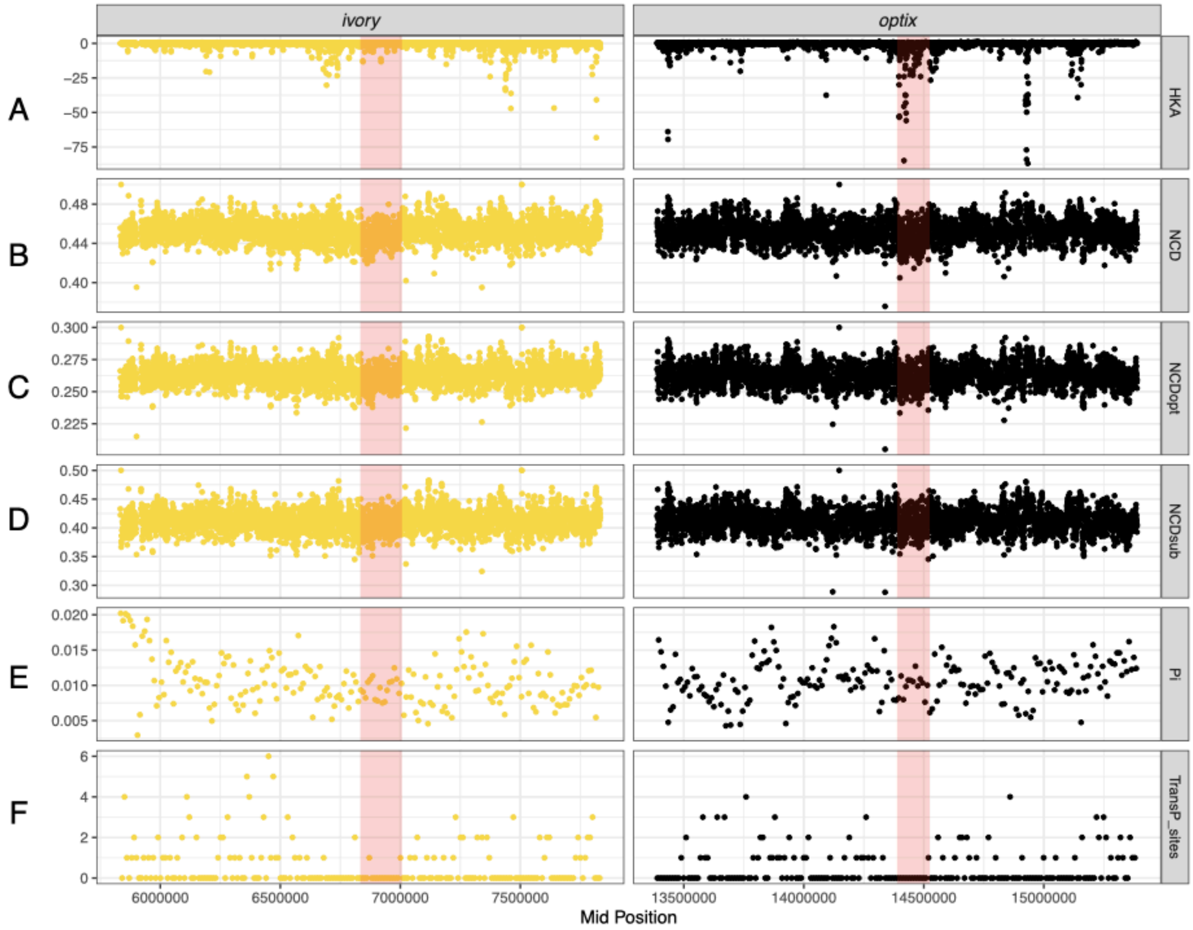


**Extended Data Fig. 32 | Multispecies balancing selection tests in *Mechanitis* at *ivory* and *optix* (left to right).** These analyses included *Mechanitis messenoides, Mechanitis lysimnia, Mechanitis macrinus, Mechanitis mazaeus, Mechanitis menapis, Mechanitis nesaea and Mechanitis polymnia*. The red-shaded regions indicate the location of the GWA peaks in the genes *ivory* and *optix*. A) HKAtrans statistics computed in 1 kb windows with a 500 bp sliding step. *Forbestra olivencia, Forbestra proceris* and *Forbestra equicola* were used to determine the ancestral state. Positive HKAtrans values indicate an excess of shared polymorphisms relative to divergence, which is consistent with ancient trans-species balancing selection. B) $NCD2_{trans}$ statistics computed in 1 kb windows with a 500 bp sliding step. This statistic measures allele frequency deviations from neutral expectations across multiple species. Values closer to 0 suggest ancient trans-species balancing selection. C) $NCD2_{trans\text{-}opt}$ statistics computed in 1 kb windows with a 500 bp sliding step. This statistic is a variant of $NCD2_{trans}$ that optimises the target frequency at neutrality. Values closer to 0 suggest ancient trans-species balancing selection. D) $NCD2_{trans\text{-}sub}$ statistics computed in 1 kb windows with a 500 bp sliding step. A variant of $NCD2_{trans}$ that treats substitutions and polymorphisms separately. Values closer to 0 suggest ancient trans-species balancing selection. E) Multispecies nucleotide diversity (π) computed in 10 kb windows. Larger values are indicative of ancient trans-species balancing selection. F) Number of transpolymorphic sites computed in 10 kb windows. Larger values are indicative of ancient trans-species balancing selection.

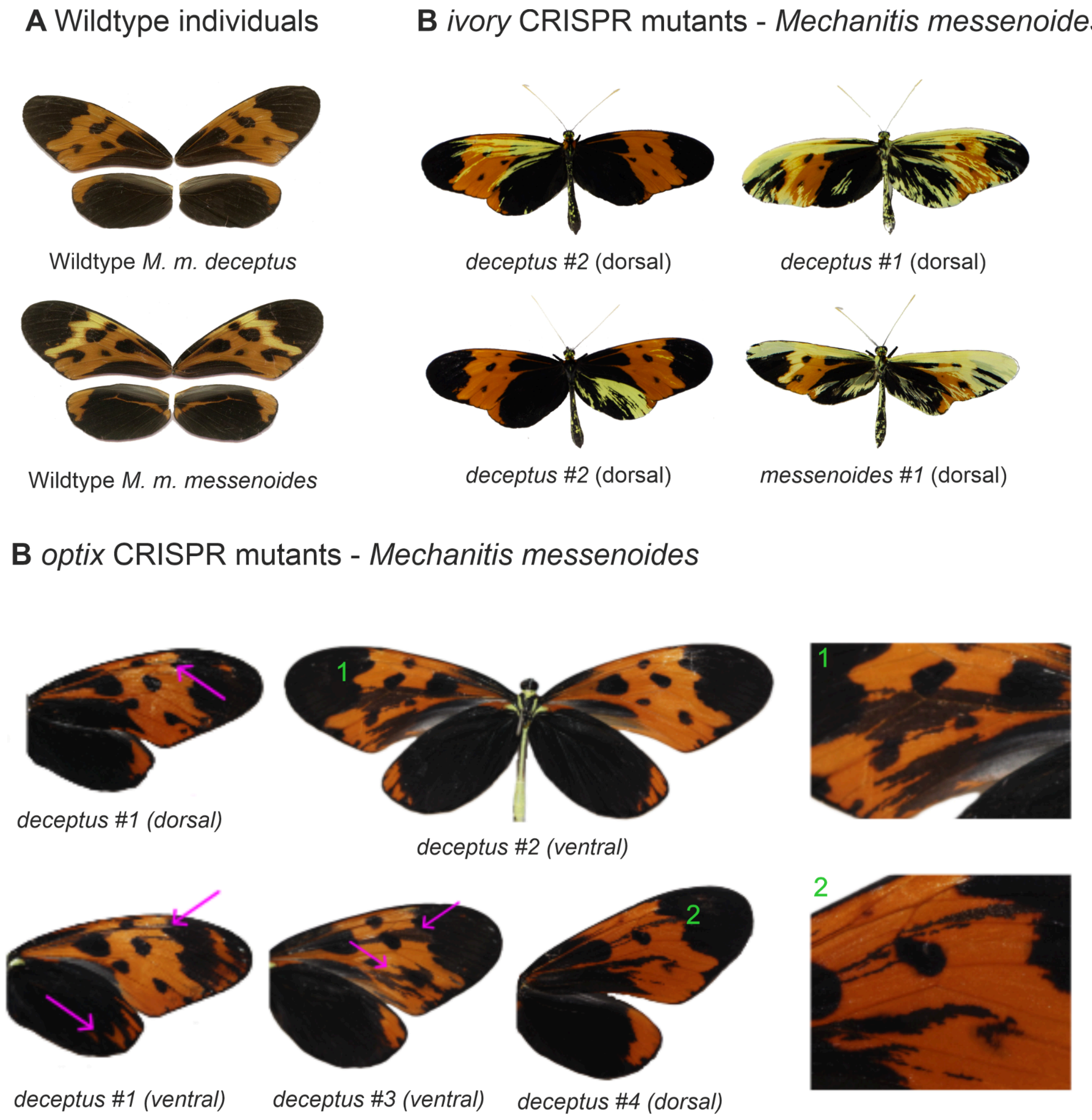


**Extended Data Fig. 33 | *Mechanitis messenoides* CRISPR *ivory* and *optix* mutants.** Mutant phenotypes can be recognized through their asymmetry (wildtype individuals have symmetric left and right wings. **A)** Wildtype individuals of the *messenoides* and *deceptus* subspecies. **B)** *ivory* mutants in which orange and black scales have turned yellow. **C)** *optix* mutants in which orange scales have turned black.

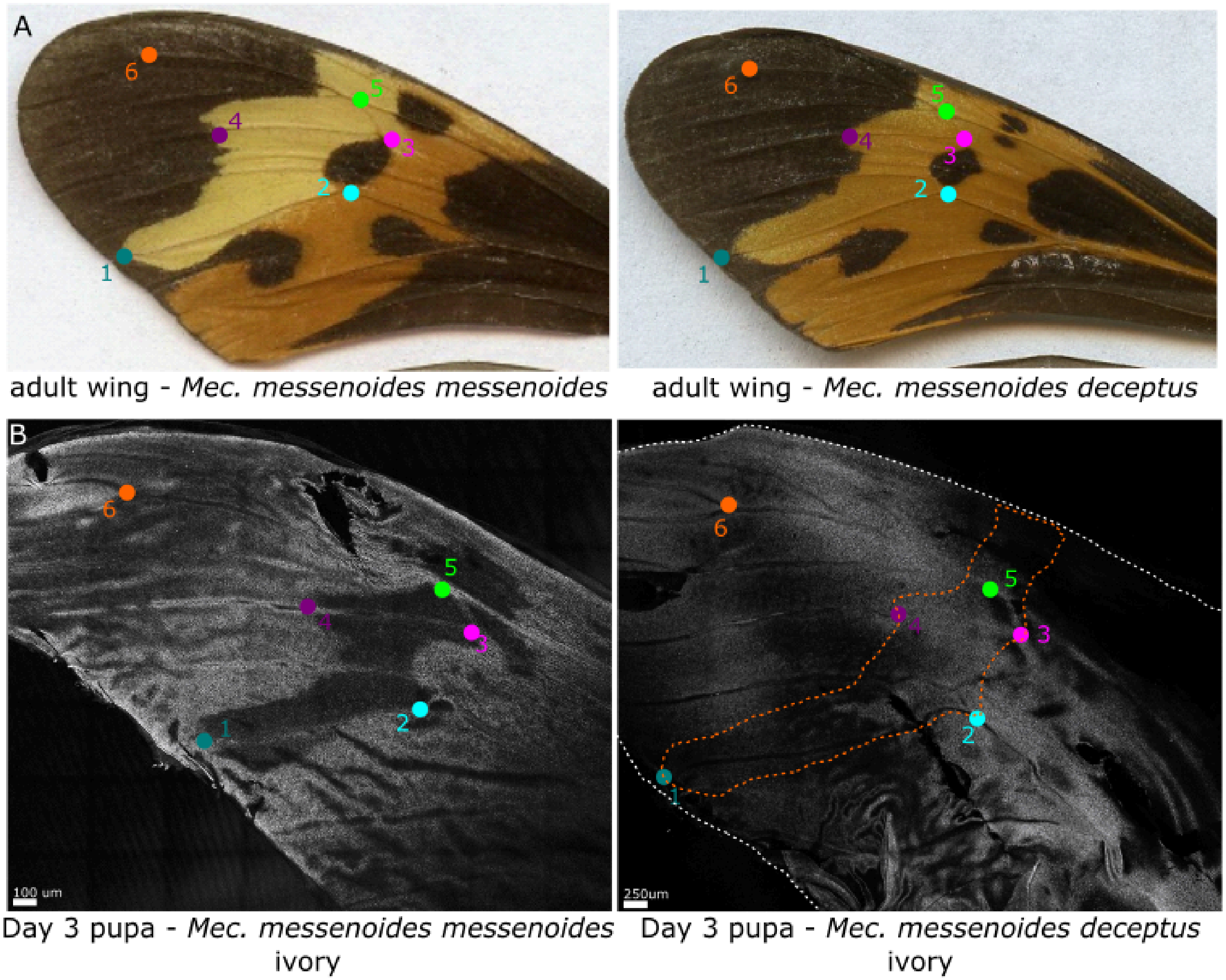


**Extended Data Fig. 34 | *Mechanitis messenoides in situ* hybridisation for *ivory***. Comparing non-yellow barred *Mec. messenoides deceptus* and yellow-barred *Mec. messenoides messenoides* shows absence of *ivory* RNA in the yellow-bar region of the forewing. The coloured dots indicate vein-based wing landmarks, demarcating the yellow bar region which lacks *ivory* expression in *Mec. messenoides messenoides*. In the non-yellow barred *Mec. messenoides deceptus*, the red dotted line demarcates the approximate region corresponding to the yellow bar region.

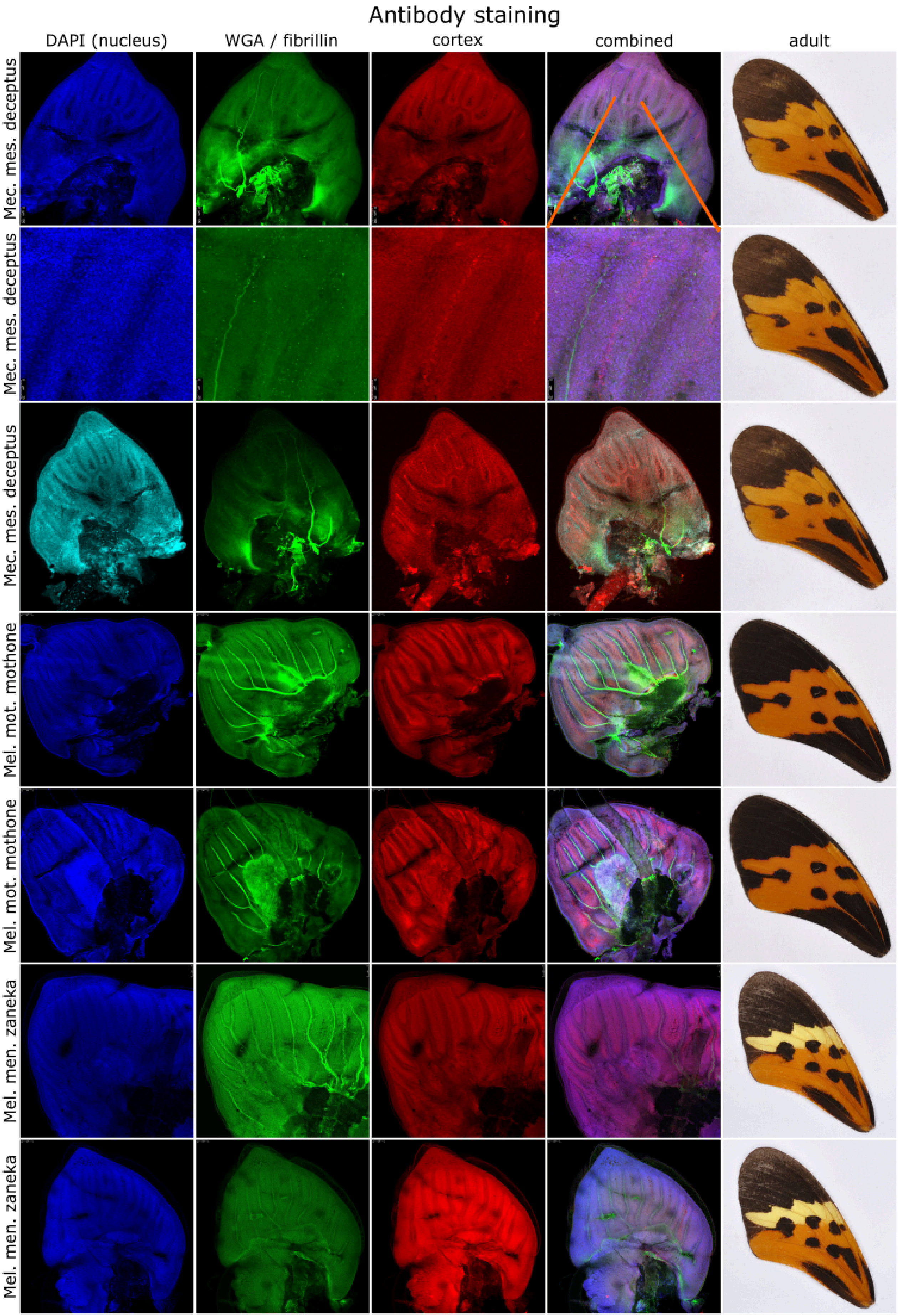
Antibody staining
DAPI (nucleus)
WGA / fibrillin
cortex
combined
adult
Mec. mes. deceptus
Mec. mes. deceptus
Mec. mes. deceptus
Mel. mot. mothone
Mel. mot. mothone
Mel. men. zaneka
Mel. men. zaneka

**Extended Data Fig. 35 | *Cortex* expression is not associated with the forewing yellow bar phenotype.** *Cortex* is expressed across the entire forewings of early fifth instar larvae. Each row shows a set of photos taken on a confocal microscope, with each column showing a different channel (column 1: nuclear DAPI staining (405nm), column 2: wheat germ agglutinin (WGA) that stains the nuclear membrane (488nm) or fibrillin, which stains the nucleolus: column 3: *Cortex* antibody (555 nm); column 4: an overlay image with all three channels combined). The last column shows the adult phenotype. Row 1-3: *Mechanitis messenoides deceptus*, with row 2 being a more zoomed image of row 1. Row 4-5: *Melinaea mothone mothone* (no yellow bar). Row 6-7: *Melinaea menophilus zaneka* (with yellow bar).

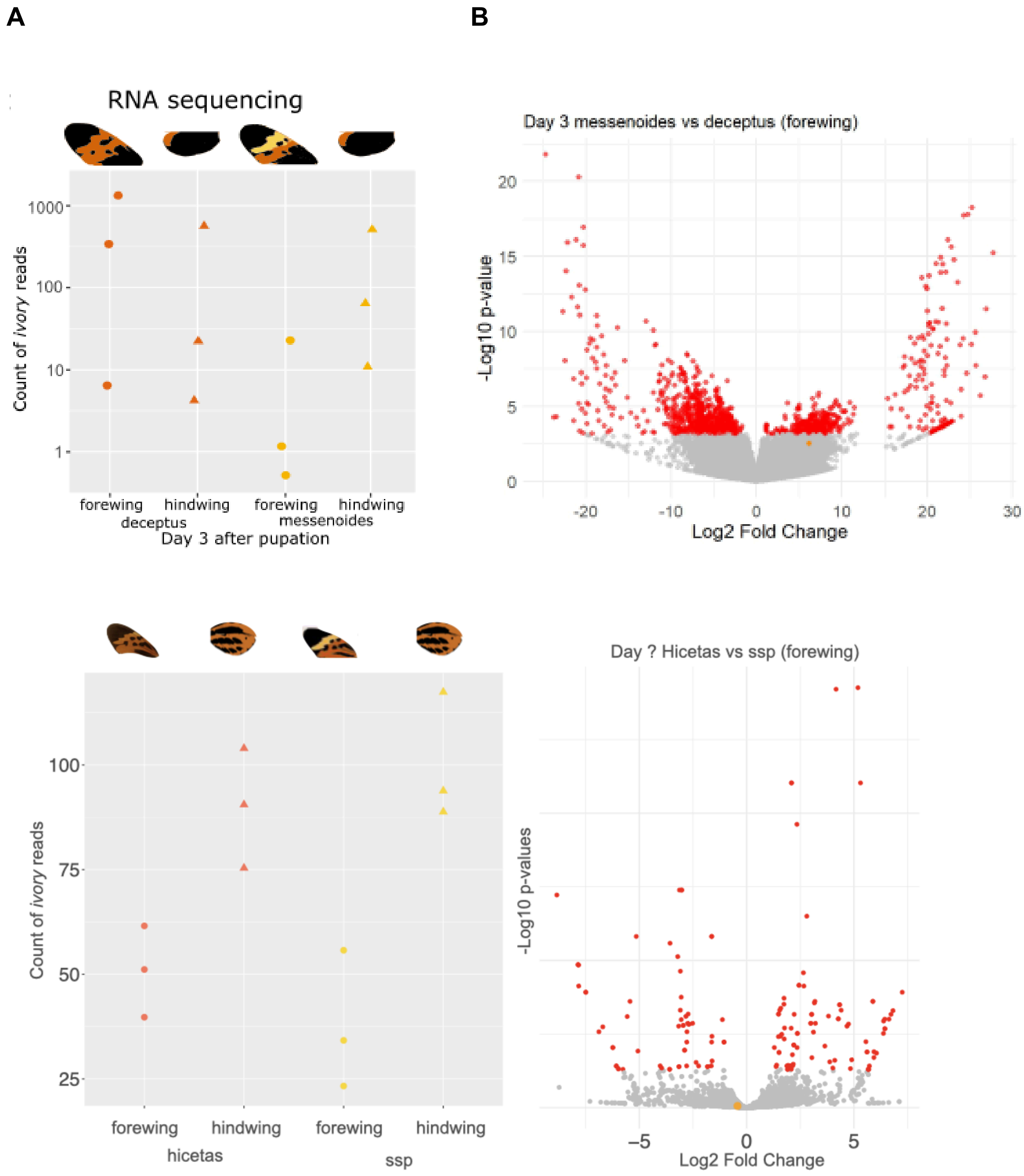


**Extended Data Fig. 36 | Differential gene expression analysis in pupal wing discs.** (A) Normalised *ivory* expression levels in the forewing and hindwing across subspecies of *Mechanitis messenoides* (top) and *Melinaea menophilus* (bottom). (B) Volcano plots of genome-wide differential expression analysis comparing yellow-barred versus non-yellow-barred forewings in *Mechanitis messenoides* (top) and *Melinaea menophilus* (bottom). Red dots indicate significantly differentially expressed genes; the orange dot corresponds to *ivory*.

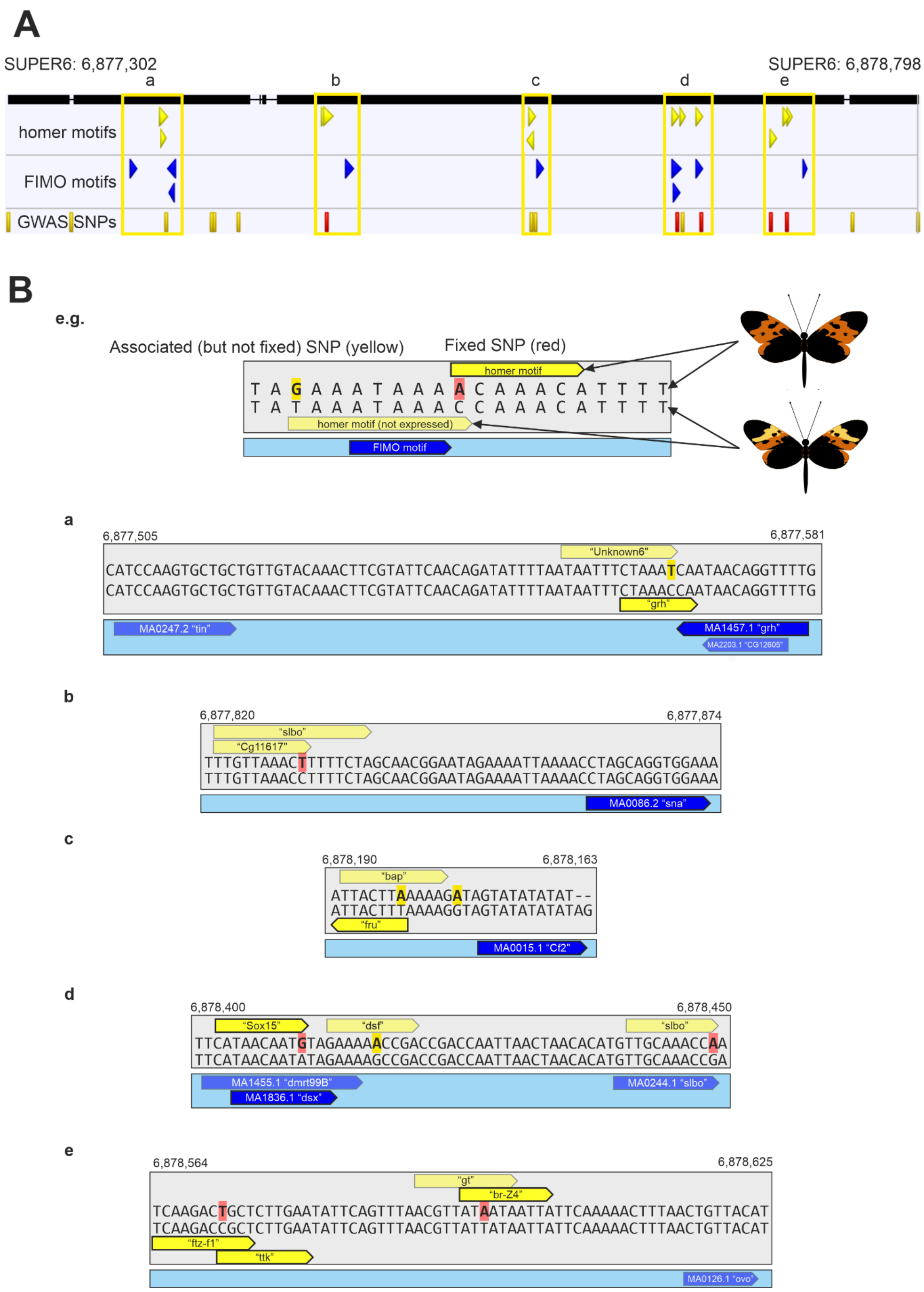


**Extended Data Fig. 37 | Transcription factor binding site analysis in *Mechantis messenoides*. A:** The GWAS peak region ranging to one (weakly- or non-associated) SNP either side of the fixed and most highly associated SNPs (SUPER_6:6877302-6878798) annotated with the motifs identified by homer (yellow arrows) which appear in 100% of sequences for one form and less than 5% of sequences for the other. Highly variable or repetitive motifs which occurred less than 3 times per sequence were excluded. Motifs identified by FIMO[108] which fall within 50 bp of an associated SNP detected by GWAS are shown in blue. FIMO motifs located on other parts of the scaffold are not shown. Fixed SNPs are highlighted in red, whilst associated SNPs which are not completely fixed are shown in yellow. **B:** Regions surrounding the homer motifs and closest FIMO motifs are shown in detail. The nucleotide sequences are the consensus sequence from the non-yellow barred form (*deceptus*, top) and yellow-barred reference form (*messenoides*, bottom). Homer[107] motifs which are enriched in the non-yellow-barred sequences are shown above this consensus sequence, whilst motifs enriched in the yellow-barred form are shown below. FIMO motifs were detected in both sets of sequences. Faded motif labels indicate transcription factors that are not expressed in forewing pupal wing discs. Associated/fixed SNPs are highlighted in the non-yellow-barred sequence according to the nucleotide present. The names of the FIMO motifs correspond to the JASPAR database, whilst the names of the homer motifs come from the "best-guess" matching motif identified by homer.

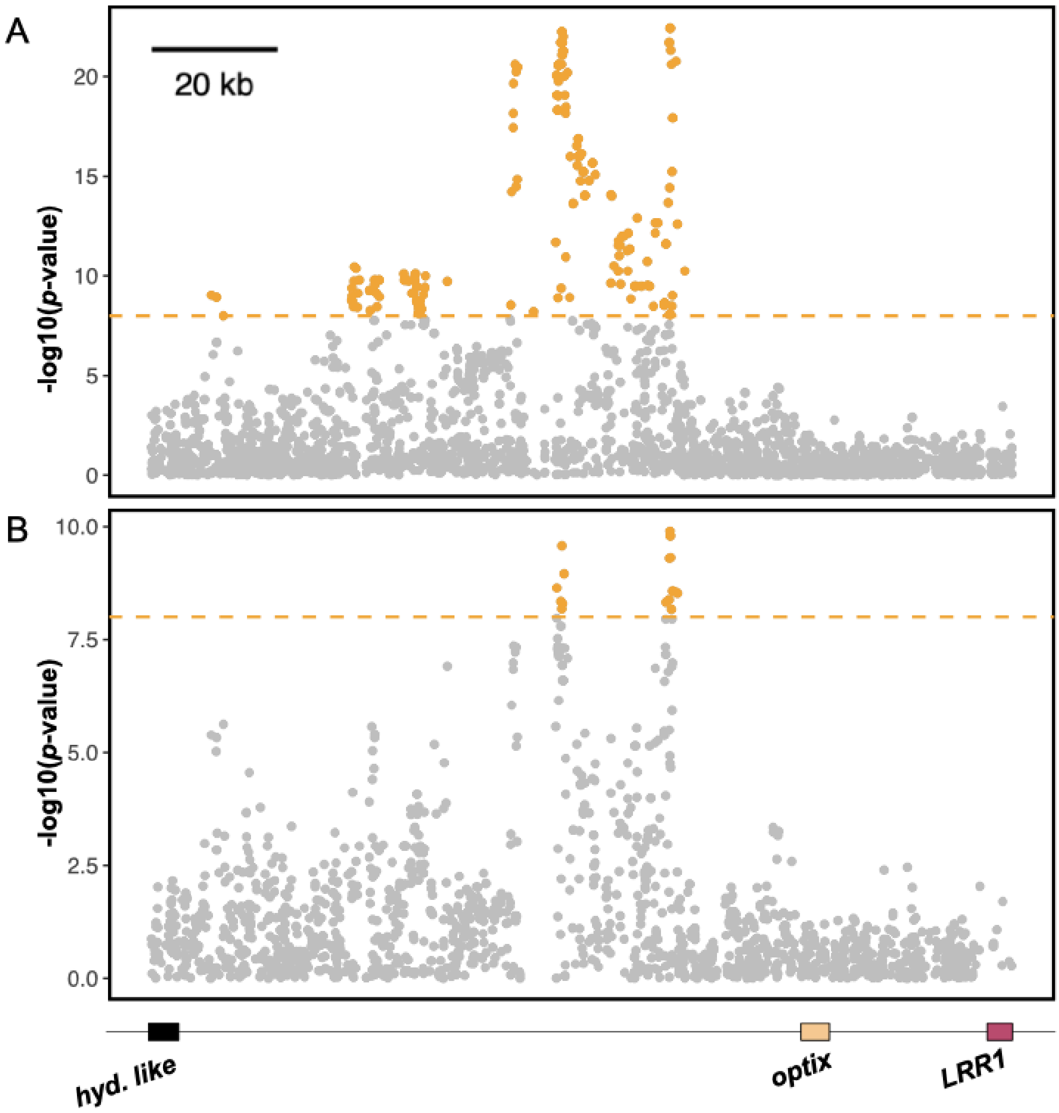


**Extended Data Fig. 38 | *Mechanitis messenoides* GWA around *optix* for hindwing black/orange using categorical grouping vs Patternize scoring for phenotyping**. (A) GWA using phenotype values based on manual, categorical grouping. Individuals were classed as either 0 (semi-melanised, “tiger-patterned” hindwing) or 1 (fully melanic, black hindwing). (B) GWA using phenotype values generated through a quantitative, colour-pattern analysis based approach using Patternize[53]. SNPs above the Bonferroni-corrected significance threshold (dashed orange line) are coloured orange.
